# Minor intron splicing efficiency increases with the development of lethal prostate cancer

**DOI:** 10.1101/2021.12.09.471104

**Authors:** Anke Augspach, Kyle D. Drake, Luca Roma, Ellen Qian, Se Ri Lee, Declan Clarke, Sushant Kumar, Muriel Jaquet, John Gallon, Marco Bolis, Joanna Triscott, José A. Galván, Yu Chen, George Thalmann, Marianna Kruithof-de Julio, Jean-Philippe P. Theurillat, Stefan Wuchty, Mark Gerstein, Salvatore Piscuoglio, Rahul N. Kanadia, Mark A. Rubin

## Abstract

Here we explored the role of minor spliceosome (MiS) function and minor intron-containing gene (MIG) expression in prostate cancer (PCa). We show MIGs are enriched as direct interactors of cancer-causing genes and their expression discriminates PCa progression. Increased expression of MiS U6atac snRNA, including others, and 6x more efficient minor intron splicing was observed in castration-resistant PCa (CRPC) versus primary PCa. Notably, androgen receptor signalling influenced MiS activity. Inhibition of MiS through siU6atac in PCa caused minor intron mis-splicing and aberrant expression of MIG transcripts and encoded proteins, which enriched for MAPK activity, DNA repair and cell cycle. Single cell-RNAseq confirmed cell cycle defects and lineage dependency on the MiS from primary to CRPC and neuroendocrine PCa. siU6atac was ∼50% more efficient in lowering tumor burden of CRPC cells and organoids versus current state-of-the-art combination therapy. In all, MiS is a strong therapeutic target for lethal PCa and potentially other cancers.

**Graphical Abstract:** U6atac expression, MiS activity, and minor intron splicing correlate with PCa therapy resistance and PCa progression to CRPC-adeno and transdifferentiation to CRPC-NE. One major MiS regulator during that process is the AR-axis, which is re-activated during CRPC-adeno and blocked in CRPC-NE. Molecularly, an increase in MiS dependent splicing promotes changes of transcriptome and proteome. This results in cell cycle activation, increased MAPK signalling and increased DNA repair. U6atac mediated MiS inhibition renders MiS splicing error-prone through increased intron retention and alternative splicing events, which results in cell cycle block and decreased MAPK signalling and DNA repair. MiS inhibition blocks all stages of PCa. Figure created with BioRender.com.

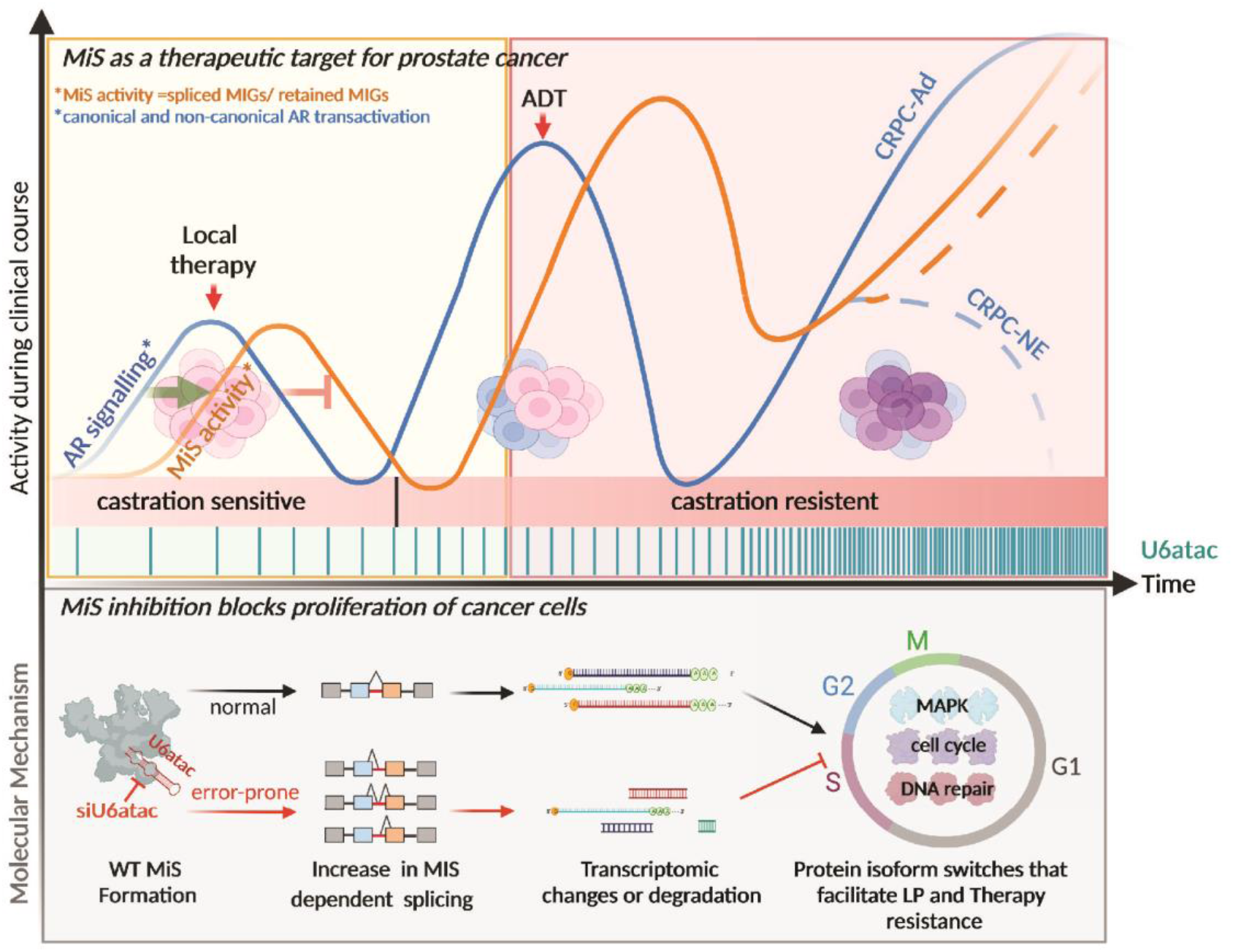

## Introduction

Cancer evolves continuously by modifying its transcriptome as it evades therapeutic interventions. Splicing, which is routinely leveraged, results in altered isoforms of key cancer genes involved in disease progression^1^. Currently, little is known about the minor spliceosome (MiS) in cancer development. Here, we explore prostate cancer (PCa) as an exemplar cancer to understand the role of MiS in disease progression.

Androgen deprivation therapy (ADT) is used to treat advanced PCa. For ADT-resistant PCa, potent second-generation androgen receptor signalling inhibitors (ARSi) such as enzalutamide and abiraterone are used. While initially effective, intrinsic or acquired resistance to ARSi in the form of castration-resistant prostate cancer (CRPC) eventually develops in this ultimately lethal disease. In CRPC, resistance to ARSi is conferred by intra-tumoral heterogeneity driven by re-activation of the AR axis, including gene body and enhancer amplifications, mutations, upregulation of AR co-activators and ligand-independent AR activity^2, 3^. Another increasingly recognized mechanism in the context of prolonged ARSi treatment is the transdifferentiation (also known as lineage plasticity) from adenocarcinoma (CRPC-adeno) to neuroendocrine PCa (CRPC-NE), which is extremely lethal and AR-indifferent^4^. Generally, CRPC-adeno and CRPC-NE are defined by expression/absence of characteristic markers such as *AR* and *KLK3* (CRPC-adeno) or *SYP* and *CHGA* (CRPC-NE), that can be quantified using AR- or NEPC-scores^5^.

While CRPC-adeno and CRPC-NE share similar genomic landscapes, they have dramatically distinct transcriptomes^6^, suggesting non-coding RNA events and RNA splicing as potential mechanisms of PCa transdifferentiation and progression. Indeed, multiple studies propose that alternative splicing of the AR transcript plays a key role in therapy resistance of CRPC-adeno^7^, whereas the splicing factor SRRM4 has been identified as a crucial driver of CRPC-NE transdifferentiation^8^. In general, non-canonical splicing has been extensively linked to prostate tumorigenesis, and many PCa relevant genes display isoform switching during cancer development and progression. However, little is known about the pathways controlling non-canonical splicing and a unified hypothesis explaining the molecular origin of those isoforms is still lacking^7, 9, 10^. Thus, understanding the mechanisms underlying aberrant splicing in PCa is essential both for predicting tumor progression and for discovering key regulators. The role of canonical splicing in cancer has been studied extensively, at present an understanding of the interplay between minor intron splicing and cancer is lacking.

Minor introns (<0.5%), spliced by the MiS, are usually found in genes with mostly major introns. These minor intron-containing genes (MIGs) execute diverse functions in disparate molecular pathways yet are highly enriched in genes that are essential for survival^11^. The essentiality of MIGs is reflected in early embryonic lethality when MiS is inhibited in mice, zebrafish, and Drosophila^12–14^. Regarding cancer, the dysregulation of minor intron splicing has been linked to the Peutz-Jegher’s syndrome and myelodysplastic syndrome, which frequently proceed to gastrointestinal cancer and acute myeloid leukemia (AML), respectively^9, 15–17^. There is further evidence that MiS components show a reliable association with an increased risk of scleroderma (U11/U12-65K protein), AML (U11-59K protein) and familial PCa (U11 snRNA)^18^. In fact, microRNA profiling studies of high-risk Finnish PCa families identified altered U11 snRNA expression as a risk factor to develop PCa^9, 19^. Taken together, we hypothesized that the minor spliceosome and MIG expression play vital roles in cancer.

Here, we show that MiS function, which is regulated by the AR-axis, increases with (prostate) cancer disease stage and degree of differentiation. In fact, the MiS component U6atac snRNA might serve as an additional marker for cancer diagnostics. We show that siU6atac-mediated MiS inhibition is more effective at blocking PCa cell proliferation than the current state-of-the-art combination therapy, such as EZH2 inhibitor/enzalutamide. Finally, we show that other MiS components can also be targeted, and that MiS inhibition also blocks proliferation of other cancer cell types. In all, our work elucidates a novel pathway, the minor spliceosome, as point of entry for therapeutics against lethal PCa, a strategy that extends to other cancer types.

## Results

### MIGs play a crucial role in cancer onset and progression

Cancer onset is often linked to disruption of a specific oncogene, but it essentially relies on a concerted activity of master regulators to sustain itself^20^. Since MIGs are highly enriched in cell cycle regulation and survival, we explored whether MIGs are enriched in biological pathways exploited by cancer-causing genes. For this, we determined whether MIG-encoded proteins were enriched amongst proteins that interact with proteins encoded by cancer-causing genes in a network of 160,881 protein-protein interactions (PPI) between 15,366 human proteins as of the HINT database^21^. We hypothesized that MIGs are prime recipients of molecular information that cascade from cancer-causing genes through their protein-protein interactions. Considering 403 cancer-causing genes in the set of 15,366 interacting proteins as of the Cancer Genome Interpreter database (Supplementary Table 1)^22^, we computed the shortest distance *d* of all proteins encoded by cancer-causing genes, where *d* = 1 indicates a direct interaction. In each distance bin, we determined the enrichment of 542 MIG-encoded proteins (inset, **Fig. 1A**) to find significant enrichment of MIGs in the immediate network neighbourhood of cancer-causing genes (permutation test: *d* = 1,2; p<10E-5) (**Fig. 1A**). However, MIGs were significantly diluted at higher distances (d > 3, p<10E-5) (green box in inset, **Fig. 1A**). Next we focused on immediate interactors of the proteins encoded by 26 PCa-causing genes (inset **Fig. 1B**, Supplementary Table 1)^22^. We found a significant enrichment of MIG-encoded proteins as immediate interactors of these 26 PCa proteins (**Fig. 1B**). Since MIGs were significantly enriched at *d=1* from cancer-causing genes, we next identified 3 PPI subnetworks of interactions between MIG encoded proteins and the 26 PCa proteins (**Fig. 1C**). In fact, we found that MIG-encoded proteins formed a significantly large (permutation test, p<6.8×10E-4) tight-knit web of interactions between 17 PCa proteins and 72 MIG-encoded proteins, capturing more than 95% of proteins (**Fig. 1B, C**). Similarly, significant (permutation test, p < 3 x 10E-5) large tight-knit subnetworks were obtained for MIG-encoded proteins and proteins encoded by all 403 cancer-causing genes (**Fig. 1A, red**). Such findings suggest that proteins encoded by cancer-causing genes in general and PCa causing-genes in particular tend to significantly interact with MIG-encoded proteins in the underlying human interaction network.

**Figure 1.**
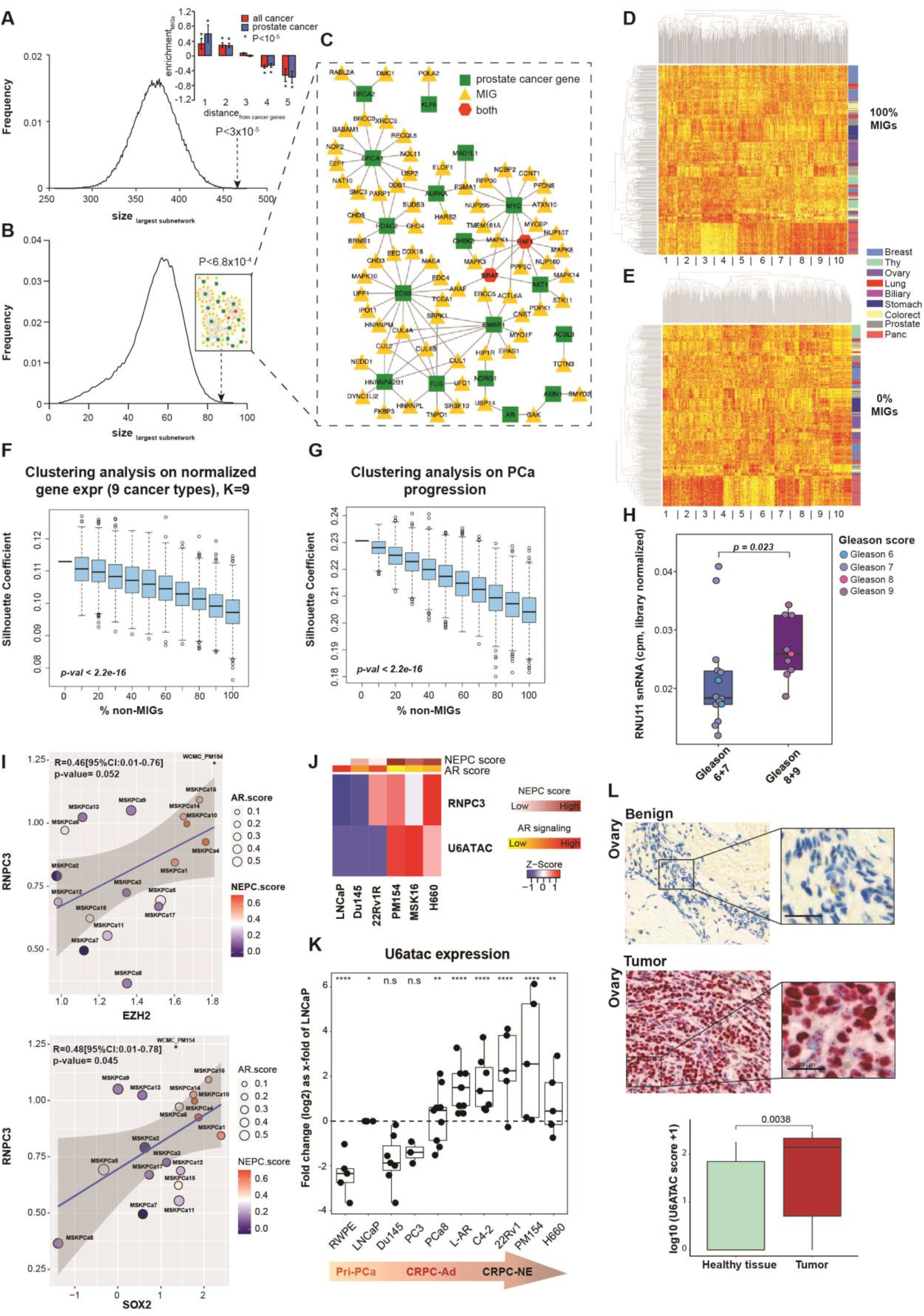
MIG expression patterns and MiS component expression correlate with PCa progression. (A) Enrichment of MIGs (542) amongst proteins interacting at a distance (d) =1, d=2, d=3, d=4 and d=5 with proteins encoded by 403 cancer and 26 prostate cancer genes in a human protein-protein interaction network. Enrichment values show the log2 fold change of the presence of MIGs in distance bins, compared to randomly sampled gene sets of equal size. Error bars indicate standard deviations. Asterisk indicates P < 10E-5. (B) Largest connected subnetwork of all interactions between prostate cancer genes and MIGs captured 87 genes (inset). Curve indicates the distribution of network sizes when we randomly sampled sets of non-MIGs of equal size (542). (C) Subnetwork of interactions between 74 MIGs and 19 prostate cancer genes (amplification of inset in 1B). (D) Heatmap gene expression data from different stages of prostate cancer progression. Gene expression data were taken from GTEx, TCGA, and SU2C; respectively, these datasets represent increasing stages of tumor progression (from healthy tissue in GTEx to advanced prostate tumors in SU2C). Clustering was performed on a gene set in which 100% of the genes are MIGs. (E) Heatmap gene expression data from different stages of prostate cancer progression. Gene expression data were taken from GTEx, TCGA, and SU2C; respectively, these datasets represent increasing stages of tumor progression (from healthy tissue in GTEx to advanced prostate tumors in SU2C). Clustering was performed on a gene set in which 0% of the genes are MIGs. (F) Relative performance of MIGs and non-MIGs with respect to clustering gene expression data from nine distinct cancer types from PCAWG (Biliary-, Breast-, Colorectal-, Lung-, Ovary-, Pancreas-, Stomach-, Thymus-, and Prostate-Adenocarcinoma). Each box corresponds to a 1000-length simulation in which non-MIGs are randomly sampled from among all non-MIGs in the genome. This plot exhibits a negative relationship between the relative abundance of non-MIGs and the quality of clustering, as quantified using the Silhouette coefficient. The P-value is based on a two-sided t-test in which each sample value is taken to be the difference between the Silhouette coefficient of one of the 1000 100% non-MIG samples and the corresponding Silhouette coefficient when only MIGs are used for clustering (i.e., the Silhouette coefficient corresponding to 0% non-MIGs). (G) Relative performance of MIGs and non-MIGs with respect to clustering gene expression data from different stages of prostate cancer progression. Gene expression data were taken from GTEx, TCGA, and SU2C; respectively, these datasets represent increasing stages of tumor progression (from healthy tissue in GTEx to advanced Prostate tumors in SU2C). Clustering was performed using an n_cluster value of K=3. Each box in this plot corresponds to a 1000-length simulation in which non-MIGs are randomly sampled from among all non-MIGs in the genome. This plot exhibits a negative relationship between the relative abundance of non-MIGs and the quality of clustering, as quantified using the Silhouette coefficient. The P-value is based on a two-sided t-test in which each sample value is taken to be the difference between the Silhouette coefficient of one of the 1000 100% non-MIG samples and the corresponding Silhouette coefficient when only MIGs are used for clustering (i.e., the Silhouette coefficient corresponding to 0% non-MIGs). (H) Boxplots representing the distribution of *RNU11* expression values (counts per million (CPM)) across primary tumor samples where *RNU11* was detected. Samples (n = 23) were grouped according to low (6+7) or high (8+9) Gleason score (n = 23, Wilcoxon Test, p-value indicated). (I) Pearson correlation analysis between RNPC3 and EZH2 (top) and RNPC3 and SOX2 (bottom). Organoids are color-coded according to their transcriptomic NEPC score, whereby a score >0.4 indicates a CRPC-NE phenotype while a score<0.4 indicates a CRPC-Adeno phenotype (Beltran et al., PMID 26855148). Organoids are size-coded according to their AR-score. (J) Heatmap showing RNA-seq expression (FPKM) of prostate cancer cell lines, ordered by increasing NEPC and decreasing AR score. The NEPC and AR score were calculated based on FPKM values of a set of 70 and 27 genes to estimate the likelihood of a test sample to be CRPC-NE or CRPC-adeno, respectively^5^ (K) U6atac snRNA expression as x-fold of siScrambled normalized to the mean of GAPDH and ACTB gene transcription in different PCa cell lines (RWPE n=5, LNCaP n=7, DU145 n= 7, PC3 n=3, PCc8 n=9, L-AR n=10, C4-2 n=7, 22Rv1 n=5, PM154 n=5, H660=5). Data are represented using box and whisker plots. Boxplots display values of minimum, first quartile, median, third quartile, and maximum (two-sided Mann-Whitney test, ns; not significant, p > 0.05, ∗p < 0.05, ∗∗p < 0.01, ∗∗∗p < 0.001). Each data point represents the x-fold of LNCaP cells from a single experiment. Experiments were performed in triplicates. (L) Representative images of the validation of RNU6atac BaseScope probes in human ovary benign and tumor tissue. Scale bars represent 50μm. Lower panel shows the U6atac score of the TMA analysis (Wilcoxon Test, p=0.0038).

That MIGs interact directly with cancer-causing proteins led us to explore the extent to which MIGs exhibit a greater degree of differential expression between distinct cancer types (relative to non-MIGs). To address this question, we first used normalized MIG expression data to perform hierarchical sample clustering (HSC) of 9 different solid cancer types, and then evaluated the extent to which this MIG expression successfully clusters samples by cancer type. We first evaluated the quality of the resultant clustering by visualizing the clustered data with a dendrogram and its associated heatmap, wherein structure can readily be seen in the case for which only MIGs were used in clustering the samples (**Fig. 1D** and Supplementary Fig. S1.1 and S1.2). To evaluate whether MIGs more successfully cluster the data relative to non-MIGs, we performed the same type of visual analysis for datasets comprising different relative abundances of MIGs and non-MIGs (Supplementary Fig. S1.1). Specifically, this controlled approach entailed progressively polluting the pool of MIGs with increasing samples of non-MIGs, such that 0%, 10%, 20%, 30%, … 100% of all genes in a given gene set were non-MIGs. In adopting this approach, we found that these progressive increases in the relative abundances of non-MIGs (within this set of genes used for clustering) gave rise to progressively deteriorating quality in data clustering. This can be seen visually by observing the deteriorating structure in the heatmaps associated with these various relative abundances of non-MIG (**Fig. 1E**; Supplementary Figs. S1.1 and S1.2).

We then took a more objective, quantitative approach to measure this deterioration in clustering (i.e., visual structure within these heatmaps). To quantitatively evaluate the extent to which a given expression dataset may be clustered, we used the Silhouette coefficient, wherein higher coefficient values designate more clearly-defined and correct clustering^23–26^. We employed a simulation (data re-sampling-based scheme) to generate 1000 gene sets for each fraction of non-MIGs (from 0% to 100% MIGs, in increments of 10%), and then calculated the Silhouette coefficient associated with each resampled gene set. This quantitative approach recapitulated what we had observed visually in the heatmaps: the Silhouette coefficients associated with clustering expression data in gene sets with higher fractions of non-MIGs resulted in less effective clustering for these 9 different cancer types (**Fig. 1F** and Supplementary Fig. S1.1 and S1.2; two-sided t-test p<2.2E-16). That MIGs enable better clustering performance than do non-MIGs suggests that MIG expression is more informative for defining differences between these 9 cancer types.

We next evaluated whether MIG expression may likewise exhibit greater differential gene expression across different stages of PCa progression (relative to non-MIGs). This was carried out for the transdifferentiation analysis across the distinct transcriptomes, illustrated by principal component analysis (Supplementary Fig. S1.3). Specifically, to obtain data representing different stages of PCa progression, we used prostate samples from GTEx (normal tissues), TCGA (primary PCa samples), and SU2C (CRPC-adeno and CRPC-NE) datasets. As with our clustering analysis on 9 distinct cancer types detailed above, we observed that MIG expression resulted in better clustering performance on these distinct phases of prostate cancer progression (Supplementary Fig. S1.4), although the visual disparities between the heatmaps associated with 0% non-MIGs and 100% non-MIGs are not as pronounced as what we had observed in the context of our pan-cancer analysis. The same simulation-based scheme in which Silhouette coefficients were measured in our pan-cancer analysis was similarly applied in the context of these broad stages of cancer progression. Again, we observed that MIG expression results in greater clustering performance than non-MIG expression (**Fig. 1G**; two-sided t-test p<2.2E-16). Together these findings show that transcription of MIGs is closely linked to cancer onset and progression.

### MiS component expression correlates with PCa progression

The upregulation of MIGs with PCa progression and transdifferentiation would necessitate increased MiS activity, which in turn is regulated by the levels of MiS components. Therefore, we queried the expression kinetics of U11 snRNA (*RNU11)*, a crucial MiS component in PCa (PRAD) TCGA data. We observed a significant (Wilcoxon Test, p=0.023) association between high-grade (Gleason score 8 and 9) PCa patients and high *RNU11* expression **(Fig. 1H** and Supplementary Table 2). Another unique MiS component is the protein RNPC3, which has been linked to cancer-associated scleroderma^27, 28^. Therefore, we queried RNA-seq data from 18 PCa organoids (CRPC-adeno and CRPC-NE), which revealed that expression of *RNPC3* showed a tendency towards positive correlation with the pluripotent stem cell and early differentiation markers *SOX2* and epigenetic regulator *EZH2*, respectively **(Fig. 1I).** We observed *RNPC3* expression paralleled PCa disease progression with the lowest expression in benign and hormone-sensitive PCa cells (LNCaP), intermediate expression in aggressive CRPC-adeno cells (22Rv1) and the highest expression in CRPC-NE cells (H660) and patient-derived organoid lines (WCM154 and Msk16) (**Fig. 1J**). The same was observed for U6atac snRNA which is regulated by high turnover rate (and is therefore a crucial regulatory node of MiS activity). High *RNU6atac* and *RNPC3* expression further correlated with high NEPC and low AR scores, which are indicators for poor PCa prognosis due to CRPC-NE lineage plasticity **(Fig. 1J).** Moreover, low U6atac levels inherently throttle MiS activity, so its increase in expression is especially informative. Indeed, quantitative-RT-PCR data confirmed that MiS component expression correlates with PCa progression: expression of the four MiS snRNAs – U6atac, U4atac, U11 and U12 – was higher in CRPC-adeno and CRPC-NE cell lines relative to hormone-responsive LNCaP and RWPE cells or to AR-low Du145 and PC3 cells (**Figs. 1K** and Supplementary Fig.1.5).

To test the prediction that U6atac expression is elevated with cancer onset and progression, we developed an *in-situ* probe specifically against U6atac (Supplementary Fig. S1.6). We used this probe to survey different cancer types, and found significantly higher expression of U6atac in cancer tissue compared to matching benign samples (Wilcox test, p=0.0038). This finding demonstrates a strong specificity of U6atac expression for highly proliferative tissues (pan-cancer Tissue Microarray (TMA)) (**Figs. 1L** and Supplementary Fig. S1.7). Consistently, thyroid and colon cancer showed a higher U6atac intensity score in cancer compared to the corresponding benign tissues (Supplementary Fig. S1.7). Together our data underscore that expression of both MiS components and MIGs show a reliable association with cancer onset and progression.

### MiS activity correlates with PCa progression

Consistent with the increased expression of U6atac in more aggressive PCa cell lines (**Fig. 1J** and **1K**), we also observed significantly increased U6atac expression in primary and metastatic PCa patient samples compared to benign prostate (Wilcox test, p=0.0004). More importantly, we discovered that metastatic PCa had significantly elevated U6atac expression compared to primary PCa (Wilcox test, p=0.013) (**Figs. 2A** and S2.1). Thus, we posited that MiS activity should increase with PCa progression. To test this, we used minor and major splicing reporter plasmids (kindly provided by the Dreyfuss Lab^29^) in different PCa cell lines reflecting the range of PCa progression and transdifferentiation. We found a trend towards more efficient minor intron splicing in therapy-resistant C4-2 and 22Rv1 cells compared to the hormone-sensitive LNCaP cells (**Fig. 2B, upper panel**) or AR-low DU145 and PC3 cells. We also observed a six-fold increase in minor intron splicing in the CRPC-NE cell and patient-derived organoid lines (PM-154^30^, PM1262^30^, MSK16^31^ and NCI-H660^32^) (**Fig. 2B, upper panel**). In contrast, major intron splicing was unaffected in all cell and organoid lines tested (**Fig. 2B, lower panel**) indicating that, unlike the major spliceosome, minor spliceosome activity dynamically increases with PCa progression. This led us to explore the molecular mechanisms underlying the regulation of MiS activity in PCa.

**Figure 2.**
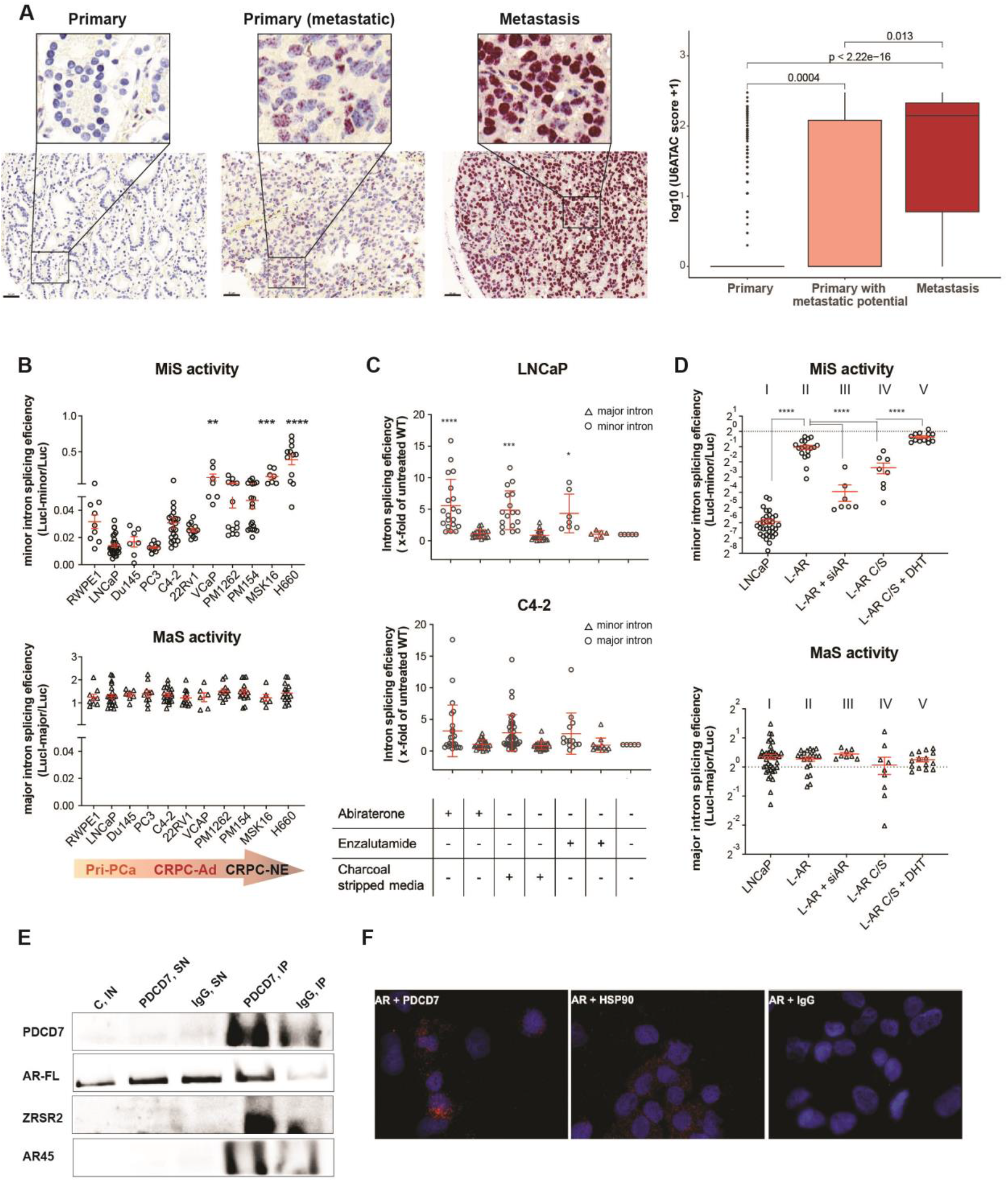
MiS activity is regulated by AR and is elevated in prostate cancer. (A) Representative images of the validation of RNU6atac BaseScope probes in human primary, primary with metastatic potential and metastatic PCa tissue. Scale bars represent 50μm. Quantification of the U6atac score of the whole TMA analysis shown on the right-hand side. Comparisons between groups were performed with Wilcox test (primary vs. primary (metastatic: p=0.004, primary vs. metastasis: p<2.22e-16, primary (metastatic) vs. metastasis: p=0.013). (B) Normalized luminescence values of minor/major spliceosome luc-reporter plasmids in different PCa cell lines of different PCa subtypes (RWPE n=9, LNCaP n=30, DU145 n= 7, PC3 n=10, C4-2 n=22, 22Rv1 n=13, VCAP n=7, PM1262 n=13, PM154 n=20, MSK16 n=7, H660=12, mean ± SEM, one-way Anova; ns p > 0.05, ∗p < 0.05, ∗∗p < 0.01, ∗∗∗p < 0.001). Each data point represents a single experiment; experiments were performed in triplicates. Abbreviations: Pri-PCa=Primary Prostate Cancer, CRPC-Ad= Castration Resistant Adeno Prostate Cancer, CRPC-NE = Castration Resistant Neuroendocrine Prostate Cancer (C) Normalized luminescence values of minor/major spliceosome luc-reporter plasmids in LNCaP and C4-2 PCa cells subjected to long-term treatment with abiraterone, enzalutamide and charcoal stripped (c/s) media (LNCaP: Abiraterone n=19, c/s n=17, Enzalutamide n=7, C4-2: Abiraterone n=24, c/s n= 29, Enzalutamide n=12, mean ± SEM, one-way Anova; ns p > 0.05, ∗p < 0.05, ∗∗p < 0.01, ∗∗∗p < 0.001). Each data point represents a single experiment; experiments were performed in triplicates. (D) Normalized luminescence values of minor/major spliceosome luc-reporter plasmids in LNCaP and L-AR cells treated with siAR, c/s media and c/s media + DHT (10uM) (LNCaP n=34, L-AR n=19, L-AR + siAR n=7, L-AR c/s n=8, L-AR c/s + DHT n=13, mean ± SEM, ordinary one-way Anova; ns p > 0.05, ∗p < 0.05, ∗∗p < 0.01, ∗∗∗p < 0.001). Each data point represents a single experiment, that were performed in triplicates. Abbreviations: c/s= charcoal stripped media, DHT= Dihydrotestosterone. (E) Co-IP of AR followed by immunoblotting for PDCD7, ZRSR2, GAPDH in cytosolic extracts from L-AR cells. Abbreviations: IN= Input, SN= supernatant, IP = Immunoprecipitation. Linked PLA secondary probes [mouse, M(+) and rabbit, R(-)]. The experiment was repeated 3 times. (F) Representative 40x confocal photomicrographs of PLA signal from AR and PDCD7 or HSP90 (positive control) heteromeric formation (red puncta) and associated negative IgG rabbit control. PLA was performed using anti-PDCD7 antibody (rabbit monoclonal) and androgen receptor antibody (mouse monoclonal) primary antibody and oligonucleotide-linked PLA secondary probes [mouse, M(+) and rabbit, R(-)]. Scale bars represent 25 μm. The experiment was repeated 3 times.

AR signaling plays a critical role in PCa progression, and is often the apex of oncogenic pathways. Therefore, we hypothesized that MiS activity across PCa progression might be linked to AR signaling. To simulate stress response and re-activation of AR signalling in PCa, we mimicked therapy resistance mechanisms, and subjected PCa cells to long term androgen deprivation therapy (ADT) and ARSi using charcoal-stripped (C/S) media, abiraterone and enzalutamide. We observed a significant increase in MiS activity in cells exposed to ADT/ARSi, whereas the treatment had only limited effects on major splicing (one-way ANOVA; ∗p =0.0117, ∗∗∗p and ∗∗∗∗p < 0.001) (**Fig. 2C)**. A similar association was observed in PC3 cells and their more highly differentiated subline PC3Pro4 as well as in hormone-sensitive LNCaP cells and a subline which overexpresses AR (L-AR and/or is ENZ-resistant (L-rENZ) (Supplementary Fig. S2.2A and B). PC3Pro4, L-AR and L-rENZ displayed significantly increased MiS activity but unchanged major spliceosome activity (Supplementary Fig. S2.2B) (one-way ANOVA; ∗∗∗p and ∗∗∗∗p < 0.001). This increase in MiS activity in those lines was further confirmed by the minor-intron containing p120 and hSCN4A minigene constructs. We observed that L-AR, L-rENZ- and PC3Pro4-cells, compared to their native LNCaP and PC3 lines, displayed a higher splicing efficiency in both reporter assays (Supplementary Fig. S2.2C). Lastly, both L-rENZ and PC3Pro4 cells expressed higher MiS snRNA levels (Supplementary Fig. S2.2D) compared to their respective WT lines.

Next, we explored the relationship of AR signaling to minor intron splicing. Here, we used luciferase reporter as the readout of minor intron splicing in therapy-sensitive LNCaP cells. The overexpression of AR led to a significant increase in MiS activity (**Fig. 2D II**) (one-way Anova; ∗∗∗∗p < 0.001), which was not observed with the major spliceosome reporter (**Fig. 2D II, lower panel**). MiS activity decreased upon siRNA against AR (**Fig. 2D III)**. AR KD was confirmed by qRT-PCR (Supplementary Fig. S2.3A). To further explore the regulation of MiS activity by AR signaling, we treated L-AR cells with hormone-depleted media to block AR activation, which was confirmed by a reduction in *KLK3* expression (Supplementary Fig. S2.3B). Again, we found that blocking AR activation resulted in decreased MiS activity **(Fig. 2D IV)**. Finally, addition of dihydrotestosterone (DHT) to the hormone-depleted media rescued MiS activity to levels observed in L-AR alone **(Fig. 2D V)**. Similar findings were observed with the p120 minor intron splicing minigene reporter (Supplementary Fig. S2.3C). In contrast, major spliceosome activity was not affected by AR modulation **(Fig. 2D, lower panel).**

The association between AR activation and MiS activity suggests the existence of an AR-MiS regulatory axis. To address this, we determined whether AR interacts with MiS proteins. Co-immunoprecipitations (Co-IPs) with an antibody directed against the AR revealed an interaction with PDCD7, one of the seven proteins unique to the MiS complex, and *vice versa,* implying a direct regulation of the MiS by AR through PDCD7 (**Fig. 2E** and Supplementary Fig. S2.3D). AR45, an AR splice variant known to heterodimerize with the full-length version, and ZRSR2, which is known to interact with the MiS^33, 34^ were used as positive controls. Further support was obtained by proximity ligation assays demonstrating an *in-situ* interaction between AR and PDCD7 as well as with the AR interactor HSP90 **(Fig. 2F** and Supplementary Fig. S2.4). Taken together, this data suggest that MiS activity is regulated by AR signalling, it can be stimulated by ADT, and that it increases with PCa disease progression to CRPC-adeno and CRPC-NE.

### MiS inhibition in prostate cancer results in aberrant minor intron splicing

Based on our finding that the MiS plays a crucial role in PCa progression, we next inhibited the MiS in PCa by targeting the U6atac snRNA, which normally exhibits higher rates of turnover but is detected at higher levels in advanced cancer stages. We used siRNA against U6atac in four PCa cell lines: LNCaP (primary PCa, therapy-sensitive), C4-2 (CRPC-adeno, therapy-resistant), and 22RV1 (CRPC-adeno, therapy-resistant) cell lines and a patient-derived organoid, PM154 (CRPC-NE, therapy-resistant). First, we chose C4-2 cells to establish the kinetics of U6atac downregulation (**Fig. 3A**). Successful downregulation (∼7-fold, unpaired t-test: p< 0.0001) of U6atac snRNA in C4-2 cells upon treatment with siU6atac was confirmed by qRT-PCR, in line with an increase in U6atac expression (∼6-fold, t-test: p<0.0001) in C4-2 cells overexpressing U6atac **(Fig. 3A**). We then explored the kinetics of MiS inhibition on minor intron splicing by analysing *COA3*, which has a single intron that is a minor intron. The resulting minor spliceosome inhibition was reflected in a progressive increase of unspliced *COA3 natural* minor intron reporter transcripts across time, with the highest change observed at 96 hours post transfection, the latest time point investigated (**Fig. 3B**). Similar results were obtained in other cell lines (Supplementary Fig. S.3.1A). This finding was further confirmed by decreased splicing of three minor intron splicing minigene reporters p120, hSCN8A and hSCN4A (Supplementary Fig. S3.1B) in siU6atac-treated cells compared to siScrambled controls. Additionally, failure to splice minor introns was observed by the decrease in luminescence signal with a luc-minor splicing reporter^35^ (Supplementary Fig. S3.1C).

**Figure 3.**
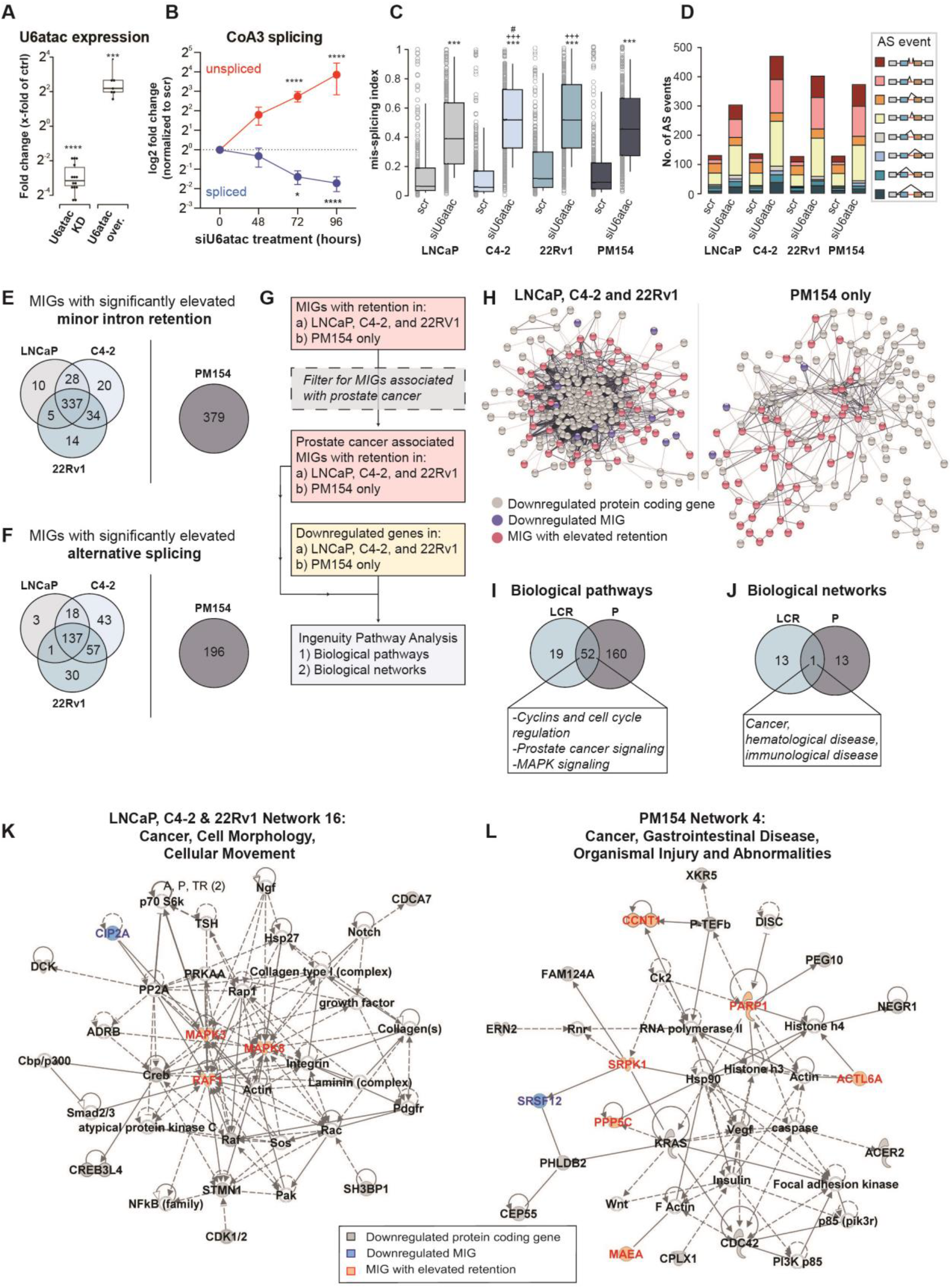
Inhibition of the MiS through U6atac siRNA effectively alters the PCa transcriptome. (A) U6atac snRNA expression as x-fold of siScrambled (control) normalized to the mean of GAPDH and ACTB gene transcription in C4-2 cells (U6atac KD n=14, U6atac over. n=7) Data are represented using box and whisker plots that display values of minimum, first quartile, median, third quartile, and maximum. Statistical analysis was evaluated using two-sided unpaired t-test, ns p > 0.05, ∗p < 0.05, ∗∗p < 0.01, ∗∗∗p < 0.001. Each data point represents a single experiment; experiments were performed in triplicates. (B) Expression of *CoA3* spliced and unspliced transcripts as x-fold of siScrambled normalized to the mean of GAPDH and ACTB gene transcription in C4-2 cells treated with siU6atac RNA for 48, 72 and 96 hours (n=3, two-way Anova; ns p > 0.05, ∗p < 0.05, ∗∗p < 0.01, ∗∗∗p < 0.001). Experiments were performed in triplicates. (C) Box-plot showing 10^th^-90^th^ percentile mis-splicing index for all minor introns that show retention (431) in LNCaP, C4-2, and 22Rv1 cell lines as well as PM154 organoid after 96h treatment with siScrambled (scr) or siU6atac. Significance determined by Kruskal-Wallis test followed by *post-hoc* Dunn’s test: asterisk (*) denotes significance compared to each siU6atac samples’ appropriate siScrambled control; plus sign (+) denotes significance compared to LNCaP siU6atac; hashtag (#) denotes significance compared to PM154 siU6atac. ∗p < 0.05, ∗∗p < 0.01, ∗∗∗p < 0.001. (D) Bar chart showing the distribution of alternative splicing (AS) events around (from two exons upstream to two exons downstream) all minor introns in LNCaP, C4-2, and 22Rv1 cell lines and the PM154 organoid after 96h treatment with siScrambled (scr) or siU6atac. Different colors depict different alternative splicing categories. (E) Venn-diagram (not to scale) depicting the overlap of MIGs with significantly elevated minor intron retention in the LNCaP, C4-2, and 22RV1 cell lines (left) and the PM154 organoid (right) after 96h siU6atac treatment compared to the appropriate 96h siScrambled control. (F) Venn-diagram (not to scale) depicting the overlap of MIGs with significantly elevated alternative splicing in the LNCaP, C4-2, and 22RV1 cell lines (left) and the PM154 organoid (right) after 96h siU6atac treatment compared to the appropriate 96h siScrambled control. (G) Pipeline for generating and analyzing interaction networks between shared (i) MIGs with significantly elevated minor intron retention and (ii) significantly downregulated protein coding genes in all three cell lines (a) or in the PM154 organoid (b). MIGs associated with prostate cancer are shown in Fig. 1C. (H) STRING network showing association of shared (i) prostate cancer-associated MIGs with elevated minor intron retention (red) and (ii) downregulated protein coding genes (grey; downregulated MIGs in blue) in all three cell lines LNCaP, C4-2, and 22RV1 (left) and PM154 organoid (right). (I) Venn-diagram (not to scale) showing the overlap of Ingenuity Pathway Analysis (IPA)-generated biological pathways for the LNCaP, C4-2, and 22RV1 (LCR) gene list (**H**; left) and PM154 (P) gene list (**H**; right). (J) Venn-diagram (not to scale) showing the overlap of Ingenuity Pathway Analysis (IPA)-generated biological networks for the LNCaP, C4-2, and 22RV1 (LCR) gene list (**H**; left) and PM154 (P) gene list (**H**; right). (K) Representative example of LNCaP, C4-2, and 22RV1-specific IPA-generated biological network. (L) Representative example PM154-specific IPA-generated biological network.

To capture a comprehensive effect of MiS inhibition, we performed ribo-depleted total RNAseq on the four cell types treated with siU6atac and siScrambled for 96 hours. Equivalent levels of U6atac KD were observed in all 4 cell lines (Supplementary Fig. S3.1D). We first determined the overall level of minor intron retention by quantifying a mis-splicing index for each cell line. Comparison of the median mis-splicing index for each sample revealed significantly elevated minor intron retention in siU6atac-treated LNCaP, C4-2, 22RV1 cells, and PM154 compared their respective scrambled siRNA control (**Fig. 3C** and Supplementary Table 3) (Kruskal-Wallis test followed by *post-hoc* Dunn’s test, ∗p < 0.05, ∗∗p < 0.01, ∗∗∗p < 0.001, respectively). In agreement with our previous report of cell type specific effects of MiS activity (**Fig. 1 and 2**), we found that the levels of minor intron retention in siU6atac-treated C4-2 cells was significantly greater than both siU6atac-treated PM154 and LNCaP cells (Kruskal-Wallis test and *post-hoc* Dunn’s test: C4-2 vs LNCaP p<0.0001; C4-2 vs PM154 p = 0.0296). Similarly, the level of minor intron retention in siU6atac-treated 22Rv1 cells was also significantly greater than siU6atac-treated LNCaP cells (Kruskal-Wallis test and *post-hoc* Dunn’s test: p=0.0001). Since MiS inhibition is also predicted to trigger alternative splicing (AS) across minor introns (in addition to elevated minor intron retention), we next quantified the number of AS-events detected in all eight samples. Indeed, we found a robust increase in the number of AS-events occurring around minor introns in all four siU6atac-treated samples, with the most abundant change occurring for cryptic splice site usage (**Fig. 3D**). Regarding both minor intron retention and AS around minor introns, the largest effect upon siU6atac treatment was observed in C4-2 cells, followed by 22RV1 cells, then PM154, and lastly LNCaP cells (**Fig. 3D** and Supplementary Table 3).

Next, we explored whether there was a cell-type specific effect of MiS inhibition by performing an intersection analysis of the MIGs with significantly elevated minor intron retention or AS. Here, we separated the three cell lines from the organoid due to their different origins, culture conditions, genomic architectures and PCa phenotypes (CRPC-NE), which was also reflected in principal component analysis (Supplementary Fig. S3.2). Of the 380, 419, and 390 MIGs with significantly elevated minor intron retention in LNCaP, C4-2, and 22RV1 cells, respectively, we found that 337 MIGs were common to all (**Fig. 3E**, Supplementary Table 3). The PM154 organoid was found to have 379 MIGs with elevated minor intron retention, of which 303 MIGs were shared with the three cell lines. GO-term enrichment analysis of each set of MIGs with elevated minor intron retention enriched for basic molecular pathways such as RNA and DNA processing, vesicle transport and mRNA splicing (Supplementary Table 4). Similarly, of the 159, 255, and 225 MIGs with significantly elevated AS events in LNCaP, C4-2, and 22RV1 cells, respectively, and 137 MIGs were common to all (**Fig. 3F**, Supplementary Table 3). In PM154 cells, we detected 196 MIGs with significantly elevated AS-events (student’s two-tailed T-test, p<0.05), of which 111 were shared with the three cell lines (Supplementary Table 3).

The aberrant minor intron splicing in siU6atac-treated samples should impact the overall transcriptome of PCa, which we captured by differential gene expression analysis (Supplementary Fig. S3.3). We set a 1 transcript per million (TPM) threshold for gene expression and, using isoDE2, we found 68 genes that were significantly upregulated (log2FC ≥ 1, *p* ≤ 0.01) and 691 genes significantly downregulated (log2FC ≤ −1, *p* ≤ 0.01) in the siU6atac-treated LNCaP cells compared to siScrambled (Supplementary Fig. S3.3). Similarly, we found 154, 228, and 121 genes upregulated in 96h siU6atac-treated C4-2, 22Rv1, and PM154 organoids, respectively, with 787, 638, and 189 genes downregulated, respectively (Supplementary Fig. S3.3). Intersection analysis of the LNCaP, C4-2, and 22Rv1 cells revealed 11 upregulated protein-coding genes common to the three cell lines, and only 1 (*CNST*) was a MIG. In contrast, we found 268 downregulated protein coding genes common to the three cell lines, and 18 were MIGs (Supplementary Table 5). Using DAVID, we discovered that the shared downregulated genes enriched for many gene ontology (GO) Terms related to the cell cycle (Supplementary Table 4). Alternatively, the 189 downregulated genes in 96h siU6atac-treated PM154 cells enriched for a single GO Term – cell differentiation – which was unique to this cell line (Supplementary Table 6). Gene Set Enrichment Analysis (GSEA) based on hallmark gene-sets on the up- and downregulated genes confirmed that pro-proliferative (E2F targets, G2M checkpoint, mitotic spindle) and DNA repair pathways were reduced upon siU6atac mediated MiS inhibition and additionally revealed a reduction in prostate-specific pathways such as androgen response or Spermatogenesis (Supplementary Fig. S3.4).

Next, we sought to untangle the downstream molecular defect of aberrant minor intron splicing in conjunction with the transcriptomic changes captured by RNAseq. For this, we took the list of MIGs with elevated minor intron retention common to either the LNCaP, C4-2, and 22Rv1 cell lines **(Fig. 3G)** and selected MIGs we had identified as direct interactors of prostate cancer-causing genes **(Fig. 1B and C).** We focused here on minor intron retention alone, and not MIGs with AS, as the molecular event is singular (the minor intron is either retained or not retained, whereas multiple AS events can occur in conjunction) and thus simplifies the downstream molecular predictions. This yielded 55 MIGs for the LNCaP, C4-2, and 22RV1 cell lines (Supplementary Table 6). We hypothesized that minor intron retention in these MIGs would exert their influence on the transcriptome, which was reflected in the downregulated non-MIGs. Subsequently, we investigated whether the MIGs and non-MIGs formed a PPI network by performing STRING network analysis on the combined list of 55 MIGs with minor intron retention plus 268 downregulated genes in all three cell lines (**Fig. 3G**). Most of the downregulated genes formed a tight-knit central network that was surrounded by MIGs with elevated minor intron retention (**Fig. 3H**). The same analysis for PM154 organoids showed a slightly more diffused network, though the MIGs with elevated retention were found to be clearly associated with an abundance of the downregulated genes (**Fig. 3H**). Subsequently, we used Ingenuity Pathway Analysis (IPA) to i) uncover the biological pathways underlying these networks, and ii) determine if the same pathways affected in the cell lines were also affected in the organoid (**Fig. 3I and J**). Of the 71 and 212 significantly enriched biological pathways for the cell lines and organoid, respectively, we found 52 were shared (Supplementary Table 7). The overlapping pathways included terms such as cyclins and cell cycle regulation, PCa signalling, and MAPK signalling (**Fig. 3I**). Notably, both the cell lines and the organoid enriched for numerous unique terms relevant to apoptosis and MAPK signaling (Supplementary Table 7). Finally, we sought to breakdown the large STRING network into functionally relevant sub-networks, and then determine if similar sub-networks were affected in the cell lines and the organoid. We found that only a single sub-network was shared in both conditions, where the three leading disease associations were cancer, haematological disease, and immunological disease (**Fig. 3J**). All remaining sub-networks for the cell lines and the organoid did not overlap. Importantly, the sub-networks found to be enriched in the three cell lines were often associated with cell cycle (e.g., G2/M DNA Damage Checkpoint Regulation and G1/S Checkpoint Regulation) and DDR (e.g., Role of BRCA1 in DNA Damage Response and DNA Double-Strand Break Repair by Homologous Recombination). In contrast the organoid sub-networks were frequently associated with oncogenic signaling pathways (e.g., PI3K/Akt signaling and PTEN signaling) and with actions executed by the cell cytoskeleton (e.g., morphology changes, EMT and actin cytoskeleton), which regulates cancer hallmarks such as invasion, metastatic spread, migratory ability and cell movement. MIGs with elevated minor intron retention were found to be involved in nearly all the sub-networks for both the cell lines and the organoid (**Fig. 3K and L**). Overall, siRNA-mediated downregulation of U6atac resulted in a robust minor intron splicing defect in a large subset of MIGs involved in cancer-relevant pathways.

### Impact of MiS inhibition on the prostate cancer proteome

To determine whether the transcriptional changes captured for each cell line extended to differential protein production, we performed LC-MS/MS on the same cell lines treated with siRNA **(Fig. 4** and Supplementary Fig. S4.1). With U6atac knock-down (log2FC > ±1, *p*≤0.05) we found 314 up- and 353 downregulated proteins in LNCaP cells; 390 up- and 753 downregulated proteins in C4-2 cells; 341 up- and 689 downregulated proteins in 22RV1 cells; and 197 up- and 455 downregulated proteins in PM154 cells (**Fig. 4A** and Supplementary Table 8). For all cells, we found that most differentially expressed MIG-encoded proteins were downregulated (**Fig. 4A**). Importantly, almost all MIGs with significantly elevated minor intron retention were found to be downregulated in the LC-MS/MS (**Fig. 4A**). For the three cell lines (LNCaP, C4-2, and 22Rv1), we found shared upregulation of only 8 proteins (**Fig. 4B**), whereas 45 proteins were downregulated in all three (**Fig. 4C**). While none of the common upregulated proteins were encoded by MIGs, 9 of the common downregulated proteins were found to be MIGs, including SLC12A7, CEP170, TCEA3, POLA2, XPO4, WDFY1, ORC3, C17orf75, and SRPK1. Besides CEP170 and TCEA3 proteins, the rest of the proteins had significantly elevated minor intron retention in their respective transcripts by RNAseq in all cell lines. More importantly, POLA2 and SRPK1 are MIG-encoded proteins that directly interact with the proteins encoded by PCa-causing genes identified in **Fig. 1C**.

**Figure 4.**
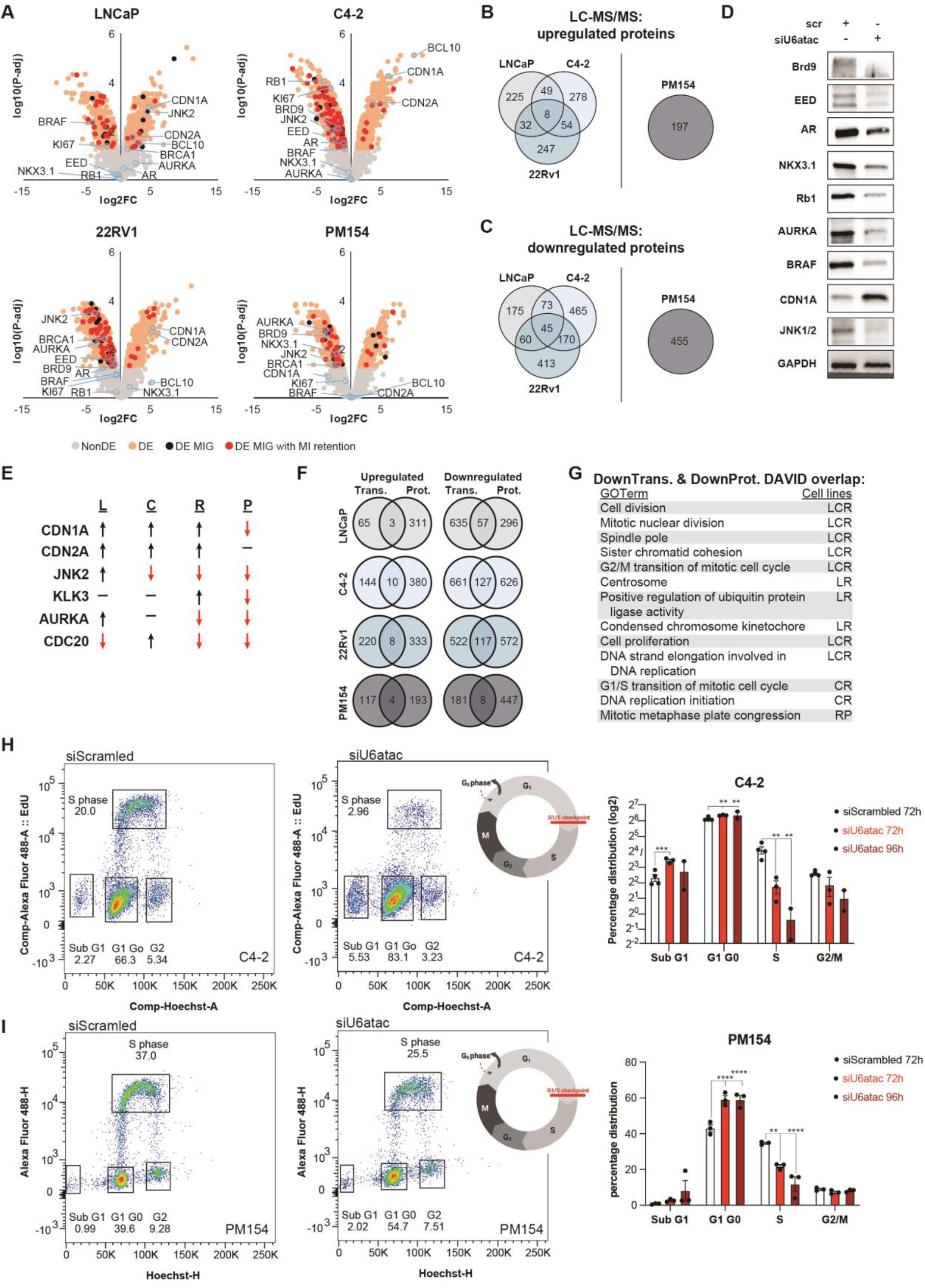
U6atac mediated MiS inhibition effectively alters the PCa proteome in a cell-type specific manner. (A) Volcano plot showing proteins most significantly increased (upper right) and decreased (upper left) in C4-2 cells treated with siU6atac (96h), as compared to the siScrambled control (pooled data from 3 co-IP replicates). The *x*-axis represents log2 fold change (FC) values, the *y*-axis represents −log10 of adjusted *p*-values. Grey dots represent non-differentially expressed (non-DE) proteins; orange dots represent differential expressed proteins (DE), black dots represent differentially expressed MIG-encoded proteins (DE MIG) and red dots represent differentially expressed MIG-encoded proteins that transcriptomically have significantly elevated minor intron retention. (B) Venn diagrams (not to scale) illustrating the overlap in proteins that are upregulated after U6atac KD in LNCaP, C4-2, 22Rv1 and PM154 cells, assessed by mass spectrometry analysis (C) Venn diagrams (not to scale) illustrating the overlap in proteins that are downregulated after U6atac KD in LNCaP, C4-2, 22Rv1 and Pm154 cells, assessed by mass spectrometry analysis (D) Immunoblot showing expression levels of selected proteins which are down- or upregulated in mass spectrometry analysis in C4-2 cell lines treated for 96 hours with siScrambled or siU6atac RNA. GAPDH was used as loading control. (E) Table summarizing differential expression of selected proteins after siU6atac treatment in LNCaP (L), C4-2 (C), 22Rv1 (R) and PM154 (P) cells. (F) Venn-diagrams (not to scale) illustrating the overlap of genes that show upregulation (left) or downregulation (right) transcriptionally (trans.) and by mass spec (prot.) in LNCaP, C4-2, 22Rv1, and PM154 cells. (G) Table summarizing the shared DAVID GO Terms for the genes that are downregulated both transcriptionally (DownTrans.) and by mass spec (DownProt.) for each cell line. L=LnCAP, C=C4-2, R=22Rv1, P=PM154. (H) Pseudocolor plots showing the gates for G1/0, S and G2 selection in C4-2 cells treated for 72h with siScrambled or siU6atac (left). Quantification of flow cytometry analysis at 72h and 96h (right), (n=3, mean ± SEM, ordinary two-way Anova; ns p > 0.05, ∗p < 0.05, ∗∗p < 0.01, ∗∗∗p < 0.001). (I) Psuedocolour plots showing the gates for G1/0, S and G2 selection in PM154 cells treated for 72h with siScrambled or siU6atac (left). Quantification of flow cytometry analysis at 72h and 96h (right), (n=3, mean ± SEM, ordinary two-way Anova; ns p > 0.05, ∗p < 0.05, ∗∗p < 0.01, ∗∗∗p < 0.001).

Comparing the proteomic data of the four analysed cell lines representing PCa disease progression revealed a cell type, and thus probably PCa subtype and context dependent MiS-dependent proteome, much like the RNAseq analysis. Each cell line expressed a unique set of up and downregulated proteins including MIGs and non-MIGs, **(Fig. 4B and C** and Supplementary Table 8) each in its own way important for cancer biology. Among others, the AR (non-MIG), Rb1 (non-MIG), the epigenetic regulator EED (MIG), a common target during PCa therapy, and the proliferation marker Ki67 (non-MIG) were strongly decreased upon siU6atac treatment in the CRPC-adeno line C4-2 (**Fig. 4D**). Similarly, the DNA damage response (DDR) proteins BRCA1 and AURKA (non-MIG) and JNK2 were decreased in therapy-resistant C4-2, 22Rv1 and PM154 lines but increased in therapy-responsive LNCaP cells (**Fig. 4E**). AURKA has been linked to lineage plasticity and neuroendocrine differentiation in PCa^36^. Consistent with previous results, we observed an increase in tumour suppressors such as the cell cycle regulators CDN1A (p21) (non-MIG) and CDN2A (non-MIG) in all three cell lines tested (**Fig. 4A**). In contrast, we found that these proteins were decreased in the siU6atac-treated neuroendocrine organoid PM154 (**Fig. 4E**). Finally, we also observed an increase in pro-apoptotic proteins, such as BCL10 (non-MIG), upon MiS inhibition (**Fig. 4A**). GSEA based on hallmark gene-sets on the up- and downregulated proteins (Supplementary Fig. S4.2) confirmed that pro-proliferative (E2F targets, G2M checkpoint, mitotic spindle) and DNA repair pathways were reduced, whereas apoptotic and stress sensing pathways (p53, unfolded protein response, UV response or hypoxia mediated oxidative stress) were increased upon siU6atac mediated MiS inhibition. Differential protein expression was validated by WB analysis (**Fig. 4D** and Supplementary Fig. S4.3).

We next explored concordance between the transcriptome and proteome data, which were generally discordant in each cell line (**Fig. 4F**). MIGs with elevated retention upon MiS inhibition have been reported to escape NMD^37^. For example, we found proteins such as EED, JNK1/2, BRAF and RAF1 to be decreased even though their expression by RNAseq remained unchanged, albeit with elevated minor intron retention and AS upon siU6atac treatment (**Fig. 4D**, Supplementary Table 6, Supplementary Fig. S3.3). We also found proteins such as the AR and Rb1 to be decreased only in the proteome, implying indirect regulation mechanisms executed through the MiS. Despite this, we identified 57, 127, 117, and 8 genes whose mRNA transcripts and encoded proteins were significantly downregulated for 96h siU6atac-treated LNCaP, C4-2, 22Rv1, cells and PM154 organoids, respectively (**Fig. 4F,** Supplementary Table 9). For example, we found the proliferation marker Ki67 not only to be downregulated in the transcriptome but also in the proteome of LNCaP and C4-2 cells. 22Rv1 cells showed a decrease in the DDR proteins BRCA1 and AURKA/B. In contrast, CDN1A (p21) was upregulated in transcriptome and proteome in all three cell lines. Upon independent submission of these four gene sets to DAVID, we found a high degree of GO Term overlap that mostly centered on cell cycle (**Fig. 4G**). For example, genes with similarly downregulated transcripts and encoded proteins in LNCaP, C4-2, and 22RV1 cells enriched for cell division, mitotic nuclear division, spindle pole, sister chromatid cohesion, G2/M transition of mitotic cell cycle, cell proliferation, and DNA strand elongation involved in DNA replication (**Fig. 4G**). Similarly, the 8 genes downregulated both transcriptionally and proteomically in the siU6atac-treated PM154 organoid enriched for a single GO Term – mitotic metaphase plate congression – which was also enriched by the siU6atac-treated 22RV1 cells (**Fig. 4G**). Taken together, siU6atac-mediated MiS inhibition altered the transcriptome and the proteome, which together enriched for cell cycle regulators. Based on the molecular signature, we next explored whether MiS inhibition would indeed affect cell cycle progression. Therefore, we employed FACS analysis at 72 and 96 hours post transfection (Supplementary Fig. S4.4A), which revealed a significant increase (two-way ANOVA, 72h: ∗∗p =0.0027, 96h: ∗∗p =0.0018) in G1/G0 phase cells and a significant decrease (two-way ANOVA, 72h∗∗p =0.0080, 96h: ∗∗p =0.0085) in S-phase cells in therapy-resistant C4-2 cells treated with siU6atac (**Fig. 4H**). PM154 organoids showed a similar result under siU6atac conditions (**Fig. 4I**). These cell cycle defects observed here revealed that MiS inhibition provokes a G1/S cell cycle arrest in PCa.

### Single cell RNAseq reveals cell cycle defects triggered by MiS inhibition

Next, we wanted to identify cellular heterogeneity in the response to siU6atac mediated MiS inhibition. We thus also performed single cell RNAseq (scRNAseq) on LNCaP cells and PM154 organoids post siU6atac. After standard data processing and quality control procedures (methods) we obtained transcriptomic profiles for 8206 siScrambled and 6730 LNCaP cells and 11181 siScrambled and 11475 PM154 organoids (Supplementary Table 10). We employed unsupervised clustering to identify heterogeneity in response to siU6atac. K-means clustering and UMAP projection of the combined data across both genotypes, which revealed 9 major cell clusters (**Fig. 5A**). We found that clusters 0, 3, 4, 6, 7 and 9 were populated by an equal number of cells treated with siScrambled and siU6atac (**Fig. 5B**). Clusters 1(Fisher’s exact test, p=3.16934e-69), 2 (Fisher’s exact test, p=1.702664e-63) and 5 (Fisher’s exact test, p=3.977987e-05) showed higher percentages of siScrambled cells, relative to siU6atac cells. Finally, cluster 8 (Fisher’s exact test, p=1.533612e-14) was predominantly populated by cells treated with siU6atac compared to siScrambled (**Fig. 5B**). GO Term enrichment analysis of the Top 5 upregulated genes, in the cells belonging to clusters 1, 2 and 5 revealed cell division, G1/S transition and mitotic nuclear division (**Fig. 5C**). In contrast, there was no Go Term enrichment for the Top 5 genes in cluster 8 (**Fig. 5C**). The enrichment of cell cycle terms combined with previous findings that MiS inhibition impacts cell cycle regulation in cancer (**Fig. 3,4**) led us to employ a list of cell cycle regulators as a means of sorting single cells based on cell cycle stage^38^. Thus, we superimposed cell cycle stage on the UMAP representation of the unsupervised data clustering (**Fig. 5D**). We found that clusters 0, 3, 4, 6, 7 and 9 showed the highest percentage of G1 cells (**Fig. 5E**), whereas clusters 1 and 2 showed the highest percentage of cells in S and G2/M phase respectively (**Fig. 5E**). Similar analysis for PM154 organoids yielded 7 clusters (Supplementary Fig.S5.2A). A majority of the cells populating cluster 5 (Fisher’s exact test, p=3.287745e-40) were from siScrambled, while cluster 6 (Fisher’s exact test, 2.738288e-57) was predominated populated by siU6atac-treated cells (Supplementary Fig. S5.2B). There was no Go Term enrichment for genes upregulated in those clusters, yet cluster 5 (Supplementary Fig, S5.2C), as with clusters 1, 2 and 5 of LNCaP cells, showed a high percentage of cells in S and G2/M phase, respectively (Supplementary Fig. S5.2D, E).

**Figure 5.**
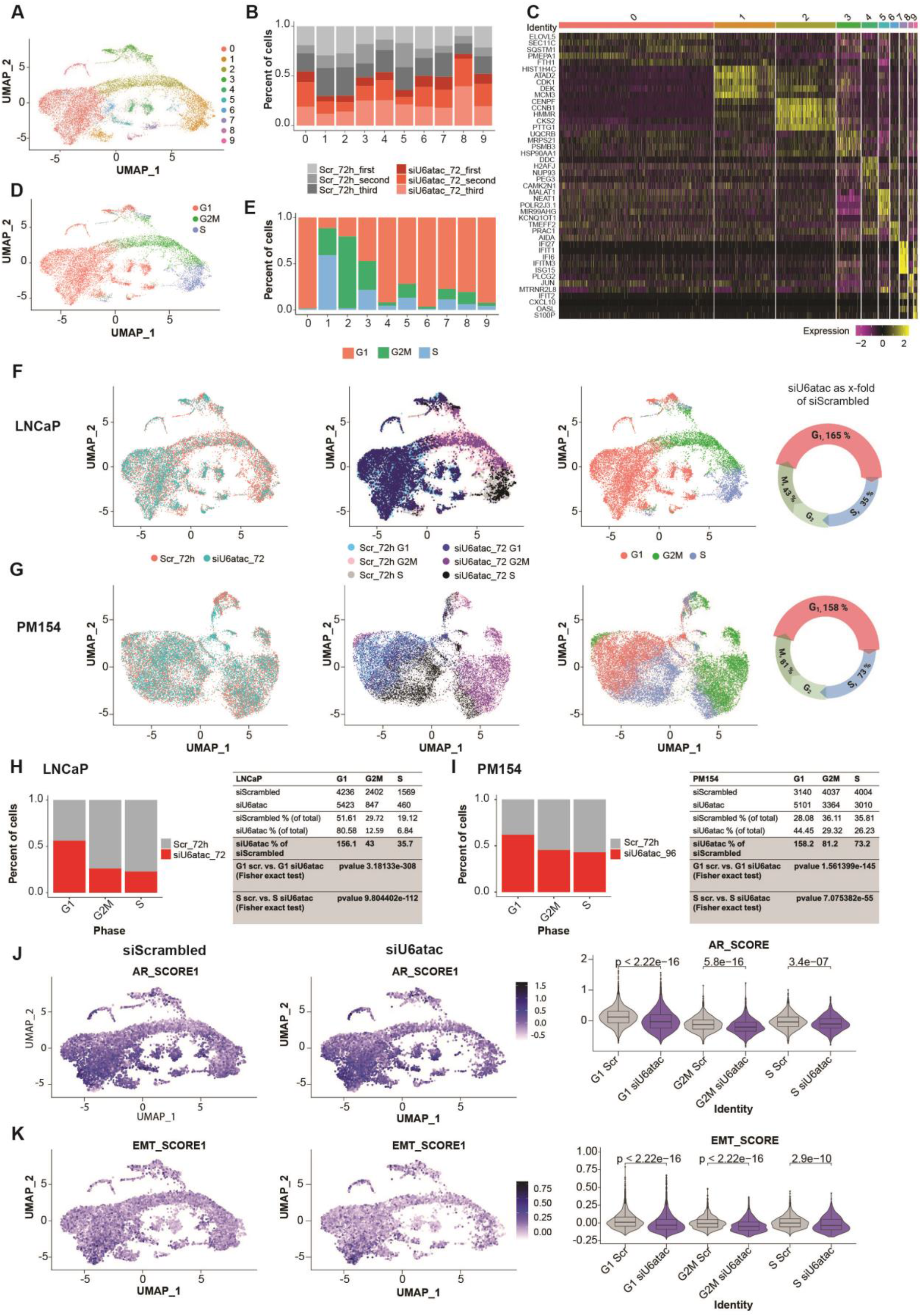
single cell RNAseq corroborates siU6atac-mediated cell cycle defects and reveals PCa lineage dependency on MiS function. (A) UMAP representation of LNCaP cell line showing the clusters at the optimal resolution (0.2). (B) Histogram of LNCaP cell line showing the contribution of each sample in each cluster. (C) Heatmap of LNCaP cell line showing the expression of top 5 marker genes in each cluster. (D) UMAP representation of LNCaP cell line showing the cell cycle phase of LNCaP cell line for each cell. (E) Histogram of LNCaP cell line showing the percentage of cell cycle phase in each cluster. (F) UMAP representation of the LNCaP cell line showing the contribution of siScrambled and siU6atac samples (on the left); the cell cycle phase of siScrambled and siU6atac samples (on the center) and cell phase of each cell (on the right). (G) UMAP representation of PM154 cell line showing the contribution of siScrambled and siU6atac samples (on the left); the cell cycle phase of siScrambled and siU6atac samples (on the center) and cell phase of each cell. (H) Histogram of LNCaP cell line showing the percentage of the cell cycle phase for siScrambled and siU6atac cells. The table is showing the number of cells for each cell cycle phase. The p-value was calculated using a Fisher’s exact test. (I) Histogram of PM154 cell line showing the percentage of the cell cycle phase for siScrambled and siU6atac cells. The table is showing the number of cells for each cell cycle phase. The p-value was calculated using a Fisher’s exact test. (J) UMAP representation of LNCaP cell line showing the AR score calculated on siScrambled and siU6atac samples. Violin plots show the AR score calculated for each cell cycle phase. The p-value was calculated using a Wilcoxon test. (K) UMAP of LNCaP cell line showing the EMT score calculated on siScrambled and siU6atac samples. Violin plots show the EMT score calculated for each cell cycle phase. The p-value was calculated using a Wilcoxon test

The comparison of distribution of cells from each condition in the 9 clusters and the distribution of cells in the different phases of cell cycle led us to investigate cell cycle defects through scRNAseq analysis (**Fig. 5F-I**). Therefore, we superimposed the cell identities (i.e., siScrambled or siU6atac) onto the UMAP of the clusters along with the cell cycle phases (**Fig. 5 F-I**). For LNCaP cells, we found a significant enrichment (Fisher’s exact test, p=3.18e-308, OR=3.89, 95% CI 3.60 – 4.19) of siU6atac-treated cells in G1 phase and a reduction of these cells in S phase (Fisher’s exact test, p=9.80e-112, OR=3.22, 95% CI 2.88 – 3.60) (**Fig. 5F and H**). Similarly, for PM154 organoids, we observed a significant enrichment of siU6atac cells in G1 (Fisher’s exact test, p=1.56e-145, OR=2.05, 95% CI 1.93 – 2.16) and a decrease in S-phase (Fisher’s exact test, p=7.07e-55, OR=1.57, 95% CI 1.48 – 1.66) (**Fig. 5G and I**). These findings are consistent with FACS analysis (**Fig. 4H and I**).

Given that siU6atac is successful at blocking cell cycle progression in PCa and that AR signaling is a crucial driver of PCa, we next explored the AR score (a crucial metric of CRPC-adeno progression) in siU6atac-treated LNCaP cells and PM154 organoids (**Fig. 5J** and Supplementary Fig. S5.2F). We discovered that siU6atac was indeed able to reduce the AR scores of both models significantly in G1 (Wilcoxon test, p<2.22e-16), G2M (Wilcoxon test, LNCaP: p=5.8e-16, PM154: p<2.22e-16) and S-phase (Wilcoxon test, LNCaP: p=3.4e-07, PM154: p<2.22e-16) (**Fig. 5J**). We did not observe any shift in the AR score post siU6atac treatment in bulk RNAseq (Supplementary Fig. S5.2H), which is not surprising, as we lose single cell resolution and dilution of expression of some of the key AR score markers. Another hallmark of cancer progression is EMT, which we investigated by using an EMT score^39^. We found that EMT score was also significantly reduced in G1 (Wilcoxon test, p<2.22e-16), G2M (Wilcoxon test, LNCaP: p=2.9e-10, PM154: p=2.9e-06) and S-phase (Wilcoxon test, LNCaP: p=3.4e-07, PM154: p=0.0089) in siU6atac-treated cells and organoids (**Fig. 5K** and Supplementary Fig. S5.2G). Next, we looked at the expression of MIGs that were identified as most crucial nodes from our PPI analysis in the scRNAseq data (**Fig. 1B and C**). Indeed, we found a significantly lower number of siU6atac-treated cells and organoids that expressed these MIGs such as AURKA, EED, and PARP1 (Supplementary Fig. S5.3 and S5.4).

Together, scRNAseq revealed siU6atac mediated transcriptomic remodeling that contributes to PCa progression and lineage plasticity by decreasing cell cycle, AR signaling and EMT.

### The minor spliceosome is essential for PCa growth and viability

Application of siU6atac significantly decreased proliferation in hormone responsive LNCaP cells and in therapy-resistant L-AR, C4-2, 22RV1 and PM154 cell and organoids (**Fig.6A** and Supplementary Fig. S6.1A and B). Previous reports have shown tissue-specific minor intron splicing as reflected by differential rate of minor intron retention across various tissues^11^. Thus, we explored whether siU6atac treatment resulted in a similar, context dependent failure to survive. Indeed, we observed a significant decrease in confluence of L-AR cells compared to LNCaP cells upon siU6atac treatment (**Fig. 6B,** Supplementary Fig. S6.2C). L-AR cells showed a higher proliferation defect than LNCaP cells, most likely due to higher doubling rate of L-AR cells (Supplementary Fig. S6.1B). L-AR cells (as well as therapy-resistant cells) were shown to possess higher MiS activity than their native hormone responsive lines. These data suggest that a higher basal MiS activity **(Fig. 2)** is indicative of an increased dependency on the minor spliceosome. Of note, viability of non-cancer cells, such as fibroblasts (HS27) and primary mouse prostate cells (M2514), were not affected by siU6atac-mediated MiS inhibition (**Fig. 6A** and Supplementary Figs. S6.1A and 6.3). In accordance with the KD data, overexpression of U6atac in C4-2 cells re-enforces cell growth (**Fig. 6C**), which agrees with previous findings showing that the MiS is essential for cell proliferation^40, 41^ and plays a major role in cell cycle progression.

**Figure 6.**
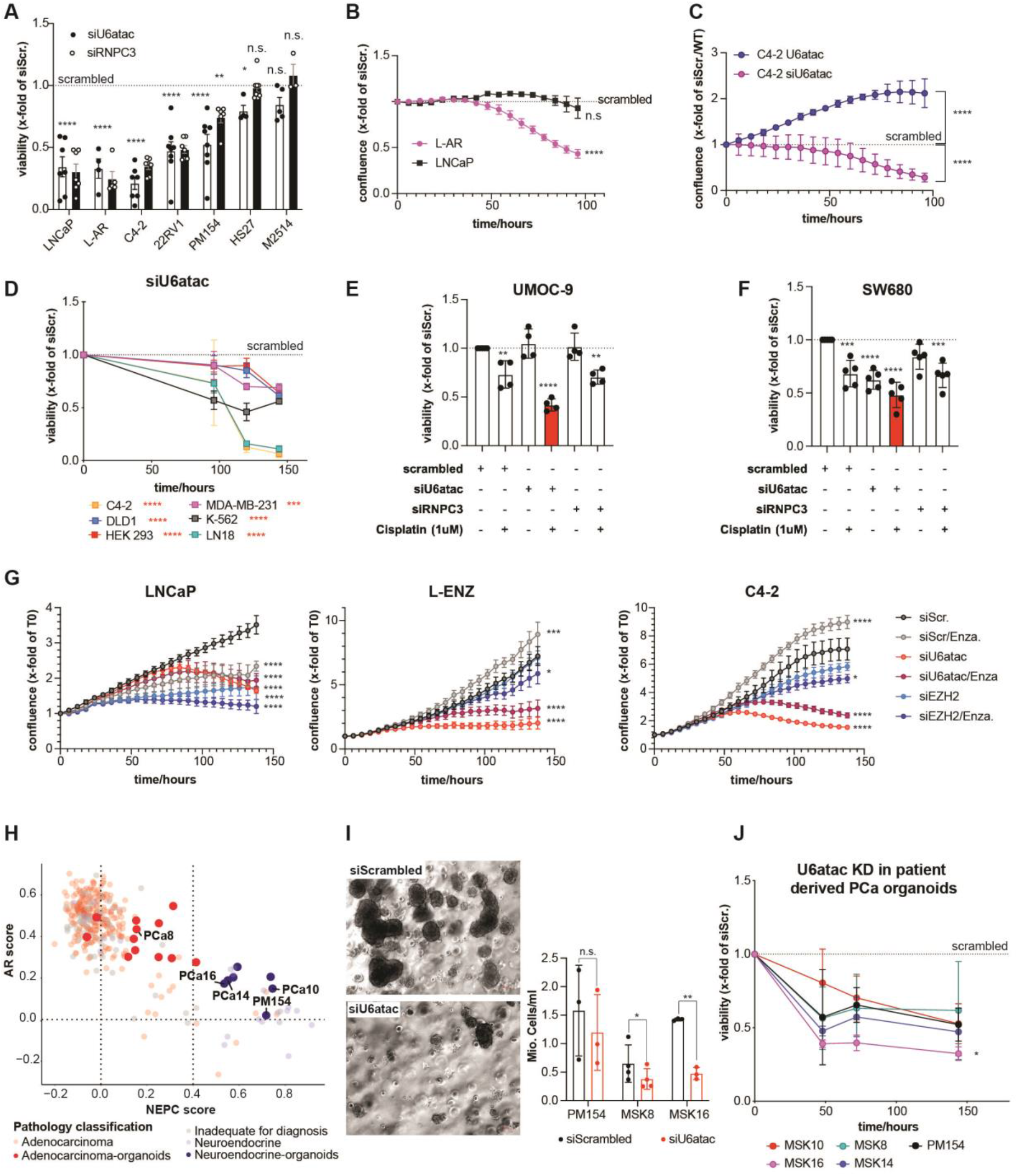
U6atac and the MiS represent potential therapeutic targets in cancer. (A) Cell viability in human PCa cell lines (LNCaP n=5, L-AR n=4, C4-2 n=6, 22Rv1 n=6, Pm154 n=6), human fibroblast cells (HS27 n=6) and primary mouse prostate cells (MS2514 n=3, MS2513 n=3) (mean ± SEM, ordinary two-way Anova; ns p > 0.05, ∗p < 0.05, ∗∗p < 0.01, ∗∗∗p < 0.001). Experiments were performed in triplicates. (B) Growth curves of LNCaP and L-AR cells treated for several time points with siU6atac. Data is normalized to the siScrambled control and represents pooled results from 4 biologically independent experiments (mean +/-SEM, ordinary two-way Anova; ns p > 0.05, ∗p < 0.05, ∗∗p < 0.01, ∗∗∗p < 0.001). (C) Growth curves of C4-2 cells stable overexpressing U6atac normalized to C4-2 cells expressing the EV plasmid or C4-2 cells treated for several time points with siU6atac normalized to C4-2 cells treated with siScrambled. Data represents pooled results from 16 (C4-2 siU6atac) and 4 (C4-2 U6atac) biologically independent experiments (mean +/- SD ordinary two-way Anova; ns p > 0.05, ∗p < 0.05, ∗∗p < 0.01, ∗∗∗p < 0.001). (D) Cell viability in different human cancer cell lines treated for several time points with siU6atac. Data are normalized to the siScrambled control and represents pooled results from 4 biologically independent experiments (mean +/-SEM ordinary two-way Anova; ns p > 0.05, ∗p < 0.05, ∗∗p < 0.01, ∗∗∗p < 0.001). (E) Cell viability in human bladder cancer UMOC-9 cells treated with siRNA against U6atac, RNPC3 and Scrambled or with Cisplatin (1uM). Data represents pooled results from 4 biologically independent experiments (mean +/-SD, ordinary one-way Anova; ns p > 0.05, ∗p < 0.05, ∗∗p < 0.01, ∗∗∗p < 0.001). (F) Cell viability in human bladder cancer SW680 cells treated with siRNA against U6atac, RNPC3 and Scrambled or with Cisplatin (1uM). Data represents pooled results from 4 biologically independent experiments (mean +/-SD, ordinary one-way Anova; ns p > 0.05, ∗p < 0.05, ∗∗p < 0.01, ∗∗∗p < 0.001). (G) Growth curves of LNCaP (n=4), L-ENZ (n=6) and C4-2 (n=4) cells treated with siScrambled, siU6atac and siEZH2 +/- Enzalutamide (10uM). Data represents pooled results from biologically independent experiments (mean +/- SEM, ordinary two-way Anova; ns p > 0.05, ∗p < 0.05, ∗∗p < 0.01, ∗∗∗p < 0.001). (H) Scatter blot representing RNAseq results from patient derived PCa organoids (big dots) superimposing clinical data from the SU2C dataset (small dots). Y-axis represents the AR-score, x-axis represents the NEPC score. (I) Brightfield microcopy of MSK8 cells 8 days after siRNA treatment against U6atac. Cells treated with Scrambled siRNA are shown as control. Scale bars (red): 50 μm. Bar blot summarizes cell counts at day 8 (n=3, paired t-test, *p=0.0307). (J) Cell viability in patient derived PCa organoids. Data is normalized to the scrambled control and represents pooled results from three biologically independent experiments (mean +/- SEM, two-way Anova; ns p > 0.05, ∗p < 0.05, ∗∗p < 0.01, ∗∗∗p < 0.001).

While we targeted U6atac to inhibit the MiS, the minor spliceosome complex consists of other unique components, including snRNAs and snRNPs. Therefore, we chose to inhibit the MiS by targeting RNPC3, another component of the MiS. Indeed, RNPC3 KD provoked similar reactions in PCa cells to siU6atac, indicating that the observed decrease in cell proliferation and viability can be attributed to a decrease in MiS activity in general and is not a U6atac-specific observation (**Fig. 6A** and Supplementary Fig. S6.1A and B). Importantly, we found that MiS inhibition not only affects PCa, but that the viability of other cancer cell lines such as colon, kidney, breast, glioblastoma, and CML was also decreased by a KD of several MiS components **(Fig. 6D** and Supplementary Fig. S6.2C and D). This finding reveals that the MiS is essential for other cancer types and that dependency on its integrity might represent a broad phenomenon that is not restricted to PCa alone. Particularly, among all tested siRNAs (RNPC3, PDCD7, U12 and U6atac), a U6atac KD turned out to be the most efficient. Surprisingly, even KD of U6atac was not sufficient to decrease viability of the bladder cancer lines UMOC9 and SW680. However, siU6atac (in combination with cisplatin) led to a significant decrease in viability **(Fig. 6E and F)**, suggesting that, in certain cases, MiS inhibition might work synergistically with current state-of-the-art treatments. In fact, CRPC requires combination therapy like inhibition of EZH2, which sensitizes towards ARSi (enzalutamide) treatment^42, 43^. Therefore, we tested whether U6atac inhibition plus enzalutamide might be similar or better than EZH2 inhibition plus enzalutamide. We observed that U6atac KD plus enzalutamide was more effective than EZH2 plus enzalutamide in CRPC C4-2 cells and in enzalutamide-resistant LNCaP (L-rENZ) cells (**Fig. 6G** and Supplementary Fig. S6.2D). Surprisingly, U6atac KD alone was significantly better than EZH2 plus enzalutamide.

Finally, we wanted to further explore the effects of MiS inhibition in a model system that captures PCa disease heterogeneity better than the previously used cell culture studies. Therefore, we applied siRNA against U6atac in PCa patient-derived organoids that were cultured in 3D. Superimposing the RNAseq results of those organoids with clinical data from the SU2C study **(Fig. 6H)** confirmed the clinical relevance of those model systems. Treatment of the organoids MSK8 (PCa8, CRPC-Ad), MSK10, 16, 14 (PCa 10,16,14, CRPC-NE) and PM154 (CRPC-NE) with siU6atac provoked a significant (two-way ANOVA, p=0.0246) reduction (Msk16: P=0.019) in organoid growth and viability **(Fig. 6I and J** and Supplementary Fig. S6.2E). Collectively, these data show that knocking down MiS components (especially U6atac) is sufficient to decrease cancer growth and viability.

In summary, we conducted the first-in-field evaluation of MiS in PCa using a multi-pronged approach that combined RNAseq, mass spectrophotometry, FACS, and scRNAseq. This demonstrates that MiS inhibition results in severe cell cycle defects through aberrant splicing of MIGs, which fundamentally impacts PCa progression, survival and lineage plasticity

## Discussion

### MIG-expression discriminates cancer from benign tissue

MIG-expression and the pathways they regulate are intimately linked to oncogenes (**Fig. 1A**). Thus, MIGs are critical bottlenecks for disparate molecular pathways downstream of these oncogenes, and as such are part of the aberrant regulatory logic of cancer related phenotypes. Therefore, the MiS likely represents another master regulator of cancer whose inhibition would trigger tumour checkpoint collapse^20^.Indeed, MIG expression was discriminatory between different cancer types, while levels of minor intron retention were informative in PCa progression (**Fig. 1**). Thus MIGs, with diverse functions, are ideal effectors of the disparate pathways driven by cancer-causing genes. Moreover, minor intron splicing is potentially a crucial regulatory node for cancer progression, which is consistent with the idea of increased cell cycle speed that cancer progression necessitates. In agreement with these findings, we found MiS snRNAs (crucial regulators of MiS activity) to be upregulated in cancer and across cancer progression (**Fig. 1I-L** and **Fig. 2A**). These findings suggest that MiS activity is enhanced and actively regulated across cancer progression. Moreover, increased U11, PDCD7 and RNPC3 expression have been associated with PCa, AML and increased anti-RNPC3 cancer associated scleroderma respectivey^18, 27, 28^. Even more drastic upregulation was observed for the MiS catalytic core snRNA U6atac, which is controlled through turnover rather than transcriptional upregulation. According to the “molecular switch theory,” U6atac is rapidly turned over and, as such, is the limiting snRNA that dictates MiS activity. Cellular stress signalling and, among others, p38MAPK activation, causes a rapid increase in U6atac expression along with enhanced splicing of MIGs that are otherwise inefficiently spliced or degraded^35^. Thus, stabilization of U6atac, which elevates its levels, can enhance minor spliceosome activity. Re-phrased, U6atac levels can control MIG expression “on-demand” when required. Consistent with this model, U6atac upregulation was observed in tumours from various tissues compared to their respective benign counterparts (**Fig. 1L**). This finding underscores the idea that U6atac snRNA levels are a crucial regulator of minor spliceosome activity during cancer progression.

### MiS activity increases with cancer progression

MIG expression is uniquely dependent on the splicing efficiency of minor introns. As such, it is not surprising that we found differential efficiencies for the splicing of minor introns when we compared different PCa subtypes. Indeed, studying U6atac levels during progression of prostate cancer showed that U6atac expression is closely correlated to the progression of PCa and is positively associated with PCa metastasis (**Fig. 2A**). Support for this idea was observed in minor intron splicing reporters, which were more efficiently spliced in cell lines representing more aggressive cancer stages (**Fig. 2B**). For example, splicing of minor introns, unlike major introns, was more efficient in different therapy-resistant CRPC-adeno cells and in various CRPC-NE organoids (**Fig. 2B).** Considering the “molecular switch theory,” MIG regulation may thus represent a dynamic mechanism of adaptation by cancer cells in response to changing environmental conditions, as seen in therapy resistance. In this sense, we found that MIG splicing increases with prolonged ARSi treatment of prostate cancer. Resistance to ARSi is almost always based on a re-activation of the AR axis in prostate cancer, which lines up with our data showing that MIG splicing is regulated by the AR in PCa (**Fig. 2D-F**). These findings, along with our previous discovery of U6atac as a powerful modulator of MiS activity, strongly suggest that controlling U6atac levels is a point of therapeutic intervention. Together these findings reveal a unique role for MiS activity in the onset and progression of cancer and positions the MiS as a potential therapeutic target.

### siU6atac blocks cell cycle of cancer cells

Given that U6atac snRNA has no other known function outside of the MiS, siU6atac specifically inhibited MiS function, which was reflected in elevated minor intron retention (**Fig. 3**). In fact, siU6atac resulted in the largest number of MIGs with elevated minor intron retention compared to published reports of MiS inhibition with other components^44–46^. Unlike MiS inhibition through other components^13, 37, 45^, the largest AS events observed in siU6atac samples were cryptic splicing events executed by the major spliceosome (**Fig. 3D**). MiS inhibition through siU6atac is expected to disrupt the U4atac-U6atac-U5 tri-snRNP that is recruited after 5’ splice site recognition by the U11/U12 di-snRNP^47, 48^. Thus, the exon definition requirement is mostly fulfilled and exon-skipping by the major spliceosome is less likely (**Fig. 3D, red**). This finding bolsters the idea that inhibition of the minor spliceosome induces specific types of AS events depending on the specific MiS component that is targeted and the cell type in which it is targeted. This finding opens a new way to target snRNAs, which allows inhibition of the minor spliceosome directly.

The cell type-specific response to MiS inhibition was reflected in the different numbers of MIGs with altered minor intron splicing for each cell line (**Fig. 3C, E**). Indeed, principal component analysis with either minor intron retention or AS showed that the response to MiS inhibition in C4-2 is closer to LNCaP, whereas 22Rv1 is closer to PM154 (Supplementary Fig. S3.3). This finding is consistent with our understanding that LNCaP, C4-2, 22Rv1 and PM154 can be placed in that order in PCa progression. The separation of PM154 organoids from the three cell lines was also evident in the overlap analysis of minor intron retention and AS events (Supplementary Fig. S3.2). STRING network analysis showed how simultaneous aberrant splicing of MIGs converge upon a central node composed of downregulated genes that predominantly enrich for cell cycle regulation. (**Fig. 3H**). Interestingly, the same analysis performed for the PM154 organoids did not yield similar results. This suggests that MiS inhibition is indeed cell type specific, tumour stage and context dependent, which is consistent with hierarchical clustering of 9 cancer types using MIG expression (**Fig. 1D**). However, IPA analysis with PCa interacting MIGs and downregulated genes in the three cell lines and organoid showed distinct cancer-related biological pathways (**Fig. 3I-L**). This underscores the idea that MiS inhibition can perturb disparate MIGs in different cell lines yet converge on the same biologically relevant endpoint for cancer (i.e., cell cycle and survival). For example, biological network analysis revealed a MAPK-centric network for the adeno cell lines whereas in the organoids a PARP1, ACTL6A, SRPK1 driven network was observed. The latter finding is consistent with the idea that the CRPC-NE organoids are molecularly distinct from the three CRPC-adeno cell lines.

Perturbation of predicted biological pathways based on transcriptomic changes are ultimately implemented by the proteins produced. Mass spectrophotometry analysis showed downregulation of many MIGs that were aberrantly spliced. However, we did not observe a one-to-one correlation (**Fig. 4F**), which is consistent with published reports about the discordance between the transcriptome and proteome^49, 50^. Nonetheless, proteins (MIGs and non-MIGs) that are up or downregulated in the siU6atac condition were relevant to PCa progression. For example, AURKA, which was shown to be important in CRPC differentiation and aggressiveness^5, 36^, was increased in therapy-sensitive LNCaP cells but decreased in CRPC-adeno cells 22Rv1 and CRPC-NE organoids PM154. Similarly, the polycomb group protein EED, known to regulate AR expression levels^51^ and a potential target in CRPC^52, 53^ was decreased in C4-2 and 22Rv1 cells. In contrast, we observed an increase in the protein levels for the G1 cell cycle arrest mediator CDN1A in all CRPC-adeno cells (**Fig. 4A-E**). While there is inherent discrepancy between the transcriptome and proteome, we found that genes with downregulation of both transcript and protein highly enriched for cell cycle regulation and DDR GOTerms (**Fig. 4G**). Thus, despite the dynamic fluctuations in the levels of mRNA and protein, the core molecular defect is that of inhibition of proliferation. Indeed, FACS analysis confirmed G1/G0 arrest in C4-2 cells and PM154 organoid (**Figs. 4H and I**).

### ScRNAseq reveals reduction in AR- and EMT-score in siU6atac-treated cells and organoids

Cell cycle defects observed by FACS analysis (**Fig. 4H and I**) were further confirmed by scRNAseq analysis, which revealed a high number of siU6atac treated cells in G1-phase (**Fig. 5F-I**). We also observed increase in the number of siU6atac treated cells in S-phase, which is consistent with the enrichment of the DDR pathway (Supplementary Table 7). Again, these findings underscore the cell type specific effects of MiS inhibition that ultimately result in a block in proliferation, a key feature for any therapeutic strategy. Consistent with this, we observed a decrease in EMT activity after MiS inhibition, which implies that targeting the MiS not only blocks cancer progression but also metastasis (Fig. 5K and supplementary Fig. S5.2). The main strategy in PCa therapy however is to block AR activity. Intriguingly, we found that MiS inhibition led to reduced AR activity score in PCa cells and organoids (**Fig. 5J** and Supplementary Fig. S5.2) This reveals inhibition of AR transcriptional activity as a novel mechanism of action of the MiS, which is potentially cancer type or even cancer stage dependent. Considering that MiS activity, and thus MiS-dependent signaling, is tissue and cancer type-specific, this finding indicates that the MiS may represent a dynamic mechanism of adaption for cancer cells in response to changing environmental conditions. The MiS may thus be a common denominator of prominent cancer driver axis such as the AR axis in PCa, that could be exploited as an all-in-one target for many cancer types.

### The minor spliceosome is essential for PCa growth and viability

Whereas siU6atac-mediated MiS inhibition substantially impacted cancer cells, it did not strongly affect human fibroblasts or benign mouse prostate cells (**Fig. 6A**). This finding has therapeutic relevance as it appears that MiS inhibition is specifically detrimental to highly proliferative cells. Although U6atac KD resulted in less viability of LNCaP cells, their proliferative index was not significantly affected by a MiS inhibition (**Fig. 6B**). This difference in response by LNCaP cells can be explained by the slow rate of cell division, which has been shown to be dependent on AR activation^54^. Consistent with previous results that nominated the AR as a regulator of MiS activity, we noticed that AR overexpression increased proliferation of LNCaP cells, thereby making them more susceptible to siU6atac-mediated MiS inhibition (**Fig. 6B**). Thus, U6atac is a promising target for CRPC-adeno PCa with persistent (re)-activation of the AR^55^. Our study underscores the importance of inhibiting the MiS, which consists of other crucial components besides U6atac. Indeed, we found that the use of siRNA against the unique MiS protein RNPC3 led to similar effects as seen with siU6atac in PCa (**Fig. 6A).** However, we found that MiS inhibition can have differential effects based on both the cell type it is inhibited in and the MiS component targeted for its inhibition. For example, we found that siU6atac, while being quite broadly effective, is best in PCa and glioblastoma cells (**Fig. 6D).** Similarly targeting other MiS components such as RNPC3 and PDCD7 showed significant results in PCa and glioblastoma cells (Supplementary Fig. S.6.2C). In contrast, U12 KD had no significant effect on viability in all cell lines tested (Supplementary Fig. S6.2C).

Interestingly, U6atac KD alone did not impact bladder cancer cells, but it was very effective when combined with cisplatin (**Fig. 6E and F).** This finding suggests U6atac inhibition might be a valid addition for combination therapy for bladder cancer. Considering that many invasive bladder cancer patients are cisplatin-ineligible, a combination of cisplatin and siU6atac might represent a promising alternative to treat those patients^56^. Similarly, enzalutamide-resistant CRPC could benefit from combination therapy that includes siU6atac. Enzalutamide alone increased CRPC growth in resistant cells, which emphasizes the need for new CRPC treatments (**Fig. 6G**). For instance, it has been shown that EZH2 inhibition overcomes enzalutamide resistance in cultured PCa cells and xenograft models^43^. Surprisingly, we found that U6atac KD by itself is sufficient to block proliferation of CRPC cells, which is better than enzalutamide, siEZH2 or siEZH2/enzalutamide (**Fig. 6G**). Importantly, we show that MiS inhibition also works in PCa patient-derived organoids (CRPC-adeno and CRPC-NE), whose transcriptome signature correlates with those from patients tested in the SU2C study (**Fig. 6H**).

Taken together, we show that MiS activity plays a crucial role in the progression of PCa and that MiS inhibition is a viable therapeutic target. We show that inhibiting different MiS components can block cancer cell proliferation and viability, but in our study, U6atac was the most effective. We show that MiS activity corresponds to AR signalling across stages of PCa progression, which is reflected by the increase in U6atac expression. Indeed, this discovery suggests that U6atac could also be employed as a diagnostic marker for lethal PCa and other cancers. Regardless of the stage of PCa, siU6atac can successfully inhibit proliferation and viability through disruption of pathways such as MAPK, cell cycle and DNA repair. While in the current study PCa is used as a model to investigate the efficacy of MiS inhibition as a therapeutic strategy, these observations could extend to other cancer types (summarized in the graphical abstract).

## Acknowledgments

We are grateful to the patients and their families that have participated in genomic, transcriptomic, and precision cancer care studies. We would like to thank Marc-David Ruepp from King’s College for providing the p120, hSCN4A and hSCN8A minigenes. As well, Gideon Dreyfuss from the Perelman School of medicine for providing the luciferase minor intron reporter construct, Charles Sawyers and Ping Mu at Memorial Sloan-Kettering Cancer Center for sharing the LNCaP-AR cell line, Inti Zlobec, Carmen Cardozo and Stefan Rheinhard from the translational research unit (TRU) at the University of Bern for their assistance with the Scorenado TMA scoring platform. We are grateful to Antonio Rodriguez from the University of Bern for providing the Bern PCBM cohort TMA. We are equally grateful to the Next Generation Sequencing platform at the University of Bern in particular Pamela Nicholson and Daniela Steinert for their assistance with bulk and scRNAseq. We thank the Proteomics Mass Spectrometry Core Facility at the University of Bern in particular Sophie Braga and Anne-Christine Uldry for their assistance with the LC/ LC-MS experiments. We acknowledge expert assistance from Mariana Ricca and Kellie Cotter at the University of Bern in preparing the manuscript for submission. This project has received funding from the SAKK Astellas GU-Oncology Award 2020 and from the Fondaction Young Investigator Research Grant 2021.

## Author’s contributions

**A.A.**: Conceptualization, data curation, formal analysis, funding acquisition, investigation, methodology, project administration, resources, supervision, validation, visualization, writing original draft. **K.D.**: formal analysis, methodology/software, validation, visualization. **D.C.**: formal analysis, methodology/software, validation, visualization. **S.K.**: formal analysis, methodology/software, validation, visualization. **E.Q.:** formal analysis, visualization. **S.R.L.:** formal analysis, visualization**. L.R.**: formal analysis, methodology/software, validation, visualization. **J.G.**: formal analysis, validation, visualization. **S.P**: Investigation **M.J.**: data curation. **J.A.G.:** Data curation, methodology**. S.W.:** methodology/software, validation, visualization. **M.B.:** formal analysis, methodology/software, validation, visualization. **J.T.**: conceptualization, data curation, writing-review & editing **G.T.**: Resources. **M.KdJ.:** Resources. **J.P.T.:** Investigation. **M.G.**: Investigation. **Y.C.:** Resources. **R.K.:** Conceptualization, investigation, methodology, supervision, validation, visualization, writing original draft, writing-review & editing. **M.A.R.**: Funding acquisition, Investigation, resources, supervision, writing-review & editing.

## STAR Methods

### Resource availability Lead contact

Further information and requests for resources and reagents should be directed to and will be fulfilled by the Lead Contact, Mark Rubin and Rahul Kanadia.

### Material availability

This study did not generate new unique reagents.

The bulk RNAseq and scRNA-seq data generated during this study have been submitted on the European Genome-phenome Archive under the accession EGAS00001005546.

The mass spectrometry proteomics data that support the findings of this study have been deposited to the ProteomXchange Consortium (http://proteomecentral.proteomexchange.org) via the PRIDE partner repository with the dataset identifier PXD026949.

### Experimental model and subject details

#### Cell and Organoid lines

LNCaP (male, ATCC, RRID: CVCL_1379), C4-2 (male, ATCC, RRID: CRL-3314), 22Rv1 (male, ATCC, RRID: CRL-2505), PC3 (male, ATCC, RRID: CRL-3470), DLD-1 (2 (male, ATCC, RRID: CCL-221), L-rENZ and L-AR cells were maintained in RPMI medium (Gibco, A1049101), supplemented with 10% FBS (Gibco, 10270106), and 1% penicillin-streptomycin (Gibco, 11548876) on poly-L-lysine coated plates. RWPE cells (male, ATCC, RRID: CVCL_3791) were maintained in Keratinocyte Serum Free Medium (Gibco, 17005075) supplemented with bovine pituitary extract and human recombinant EGF (included), and 1% penicillin-streptomycin (Gibco, 11548876). HEK293T cells (female, ATCC, RRID: CVCL_0063), VCaP (male, ATCC, RRID: CRL-2876), MDA-MB-231 (female, ATCC, RRID: HTB-26), K-562 (female, ATCC, RRID: CCL-243), LN-18 (male, ATCC, RRID: CRL-2610) PC-3M-Pro4 and DU145 cells (male, ATCC, RRID: CVCL_0105) were maintained in DMEM (Gibco, 31966021), supplemented with 10% FBS, and 1% penicillin-streptomycin. NCI-H660 cells (male, ATCC, RRID: CRL-5813) were maintained in RPMI medium (Gibco, A1049101), supplemented with 5% FBS, 1% penicillin-streptomycin (Gibco, 11548876), 0.005 mg/ml Insulin (Sigma-Aldrich, I9278), 0.01 mg/ml Apo-Transferrin (Sigma-Aldrich, T1147), 30nM Sodium selenite (Sigma-Aldrich, S9133), 10 nM Hydrocortisone (Sigma-Aldrich, H6909) 10 nM beta-estradiol (Sigma-Aldrich, E2257) and L-glutamine (for final conc. of 4 mM) (Sigma-Aldrich, G7513). PC-3M-Pro4 cells were a kind gift from Dr. Kruithof-De Julio. LNCaP-AR cells were a kind gift from Dr. Sawyers and Dr. Mu (Memorial Sloan Kettering Cancer Center)^57^. L-ENZ cells were established through constant enzalutamide exposure. Briefly low passaged LNCaP cells were treated over night with 20uM enzalutamide in C/S media. The media was exchanged to normal RPMI (10% FBRS, 1% P/S) the next day and surviving LNCaP cells (∼10%) were maintained until they reached a confluency of ∼80%. This procedure was repeated twice. Subsequently the enzalutamide concentration was increased for three treatments to 40uM and for 25 treatments to 80uM. Cells are treated since them once a week with 80uM enzalutamide.

All cell lines were grown at 37 °C with 5% CO_2_. All cell lines were authenticated by STR analysis and regularly (every 3 month) tested for mycoplasma.

MSKCC-PCa8,10,14 and 16 CRPC-Adeno patient derived organoids were a kind gift from Dr. Chen^31^ (Memorial Sloan Kettering Cancer Center). All organoids including WCM154 were maintained in three-dimension according to the previously described protocol^30, 31^. Briefly Advanced DMEM (Thermo Fisher Scientific, 31966047) with GlutaMAX 1x (Thermo Fisher Scientific, 35050061), HEPES 1mM (Thermo Fisher Scientific, 15630056), AA 1x (Life Technologies, 15240-062), 1% penicillin-streptomycin, B27 (Thermo Fisher Scientific,17504001), *N*-Acetylcysteine 1.25 mM (Sigma-Aldrich, A9165), Recombinant Murine EGF 50 ng/ml (PeproTech, 315-09), Human Recombinant FGF-10 20 ng/ml (Peprotech, 100-26), Recombinant Human FGF-basic 1 ng/ml (Peprotech, 100-18B), A-83-01 500 nM (Tocris, 29-391-0), SB202190 10 μM (Sigma-Aldrich, S7076), Nicotinaminde 10 mM (Sigma-Aldrich, N0636), (DiHydro) Testosterone 1 nM (Fluka, 10300), PGE2 1 μM (Tocris, 2296), Noggin conditioned media (5%) (PeproTech, 120-10C) and R-spondin conditioned media (5%) (PeproTech, 315-32). The final resuspended pellet was mixed with growth factor-reduced Matrigel (VWR, BDAA356239) in a 1:2 volume ratio. Droplets of 40 μl cell suspension/Matrigel mixture were pipetted onto each well of a six-well cell suspension culture plate (Huberlab, 7.657185) To solidify the droplets the plate was placed into a cell culture incubator at 37 °C and 5% CO_2_ for 30 min. Subsequently 3 ml of human organoid culture media was added to each well. 50 % of the media was exchanged every 3−4 day during organoid growth. organoids were passaged as soon as they reached a size from 200 to 500 um. To this end, organoid droplets were mixed with TrypLE Express (Gibco) and placed in a water bath at 37 °C for a maximum of 5 min. The resulting cell clusters and single cells were washed and re-cultured, according to the protocol listed above.

#### In situ validation collection

Tissue micro-arrays were kindly provided by the Translational Research Unit (TRU) Platform, Bern (www.ngtma.com). For PCa we used TMAs from the Bern PCBM cohort^58^ (28 patients) and a tissue microarray of 210 primary prostate tissues, part of the European Multicenter High Risk Prostate Cancer Clinical and Translational research group (EMPaCT)^59–61^.

### Method details

#### Mass spectrometry analysis

LNCaP, C4-2, 22Rv1 and PM154 cells (400 000) were seeded in a 6 well and treated for 96 hours with siScrambled or siU6atac RNA (16 pmol). 96 hours post transfection cells were harvested and 50% of the cell pellet was used for U6atac KD confirmation by qRT-PCR. The remaining pellet was washed twice with PBS and subjected to mass spectrometry (MS) analysis:

Cells were lysed in 8M urea/100mM Tris pH8 / protease inhibitors with sonication for 1 minute on ice with 10 seconds intervals. The supernatant was reduced, alkylated and precipitated overnight. The pellet was re-suspended in 8M urea/50mM Tris pH8 and protein concentration was determinate with Qubit Protein Assay (Invitrogen).10µg protein was digested with LysC 2hours at 37C followed by Trypsin at room temperature overnight. 800ng of digests were loaded in random order onto a pre-column (C18 PepMap 100, 5µm, 100A, 300µm i.d. x 5mm length) at a flow rate of 50µL/min with solvent C (0.05% TFA in water/acetonitrile 98:2).

After loading, peptides were eluted in back flush mode onto a home packed analytical Nano-column (Reprosil Pur C18-AQ, 1.9µm, 120A, 0.075 mm i.d. x 500mm length) using an acetonitrile gradient of 5% to 40% solvent B (0.1% Formic Acid in water/acetonitrile 4,9:95) in 180min at a flow rate of 250nL/min. The column effluent was directly coupled to a Fusion LUMOS mass spectrometer (Thermo Fischer, Bremen; Germany) via a nano-spray ESI source.

Data acquisition was made in data dependent mode with precursor ion scans recorded in the orbitrap with resolution of 120’000 (at m/z=250) parallel to top speed fragment spectra of the most intense precursor ions in the Linear trap for a cycle time of 3 seconds maximum.

#### Generation of U6atac overexpressing cell lines

LV290591 – RNU6ATAC Lentiviral Vector (Human) (CMV) (pLenti-GIII-CMV-GFP-2A-Puro) as well as the corresponding empty vector control were purchased from ABM. DNA was amplified via chemical transformation of One Shot Mach1 T1 Phage-Resistant Chemically Competent E. coli cells (Invitrogen, C862003). Lentivirus was produced in HEK293T cells by transfection with the constructs, and subsequent virus containing media was used to transduce C4-2 cells. Three days post transduction the cells were subjected to puromycin selection (1 µg/mL). After the selected cells reached a confluence of 80%, they were FACS sorted for GFP positivity. This was repeated 3 times.

#### Drug treatments

For DHT stimulation experiments cells were starved of hormone for 48 hours in phenol red-free RPMI media (Gibco, 11-835-030) with 10% charcoal stripped FBS (Gibco, A3382101), then treated with 10 nM dihydrotestosterone (Fluka, 10300) for 24 hours.

For long-term ADT treatment cells were exposed weekly to 20 µM enzalutamide (Selleck Chemicals, S1250) or 10 uM Abiraterone.

For growth experiments cells were treated with siRNA and two hours later with 20uM enzalutamide. Enzalutamide was refreshed 3 days later.

For splicing reporter assays cells were exposed to Anisomycin 1ug/ml for 4h (Sigma Aldrich, A9789).

#### Proximity Ligation assay

L-AR cells (50 000) were seeded in µ-Slide 8 Well (ibidi, 80826). The next day cells were washed once in ice-cold PBS and fixed in 4% PFA for 10 minutes. Subsequently cells were permeabilized with PBS +0.2% Triton for 10 minutes. Proximity ligation assay using the Duolink® In Situ Red Starter Kit Mouse/Rabbit (Sigma-Aldrich, DUO92101-1KT) was performed according to the manufacturer’s instructions. Briefly primary monoclonal antibodies against mouse-AR (Thermo Fisher Scientific, MA5-13426), rabbit-HSP90 (Abcam, ab203085), rabbit-PDCD7 (Abcam, 121258) and rabbit IgG Isotype Control antibody (Thermo Fisher Scientific, 026102) were diluted in Duolink Antibody Diluent (1:50, 1:200, 1:100 and 1:1000). Cells were incubated in the AB solution over night at 4C. The next day cells were washed twice and incubated for one hour at 37C in a moisture chamber with PLUS and MINUS PLA probes. Subsequently cells were washed twice and incubated at 37C (humidity chamber) for 30 min in the ligation mix and 100 minutes in the amplification solution. After two final washes for 10 minutes slides were mounted with DAPI containing media and monitored with a fluorescence microscope (LEICA, DMI4000 B).

#### Cell transfection and siRNA-mediated knock-down

##### Cells

ON-TARGET plus siRNA SMARTpool siRNAs against *U6atac, AR, EZH2, PDCD7, mouse RNPC3* and the Non-targeting (siScrambled) siRNA were purchased from Dharmacon. siRNAs against RNU6atac and RNU12 and the Silencer Select Negative Control were purchased from Thermo Fisher Scientific and siRNA against mouse U6atac was purchased from Ambion. Transfection was performed for the respective timepoints on attached cells using the Lipofectamine™ RNAiMAX Transfection Reagent (Thermo Fisher Scientific, 13778150) to the proportions of 16pmol of 20 μM siRNA per well.

##### Organoids

Before transfection organoids were cultured for 2-3 weeks in human organoid growth medium. Media was removed and organoids were first mechanically dissociated. To obtain single cells organoids were trypsinized in 1ml TriplE (Thermo Fisher Scientific, 12605036) for 15-18 minutes at 37C. The reaction was stopped with 1ml growth media and cells were spun for 5 minutes at 300g. Subsequently the cells were strained and counted. Per condition one million cells were plated in a 6 well. Lipofectamine™ RNAiMAX complexes were prepared according to the standard Lipofectamine™ RNAiMAX protocol. In short, 5ul of RNAiMAX reagent and 40 nM of siRNA plus 10% FBS were each diluted in 125 ul Opti-MEMH medium. Both mixes were pooled and incubated for 10 minutes before the siRNA-reagent complex was added to the cells. Cell/siRNA mix was centrifuged at 600 g at 32C for 60 min, and then incubated over night at 37C. The next day cells were resuspended and collected by centrifugation (300g, 5min, RT). The pellet was resuspended in 280 ul Matrigel and the mix was separated into 7 drops that were added into a 6 well. Organoids were grown in human organoid media for 96h (CTG assay) or seven days (cell counting assay).

#### RNA extraction from cells and qRT-PCR

Cells were harvested for RNA isolation using the ReliaPrep™ miRNA Cell and Tissue Miniprep System (Promega, Z6212). Synthesis of complementary DNAs (cDNAs) using FIREScript RT cDNA Synthesis Kit (Solis BioDyne, 06-15-00200) and real-time reverse transcription PCR (RT-PCR) assays using HOT FIREPol EvaGreen qPCR Mix Plus (Solis BioDyne, 08-24-00020) were performed using and applying the manufacturer protocols. Quantitative real-time PCR was performed on the ViiA 7 system (Applied Biosystems). All quantitative real-time PCR assays were carried out using three technical replicates. Relative quantification of quantitative real-time PCR data used GAPDH, ACTB as housekeeping genes. Primer sequences are listed in Supplementary Table 11.

#### Single cell sequencing

Cell counting and viability assessments were conducted using a ViCell XR Cell counter and viability analyzer (Beckman Coulter, BA30273). Thereafter, GEM generation & barcoding, reverse transcription, cDNA amplification and 3’ gene expression library generation steps were all performed according to the Chromium Next GEM Single Cell 3’ Reagent Kits v3.1User Guide (10x Genomics CG000204 Rev D) with all stipulated 10x Genomics reagents. Generally, 11.8.-27.5 µL of each cell suspension (600-1’400 cells/µL) and 15.7-31.4 µL of nuclease-free water were used for a targeted cell recovery of 10’000 cells. GEM generation was followed by a GEM-reverse transcription incubation, a clean-up step and 10-12 cycles of cDNA amplification. The resulting cDNA was evaluated for quantity and quality using a Thermo Fisher Scientific Qubit 4.0 fluorometer with the Qubit dsDNA HS Assay Kit (Thermo Fisher Scientific, Q32854) and an Advanced Analytical Fragment Analyzer System using a Fragment Analyzer NGS Fragment Kit (Agilent, DNF-473), respectively. Thereafter, 3ʹ gene expression libraries were constructed using a sample index PCR step of 11-12 cycles. The generated cDNA libraries were tested for quantity and quality using fluorometry and capillary electrophoresis as described above. The cDNA libraries were pooled and sequenced with a loading concentration of 300 pM (150 pM in runs using XP workflow), paired end and single indexed, on an illumina NovaSeq 6000 sequencer using a NovaSeq 6000 S2 Reagent Kit v1.5 (100 cycles; illumina 20028316) and two NovaSeq 6000 S4 Reagent Kits v1.5 (200 cycles; illumina 20028313). The read set-up was as follows: read 1: 28 cycles, i7 index: 8 cycles, i5: 0 cycles and read 2: 91 cycles. The quality of the sequencing runs was assessed using illumina Sequencing Analysis Viewer (illumina version 2.4.7) and all base call files were demultiplexed and converted into FASTQ files using illumina bcl2fastq conversion software v2.20. All steps were performed at the Next Generation Sequencing Platform, University of Bern.

#### Bulk RNA sequencing

Total RNA was extracted from LNCaP, C4-2, 22Rv1 and PM154 cells treated for 96h with siU6atac or siScrambled. The recommended DNase treatment was included. The quantity and quality of the extracted RNA was assessed using a Thermo Fisher Scientific Qubit 4.0 fluorometer with the Qubit RNA BR Assay Kit (Thermo Fisher Scientific, Q10211) and an Advanced Analytical Fragment Analyzer System using a Fragment Analyzer RNA Kit (Agilent, DNF-471), respectively. Thereafter cDNA libraries were generated using an illumina Stranded Total RNA Prep, Ligation with Ribo-Zero Plus (illumina, 20040529) in combination with IDT for Illumina RNA UD Indexes Sets A and B (Illumina, 20040553 and 20040554, respectively). The illumina protocol was followed exactly with the recommended input of 100 ng total RNA. The quantity and quality of the generated NGS libraries were evaluated using a Thermo Fisher Scientific Qubit 4.0 fluorometer with the Qubit dsDNA HS Assay Kit (Thermo Fisher Scientific, Q32854) and an Advanced Analytical Fragment Analyzer System using a Fragment Analyzer NGS Fragment Kit (Agilent, DNF-473), respectively. As a further quality control step, prior to NovaSeq 6000 sequencing, the pooled cDNA library pool underwent paired end sequencing using iSeq 100 i1 reagent v2, 300 cycles (illumina, 20040760) on an iSeq 100 sequencer. The library pool was re-pooled to ensure an equal number of reads/library and then paired end sequenced using a NovaSeq 6000 S4 reagent kits v1.5, 300 cycles (illumina, 20028312) on an Illumina NovaSeq 6000 instrument. The quality of the sequencing runs was assessed using illumina Sequencing Analysis Viewer (illumina version 2.4.7) and all base call files were demultiplexed and converted into FASTQ files using illumina bcl2fastq conversion software v2.20. The average number of reads/ libraries was 82 million. The RNA quality-control assessments, generation of libraries and sequencing runs were performed at the Next Generation Sequencing Platform, University of Bern, Switzerland.

#### U6atac in situ hybridization

mRNA ISH was performed by automated staining using Bond RX (Leica Biosystems) and Basescope® technology (Advanced Cell Diagnostics, Hayward, CA, USA). All slides were dewaxed in Bond dewax solution (product code AR9222, Leica Biosystems) and heat-induced epitope retrieval at pH 9 in Tris buffer based (code AR9640, Leica Biosystems) for 15 min at 95° and Protease treatment for 5 min. The following probes from RNAscope 2.5 LS (Advanced Cell Diagnostics) were used: BaseScope™ LS Probe - BA-Hs-RNU6ATAC-1zz-st-C1ref 1039918, PPIB-1zz ref 710178 and DapB-1zz ref 701028, were used as positive and negative control respectively. Probe efficiency was tested using U6atac overexpressing C4-2 cells (5 million) of which 50% were treated with siU6atac RNA.

All probes were incubated at 37° for 120 min. Basescope^TM^ 2.5 LS Assay (Ref 323600, Advanced Cell Diagnostics) was used as pre-amplification system. Subsequent the reaction was visualized using Fast red as red chromogen (Bond polymer Refine Red detection, Leica Biosystems, Ref DS9390) for 20 min. Finally, the samples were counterstained with Haematoxylin, air dried and mounted with Aquatex (Merck). Slides were scanned and photographed using Pannoramic 250 (3DHistech). U6atac intensity was scored manually by a pathologist (Mark Rubin) blinded to the clinical data, using the digital online TMA scoring tool Scorenado^62^ (University of Bern, Switzerland) especially developed for TMA scoring on de-arrayed spots.

For the analysis of TMA data, samples annotated as ‘center’ were used. U6ATAC score was calculated by multiplying the percentage of positive cells by the intensity. The sample with the highest score was used where more than one value was recorded for a block. Comparisons between groups were carried out using Wilcox test.

325 primary, 25 primary with metastatic potential and 32 metastatic samples, from 24 patients were used for the comparison of U6ATAC expression in PCa and PCBM.

#### Flow Cytometry

C4-2 and PM154 cells were seeded in a 6 well (500 000/well) and transfected with siU6atac or siScrambled RNA for 72 and 96 hours as previously described. Flow Cytometry cell cycle analysis was performed using the Click-iT™ EdU Alexa Fluor™ 488 Flow Cytometry Assay Kit (Thermo Fisher Scientific, C10420). Briefly EdU (10uM) was added into the media and cells were incubated for one hour at 37C. cells were washed with 1% BSA in PBS and fixed in 100ul Click-iT fixative for 15 minutes. After three additional washing step cells were permeabilized for 15 minutes in 100ul 1xClick-iT saponin based reagent. Click-iT reaction cocktail was prepared according to manufacturer’s instructions and 500ul reaction mix/ condition were incubated for 30 min with the cells at room temperature. Cells were washed and resuspended in 500 ul saponin-based permeabilization buffer. Hoechst (1ug/ml) was added 20 minutes prior analysis to the reaction mix. Cells were analyzed using the FACSDiva Software on a BD LSR II Flow Cytometer (BD Biosciences) in the FACSlab Core facility of the University of Bern. Data was further quantified with FlowJo 10.7.1. Values were calculated as fold-change as compared to siScrambled treated controls.

#### Immunoblotting

Cells were lysed in GST-Fish buffer (10 % (v/v) Glycerol, 50 mM Tris-HCl pH7,4, 100 mM NaCl, 1 % (v/v) Nonidet P-40, 2 mM MgCl_2_, 1 mM PMSF) with freshly added protease and phosphatase inhibitors. Total protein concentration was measured using the Pierce BCA Protein Assay Kit (Thermo Fisher Scientific). 50 ug protein samples were resolved 4-15% Mini-Protean TGX gels (BioRad, 456-1084) in SDS-PAGE and transferred to nitrocellulose membranes using the iBlot2 system (Thermo Fisher, IB23001). Blots were blocked for 1 hour at room temperature in 5% milk/TBST or BSA/TBST and incubated overnight at 4 °C with primary antibodies (Supplementary Table 11) which were dissolved in 5% BSA/TBST buffer. After 3 washes, the membrane was incubated with secondary antibody conjugated to horseradish peroxidase for 1 h at room temperature. After 3 washes, signal was visualized by chemiluminescence using the Luminata Forte substrate (Thermo Fisher Scientific, WBLUF0100) for strong antibodies and WesternBright Sirius-HRP Substrate (Witec AG, K-12043-D10) for weak antibodies. Images were acquired with the FUSION FX7 EDGE Imaging System (Witec AG).

#### Luciferase reporter assay

CMV-luc2CP/ARE (major intron splicing reporter), CMV-luc2CP (empty vector backbone control) and luc1CFH4 (minor splicing intron reporter) were a kind gift from Dr. Gideon Dreyfuss^29^ (University of Pennsylvania). DNA was amplified via chemical transformation of One Shot Mach1 T1 Phage-Resistant Chemically Competent E. coli cells (Invitrogen, C862003) and Sanger sequenced.

To determine the minor/major intron splicing rate cells were seeded in white 96 well plate (Huberlab, 7.655 098) (8000/well) and treated according to assay conditions. 24 hours prior analysis cells of each condition were co-transfected with each reporter plasmid and the empty vector backbone plasmid. In short 1.5ul of P3000 reagent plus 0.5 ug of DNA and 1.5 ul Lipofectamine P300 were each diluted in 25ul Opti-MEMH medium. Both mixes were pooled and incubated for 20 minutes before the solution was added to the cells. Luciferase expression was measured with the Dual-Glo® Luciferase Assay System (Promega, E2940): media was removed, 100ul of PLB were added and cells were frozen for two hours at −20C. After a one hour shaking step 100 ul of LAR substrate was added and Firefly luciferase expression was measured with a Tecan Infinite M200PRO reader. Values were calculated as x-fold of CMV-luc2CP expression and subsequently as the x-fold of the respective reference control.

#### Minigene reporter assay

pRSV2-p120-AmpR XL10 (p120), pCMV7.1 SCN4A frag (hSCN4A) and pLUX hSCN8A-minigene AmpR STbl3 (hSCN8a) were a kind gift from Dr. Mark-David Ruepp^63^ (King’s College London). DNA was amplified via chemical transformation of One Shot Mach1 T1 Phage-Resistant Chemically Competent E. coli cells (Invitrogen, C862003).

To determine the minigene splice index cells were seeded in a 6-well (350 000/well) and treated according to assay conditions. SiRNA was added 96 hours prior measurement. 72 hours prior measurement cells were transfected with the respective minigene. Briefly 10 ul of P3000 reagent plus 1.5 ug of DNA and 7.5 ul Lipofectamine P3000 were each diluted in 125ul Opti-MEMH medium. Both mixes were pooled and incubated for 20 minutes before the solution was added to the cells. 48 hours before the measurement media was exchanged for media with 10% charcoal stripped FBS (Gibco, A3382101) and 24 hours prior measurement 100 nM DHT was added to the respective condition. qRT-PCR was performed to verify the KD and to determine the splice index of each minigene. The minigene splice index was calculated by forming the ratio of normalized mRNA levels of cells transfected with the minigene versus mRNA levels of WT cells to consider the transfection efficiency. Subsequently the values corresponding to the spliced minigene were divided by the values corresponding to the unspliced minigene.

#### Cell growth experiments

##### Viability

Cells were seeded in a 6 well (400 000) and treated according to assay conditions over night. Cells were then seeded in Poly-L-Lysine coated 96-well plates (8000 cells/well, n=3 per condition) and WCM154 cells were seeded in a collagen-coated 96-well plates (5000 cells/well, n=3 per condition). Remaining cells were used for U6atac KD control via qRT-PCR. Cell viability was determined after 24, 48, 72, and 96 h with a Tecan Infinite M200PRO reader using the CellTiter-Glo® Luminescent Cell Viability Assay according to manufacturer’s directions (Promega, G9243). Viability values were calculated as *x*-fold of cells transfected with siRNA for 0 h. Cell confluence (n=4 per condition) was determined using the Incucyte S3 instrument and the IncuCyte S3 2018B software (Essen Bioscience, Germany). Values were calculated as x-fold of timepoint 0 and then as fold-change in confluency as compared to siScrambled treated controls.

##### Organoids

Organoids were transfected with siRNA as described previously. The following day 160000 cells were resuspended in 320 ul Matrigel and drops of 40ul (one drop/well, four timepoints, n=2) were plated in a suspension 48-well plate (Huberlab, 7.677 102). Remaining cells were plated in a 6-well for q RT-PCR U6atac KD control. The 24-well plate was incubated for three minutes in 37C and for 20 minutes upside down in 37C. Subsequently 500ul of organoid media were added and viability was measured using the CellTiter-Glo 3D Cell Viability Assay (Promega, G9683).

##### Co-immunoprecipitation

For the co-immunoprecipitation (co-IP), cytosolic fractions of LNCaP-AR cells were isolated using the Universal CoIP Kit (Active Motif, 54002). Chromatin of the cytosolic fraction was mechanically sheared using a Dounce homogenizer Fisher Scientific, 11898502). Cytosolic membrane and debris were pelleted by centrifugation and protein concentration of the cleared lysate was determined with the Pierce BCA Protein Assay Kit (Thermo Fisher Scientific, 23227). One microgram of the rabbit anti-PDCD7 (ab131258, Abcam) and rabbit anti-AR (ab133273, Abcam) antibodies and 1 μg of rabbit IgG Isotype Control antibody (Thermo Fisher Scientific, 026102) were incubated with 1 mg protein supernatant overnight at 4 °C with gentle rotation. The following morning, 30 μl of Protein G Magnetic Beads (Active Motif) were washed twice with 500 μl CoIP buffer and incubated with Antibody-containing lysate for 2 hours at 4 °C with gentle rotation. Bead-bound PDCD7 or AR complexes were washed twice with CoIP buffer and subsequently twice with a buffer containing 150 mM NaCl, 50 mM Tris-HCL (pH 8) and Protease and Phosphatase inhibitors. Washing procedure was executed at 4 °C with gentle rotation. Bead-bound protein, supernatant and Input controls were reduced and denatured in 40 μl Laemmli buffer containing DTT through boiling for 5 min at 95 °C. Magnetic beads were removed from solution using Magnetized Pipette Racks (Thermo Fisher Scientific, 11757325) and 20 μl of reduce protein was loaded on an SDS-PAGE gel with subsequent immunoblotting using iBlot (Life Technologies). Membranes were blocked in 5% BSA solution and then incubated over night with respective antibodies against targets of interest: AR, PDCD7, AR45 (Androgen Receptor Antibody (Carboxy-terminal Antigen), Cell Signaling Technologies, 54653S), ZRSR2 (Abcam, ab223062). Protein signal was detected using HRP-labeled native anti-rabbit IgG antibody (CST, #5127) and Luminata Forte substrate (Thermo Fisher Scientific, WBLUF0100) using the FUSION FX7 EDGE Imaging System (Witec AG).

### Quantification and statistical analysis

#### Protein-protein interaction network analysis

Assuming that cancer genes perturb a large molecular network through their interactions with other genes, distance to MIGs in a protein-protein interaction network of 160,881 interactions between 15,366 human proteins as of the HINT database was determined^21^. In particular, a MIG that directly interacts with a cancer-causing gene is a distance *d*=1 away, while a protein that is separated by a protein in between is a distance *d*=2 away from a cancer-genes in question. To find these interaction distances, a list of 403 cancer-causing genes from 186 different cancer types as of the Cancer Genome Interpreter database was considered (Supplementary Table 1) (https://www.cancergenomeinterpreter.org)^22^ in the underlying protein-protein interaction network of 160,881 interactions between 15,366 human proteins as of the HINT database^21^. A profile of the numbers of MIGs with a given distance *d* away from each cancer gene, 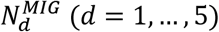 was obtained. To assess if the presence of MIGs in the vicinity of cancer genes is significant, sets of 542 MIGs found in the underlying interaction networks were randomly sampled. In particular, the corresponding distances of cancer genes to these randomly sampled MIGs was measured and a profile of numbers of random MIGs with a given distance *d* away from cancer genes 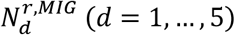 was obtained. The enrichment of MIGs is defined a distance *d* away from cancer genes as 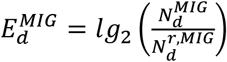, with average *E* over 100,000 random samples of MIGs.

To find a tight-knit web of MIGs and cancer genes, such direct interactions of cancer genes and MIGs were distracted in the underlying human protein-protein interaction network and determined the size of the largest connected subnetwork *S*. Randomly sampling MIGs N = 100,000 times. The sizes of the largest subnetworks was calculated using these random sets, *S_r_*, and the significance of *S* by 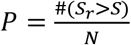 was determined.

#### Principal component analysis

MIG expression data from 3 sources: prostate samples from GTEx (nominally healthy prostate tissue), prostate cancer samples from TCGA (primary prostate cancer samples), and prostate cancer samples from SU2C (advanced prostate cancer) was merged. Gene expression values were normalized following the protocol adopted by GTEx (and detailed in our pan-cancer analysis). PCA analysis on this normalized gene expression matrix was carried out, thereby enabling visualization of the resultant data as the projections onto the space spanned by the first 2 PCs. Notably, this visualization appears to capture the progression from these 3 broad phases of prostate cancer progression, from healthy tissue in GTEx (at lower ends of the first PC) to advanced stages in SU2C (at higher ends of the first PC).

#### Quantitative comparisons between MIG- and non-MIG-based gene expression clustering

The Silhouette coefficient^64^ was used in order to characterize the relative performance of gene expression clustering for gene sets containing different relative abundances of MIGs and non-MIGs. The Silhouette coefficient provides an objective metric for measuring what is visually discerned to be structure (or any lack thereof) in a given heatmap (Supplementary Fig. S1.1 and 1.2). This coefficient constitutes an unsupervised approach to provide a score ranging −1 and 1, with scores closer to 1 indicative of well-defined and dense clustering (i.e., more meaningful structure in a given heatmap). This coefficient quantifies how similar a given data point (i.e., sample) is to its own cluster relative to different clusters. In our study, a data point consists of a N-length vector, where N is the number of distinct samples in an expression matrix, and the number of distinct clusters is pre-defined to be n_cluster = 9 in our pan-cancer analysis, since this analysis was carried out on a dataset of 9 distinct cancer types. The Silhouette coefficient has also been adopted for similar purposes in previous studies^23–26^

Prior to calculating the Silhouette coefficient, an agglomerative clustering on a normalized gene expression matrix was performed. Gene expression values from 9 cancer types (Biliary-, Breast-, ColoRect-, Lung-, Ovary-, Panc-, Prost-, Stomach-, Thy-adenocarcinoma) (Supplementary Fig. S1.2) totaling N=444 samples were taken from PCAWG, and the gene expression normalization was performed using the same approach as that adopted by GTEx (PMID: 29022597). Briefly, the entire gene expression matrix was normalized using quantile normalization. Then, inverse quantile normalization was applied to this quantile-normalized matrix in order to map to a standard normal (this also enabled us to remove outliers). For the PCa progression analysis, gene expression values from GTEx, TCGA, and SU2C were merged into one expression matrix (Supplementary Fig. S1.3). Exactly as we had done with the pan-cancer analysis, the pre-normalized expression matrix was normalized using quantile normalization, followed by inverse quantile normalization to map to a standard normal distribution; this step also removed outlier genes. Again, this normalization scheme was the same as that adopted previously by GTEx (PMID: 29022597). Using this normalized gene expression matrix, clustering was performed by using hierarchical clustering by employing the Euclidean metric and Ward linkage.

The entire analysis was run using different expression matrices with varying fractions of non-MIGs, and the results are plotted in Fig. 1F and G. Each box in this plot (for instance, the right-most box, which represents 100% non-MIG sets) corresponds to a 1000-length simulation in which non-MIGs are randomly sampled from among all non-MIGs in the genome. Thus, for the case of the right-most box, 1000 random sets are sampled, each of which has a composition of 100% non-MIGs. The P-value is based on a two-sided t-test in which each sample value is taken to be the difference between the Silhouette coefficient of one of the 1000 100% non-MIG samples and the corresponding Silhouette coefficient when only MIGs are used for clustering (i.e., the Silhouette coefficient corresponding to 0% non-MIGs).

#### MSI analysis

Primary tumor RNA-seq patient samples from The Cancer Genome Atlas (TCGA) were randomly queried using the GenomicDataCommons package in R (http://github.com/Bioconductor/GenomicDataCommons). The minor intron retention pipeline developed by Olthof et. al. (https://github.com/amolthof/minor-intron-retention)^11^ was run on the queried TCGA samples from the following cohorts: breast invasive carcinoma (BRCA; N=20), cholangiocarcinoma (CHOL; N=20), colon adenocarcinoma (COAD; N=20), lung adenocarcinoma (LUAD; N=20), ovarian serous cystadenocarcinoma (OV; N=20), pancreatic adenocarcinoma (PAAD; N=20), prostate adenocarcinoma (PRAD; N=16), and thyroid carcinoma (THCA; N=20). For the prostate cancer analyses, the following additional samples from the Stand Up to Cancer dataset were analyzed: androgen receptor (AR; N=20) and neuroendocrine (NE; N=22). NE samples were samples which had an NEPC score of greater than 0.4 while AR samples had an NEPC score of less than or equal to 0.4. Prostate cancer samples from the Genome-Tissue Expression Portal were analyzed as well (GTEX; N=20). A Kruskal-Wallis with post-hoc Dunn’s Test was performed between the TCGA cohorts and the prostate cohorts. A heatmap was generated with the gplots package in R with the default clustering method for the pan-cancer TCGA cohorts and the prostate cohorts across the MIGs (https://CRAN.R-project.org/package=gplots). A GO Enrichment Analysis was performed on the genes that clustered for each cancer cohort, grouping genes with a mean MSI value in the ranges of 0, 0 to 0.04, 0.04 to 0.75, and 0.75 to 1.

#### RNU11 quantification according to Gleason score

Gene-expression data of primary prostate cancer specimen was retrieved from The Cancer Genome Atlas (TCGA) in form of raw-counts. Sequencing reads were aligned to the human reference genome (hg38) using *STAR*^65^. Gene-expression was quantified at gene-level using *Gencode* annotations (v29)^66^. Subsequent analysis and library-size normalization were performed using *edgeR* pipeline^67^. RNU11 mapping reads were identified in 23 out of 497 samples (5%) which reflects the difficulty of capturing this gene product using canonical PolyA+ sequencing techniques. Nonetheless, a clear association between Gleason score and RNU11 mRNA expression in these 23 samples was identified (Supplementary Table 3). Significance was assessed using non-parametric Wilcoxon Test.

#### Single-cell RNAseq

##### Single-cell preprocessing and quality control

Cell ranger analysis pipeline v6.0.1 was used to align reads to the human genome reference sequence (GRCh38) and generate a gene-cell matrix from these data. The gene expression matrix was analyzed using Seurat 4.0.3 (https://github.com/satijalab/seurat)^68^. We removed low quality cells and multiplets by excluding genes detected in less than 5 cells and by discarding cells with more than 10000 fewer than 1000 detected genes. Cells containing mitochondrial gene counts greater than 25% were also removed.

UMI counts were normalized with the NormalizeData Seurat function using the LogNormalize normalization method with default parameters (10000 scale.factor).

##### Cell cycle phase classification and cell scores

Prediction of cell cycle phase for each cell was performed using Seurat CellCycleScoring function. A score was computed and a cell phase (G2/M, S and G1) was assigned to the cell as described previously^38^. Fisher’s exact test was performed to check whether the sIU6 cells have significantly a different number of cells than the Scr in G1 or S phase using the R fisher.test function.

We used Seurat AddModuleScore function to evaluate the degree to which individual cells express a certain pre-defined gene set. We defined scores to estimate the activities of prostate AR pathway, and EMT state, as described previously^39^. The AR pathway gene set included *AR*, *KLK3*, *KLK2*, *FKBP5*, *TMPRSS2*, *FOXA1*, *GATA2*, *SLC45A3* and EMT state *CDH2*, *CDH11*, *FN1*, *VIM*, *TWIST1*, *SNAI1*, *ZEB1*, *ZEB2* and *DCN*. Violin plots were drawn using Seurat and *p*-values were calculated using Wilcoxon test (Hadley Wickham, ggplot2: Elegant Graphics for Data Analysis 2016)

##### Multiple datasets integration and Batch correcting

For merging multiple datasets and minimizing the batch effect between them, we integrated our 6 samples (3 replicates SCR and 3 replicates siU6) for each cell line following the procedure of Seurat v4.0.3.^69^

Briefly, we selected the most variable genes for each dataset using the FindVariableFeatures function (selection.method =“vst”) and ranked them according to the number of datasets in which they were independently identified as highly variable. The 2000 most variable genes were thus integrated by merging pairs of datasets according to a given distance.

Integration anchors, representing two cells that are predicted to originate from a common biological state in both datasets using a Canonical Correlation Analysis (CCA), were done using the FindIntegrationAnchors function. The expression of the target dataset was corrected using the difference in expression between the two expression vectors for each pair of anchor cells. This step was performed using the IntegrateData function. This process resulted in an expression matrix containing the batch-effect-corrected expression for the 2000 selected genes for all cells from the 6 samples for each cell line.

##### Dimension reduction and clustering

A PCA was performed on the scaled data using RunPCA Seurat function (npcs = 30). Uniform Manifold Approximation and Projection (UMAP), a nonlinear dimension reduction method, was run using RunUMAP Seurat package function in order to embed cells in a 2-dimensional space. A K-nearest neighbor graph (KNN) based on the Euclidean distance in PCA space was constructed to cluster the cells with the Louvain algorithm (resolution = 0.2) using the FindNeighbors and FindClusters Seurat functions. We selected the optimal clustering resolution using the clustree R package(v0.4.3)^70^. Barplots were performed using dittoSeq R package^71^.

##### Differential gene expression analysis

Differentially expressed genes (DEGs) were identified between the different clusters using the FindAllMarkers function from the Seurat package (one-tailed Wilcoxon rank-sum test, *p* values adjusted for multiple testing using the Bonferroni correction). To compute DEGs, all genes were tested provided they were expressed in at least 25% of cells in either of the two compared populations, and the expression difference on a natural log scale was at least 0.25. The heatmap was produced using the DoHeatmap Seurat function by selecting the top five genes for each cluster.

#### RNAseq

##### Gene expression

Paired-end, total RNA reads for each replicate (N=4) were mapped to the mm10 genome via Hisat2 (Kim et al., 2015). Reads mapping to multiple locations were removed. Gene expression values were calculated using IsoEM2 (Mandric et al., 2017). Differential gene expression was calculated by IsoDE2 (Mandric et al., 2017), which uses 200 rounds of iterative bootstrapping to produce a 95% confidence interval for the expression of each gene, then statistically compares these values between experimental conditions. A threshold of log2FC ≥ 1, *P* ≤ 0.01 for upregulation, and log2FC ≤ −1, *P* ≤ 0.01 for downregulation was employed.

##### Minor intron retention

We report here minor intron retention as a mis-splicing index through the methodology described in Olthof et al., 2019. Briefly, uniquely mapped reads from the region of interest around minor introns (from two exons upstream to two exons downstream) were extracted. The mis-splicing index was then calculated by summing reads that map to the 5’ splice site and 3’ splice site of a minor intron, divided by the sum of reads that map to the 5’ splice site, 3’ splice site, and 2x canonically spliced reads. We only considered introns that pass our filtering criteria, which requires >4 exon-intron boundary reads, ≥1 read mapping to both the 5’ splice site and 3’ splice site, and >95% intron coverage, in all replicates of a condition as retained. Statistically significant global minor intron retention was determined using a Kruskal-Wallis test with post-hoc Dunn’s test (*P* ≤ 0.05). Determination of individual MIGs with significantly elevated minor intron retention was calculated using a two-tailed student’s T-test (*P* ≤ 0.05)

##### Alternative splicing

We employed the methodology reported by Olthof et al., 2017 for our alternative splicing analysis. Briefly, we used BEDTools to classify differential 5’ splice site and 3’ splice site usage around the region of interest for all minor introns and binned them into one of 8 categories (Fig. 3D). We then calculated a mis-splicing index by quantifying the number of AS reads divided by the sum of AS reads and canonically spliced reads per category per sample. We used a filtering criteria wherein we only considered introns to have AS if the average mis-splicing index for all replicates for each condition was > 10%. Additionally, we normalized the number of reads supporting an AS event by the total sequencing depth. As such, we only included AS events with >1 read per 3 million uniquely mapped reads for analysis. Determination of individual MIGs with significantly elevated AS was calculated using a two-tailed student’s T-test (*P* ≤ 0.05).

##### DAVID analysis

Gene lists were submitted to DAVID for gene ontology (GO) enrichment. We considered only GO Terms with Benjamini-Hochberg adjusted *P*-value ≤ 0.05 as significant.

##### STRING and Ingenuity Pathway Analysis

For LNCaP. C4-2, and 22Rv1, overlapping MIGs with significantly elevated minor intron retention that were also found to be associated with prostate cancer-causing genes were grouped with overlapping protein coding genes with significant downregulation. This list was submitted to STRING under the default parameters to obtain the gene interaction network. Subsequently, the same list was submitted to IPA as a core analysis using default parameters. All reported biological networks and pathways from IPA were significant using a Benjamini-Hochberg adjusted *P*-value ≤ 0.05 cut-off. The same analysis was performed for PM154 alone.

##### Principal component analysis

Principal component analyses were performed using clustvis (Metsalu & Vilo, 2015) with default parameters. Ellipses show 95% confidence interval.

##### GSEA analysis

To perform gene set enrichment analysis RNAseq data was pre-ranked using the metric: log10(Pvalue) / sign (logFC). GSEA was performed using the GSEA v.4.0.3 software. Hallmark gene sets, obtained from the GSEA website (www.broadinstitute.org/gsea/) was used for enrichment of siU6atac related pathway genes. Dotplot was used to visualize the most significant enriched terms. Normalised enrichment score (NES) and False discovery rate (FDR) were applied to sort siU6atac pathway enrichment after gene set permutations were performed 1000 times for the analysis.

#### MassSpec

MS data was interpreted with MaxQuant (version 1.6.1.0) against a SwissProt human database (release 2019_07) using the default MaxQuant settings, allowed mass deviation for precursor ions of 10 ppm for the first search and maximum peptide mass of 5500 Da; match between runs with a matching time window of 0.7 min was activated, but prevented between different groups of replicates by the use of non-consecutive fractions. Furthermore, the four cell lines were treated as different parameter groups and normalized independently. Settings that differed from the default also included: strict trypsin cleavage rule allowing for 3 missed cleavages, fixed carbamidomethylation of cysteines, variable oxidation of methionines and acetylation of protein N-termini.

Protein intensities are reported as MaxQuant’s Label Free Quantification (LFQ) values, as well as Top3 values (sum of the intensities of the three most intense peptides); for the latter, variance stabilization was used for the peptide normalization. Missing peptide intensities were imputed in the following manner, provided there was at least one identification in the group: two missing values in a group of replicates would be replaced by draws from a Gaussian distribution of width 0.3 x sample standard deviation centred at the sample distribution mean minus 1.8× the sample standard deviation, whereas a single missing value per group would be replaced following the Maximum Likelihood Estlimation (MLE) method. Imputation at protein level for both LFQ and Top3 values was performed if there were at least two measured intensities in at least one group of replicates; missing values in this case were drawn from a Gaussian distribution of width 0.3 x sample standard deviation centered at the sample distribution mean minus 2.5x the sample standard deviation. Differential expression tests were performed using the moderated t-test empirical Bayes (R function EBayes from the limma package version 3.40.6) on imputed LFQ and Top3 protein intensities. The Benjamini and Hochberg method was further applied to correct for multiple testing. The criterion for statistically significant differential expression is that the maximum adjusted p-value for large fold changes is 0.05, and that this maximum decrease asymptotically to 0 as the log2 fold change of 1 is approached (with a curve parameter of one time the overall standard deviation). The protein imputation step was repeated 20x so as to be able to flag those proteins that are persistently significantly differentially expressed throughout the cycles.

Imputed iTop3 was used to calculate relative protein abundances. Differential expression was calculated using the Empirical Bayes test. Protein upregulation and downregulation was determined by setting a threshold of Benjamini-Hochberg adjusted *P-*value ≤ 0.05, log2FC ≥ 1 or ≤ −1, respectively.

### Additional resources

#### Data sharing statement

The data sets, including the bulk RNAseq- and the scRNAseq-datasets will be made available through the European Genome-phenome Archive under the accession EGAS00001005546 upon publication of the current manuscript. The LC/LC-MS-datasets will be made available through the ProteomXchange Consortium (http://proteomecentral.proteomexchange.org) via the PRIDE partner repository with the dataset identifier PXD026949.

