## Supplementary material for "Minor intron splicing efficiency increases with the development of lethal prostate cancer": Suplementary Figures

[1] Department for BioMedical Research, University of Bern, 3008 Bern, Switzerland
[2] Physiology and Neurobiology Department, University of Connecticut, CT 06269, USA
[3] Institute of Pathology and Medical Genetics, University Hospital Basel, 4056, Basel, Switzerland
[4] Department of Computer Science, Yale University, New Haven, CT 06520, USA
[5] Yale College, New Haven, CT 06520, USA
[6] Program in Computational Biology and Bioinformatics, Yale University, New Haven, CT 06520, USA
[7] Department of Molecular Biophysics and Biochemistry, Yale University, New Haven, CT 06520, USA
[8] Institute of Oncology Research, 6500 Bellinzona, Switzerland
[9] Institute of Pathology, University of Bern, Bern, 3008 Bern, Switzerland
[10] Human Oncology and Pathogenesis Program (HOPP), Memorial Sloan Kettering, NY 10065, USA
[11] Department of Medicine, Memorial Sloan Kettering, NY 10065, USA
[12] Department of Medicine, Weill Cornell Medicine, NY 10065, USA
[13] Department of Urology, Inselspital, Bern University Hospital, 3008 Bern, Switzerland
[14] Faculty of Biomedical Sciences, Università della Svizzera italiana, 6900 Lugano, Switzerland
[15] Department of Computer Science, University of Miami, FL 33146, USA
[16] Sylvester Comprehensive Cancer Center, Univ. of Miami, FL 33136, USA
[17] Department of Biology, University of Miami, FL 33146, USA
[18] Department of Biomedicine, University of Basel, 4031 Basel, Switzerland
[19] Institute for Systems Genomics, University of Connecticut, Storrs, CT, USA
[20] Bern Center for Precision Medicine, University of Bern and Inselspital, 3008 Bern, Switzerland
[#] co-senior author

**Supplementary Fig. S1.1 (related to Fig. 1)**

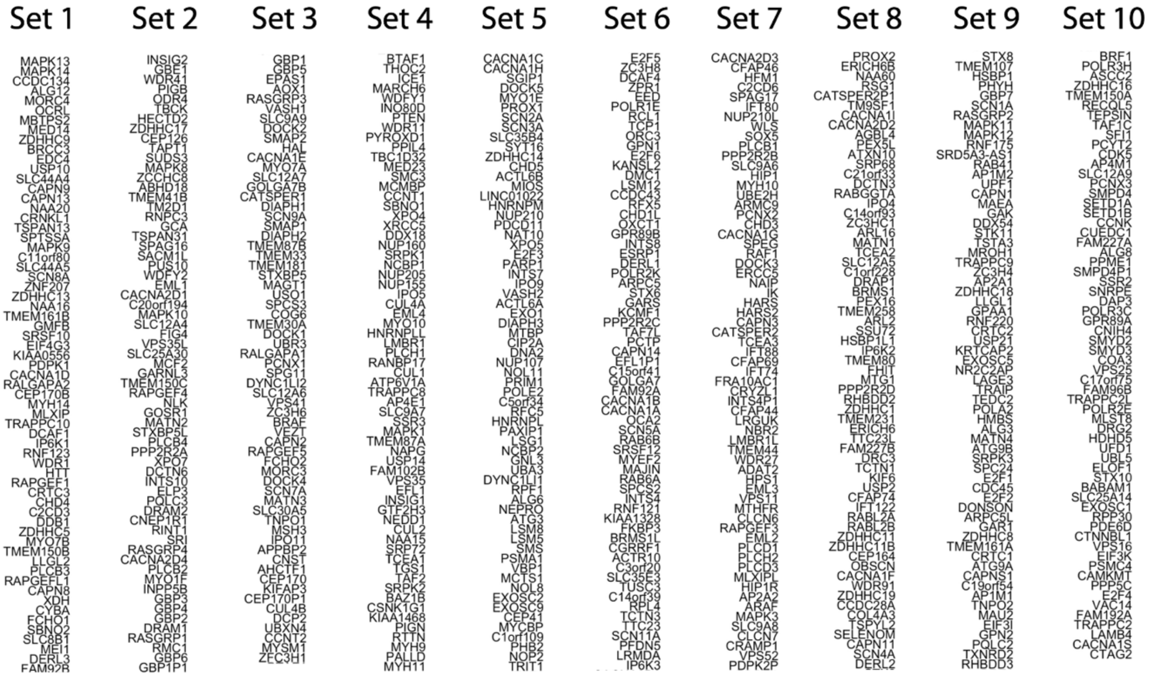


**Gene sets (1-10) used for heatmap in Fig. 1D**.

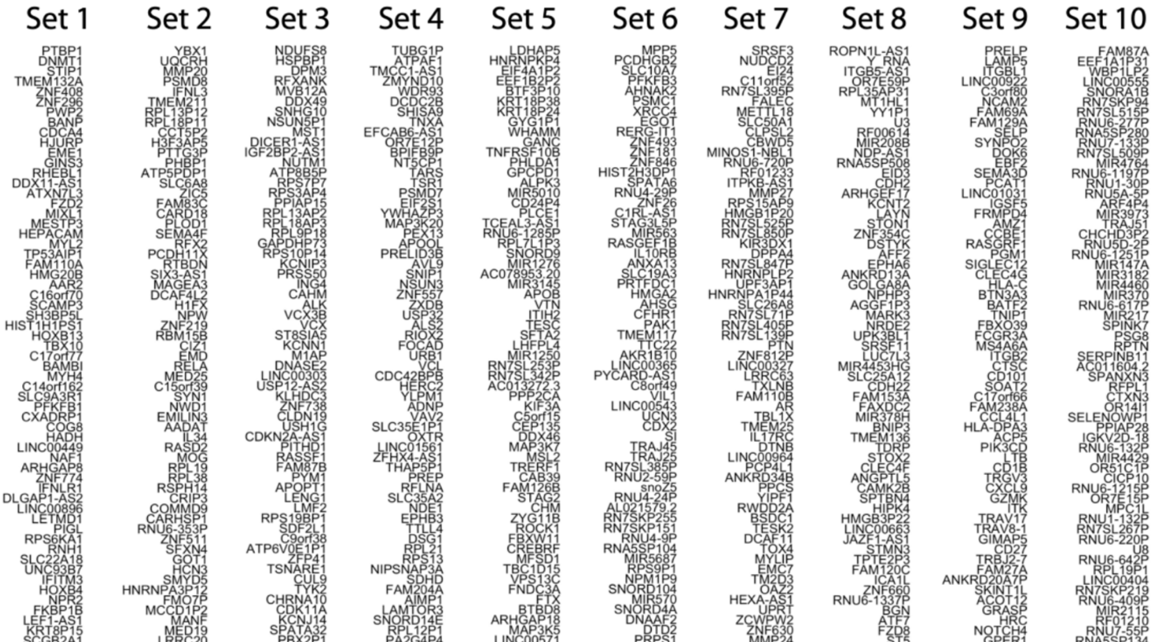


**Gene sets (1-10) used for heatmap in Fig. 1E**.

**Supplementary Fig. S1.2 (related to Fig. 1)**

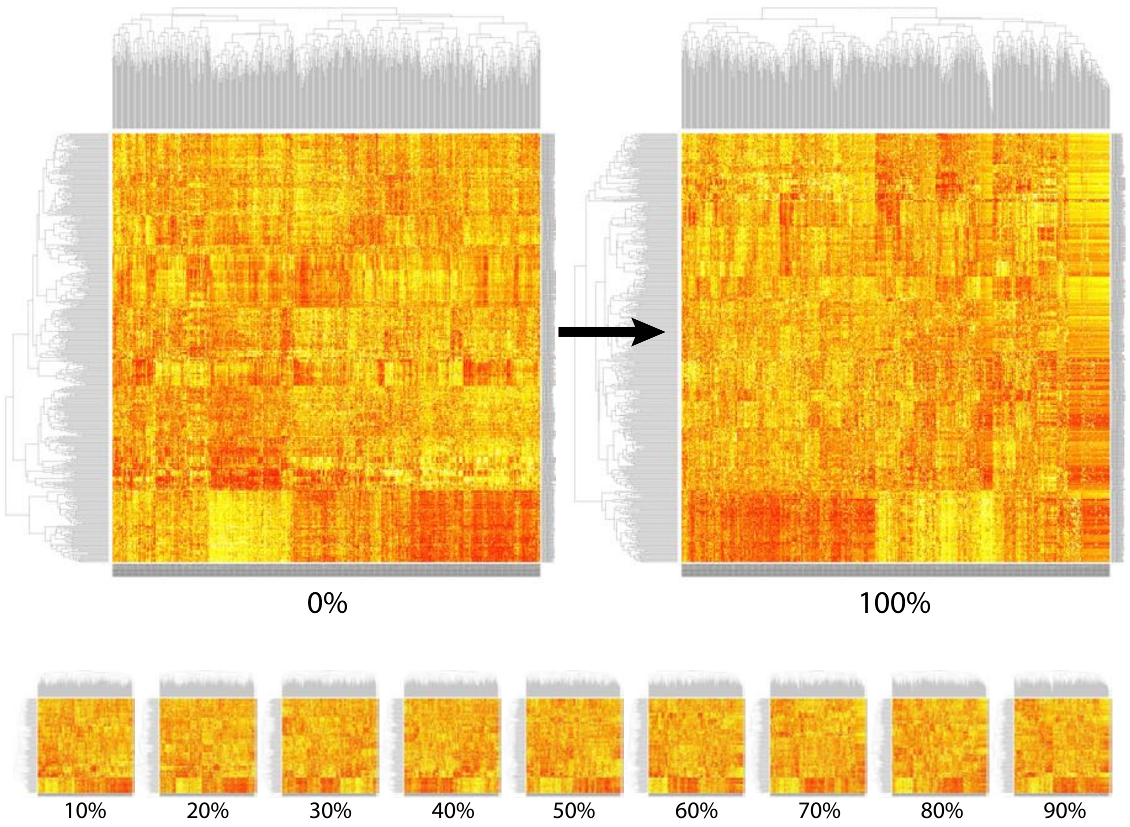


**MIGs segregate tumours from benign tissue**. 11 visual representations of the relative performance of MIGs and non-MIGs with respect to clustering gene expression data. Agglomerative MIG clustering on a normalized gene expression matrix. Gene expression values from 9 cancer types (Biliary-, Breast-, ColoRect-, Lung-, Ovary-, Panc-, Prost-, Stomach-, Thy-adenocarcinoma) totaling N=444 samples were taken from PCAWG. Pre-normalized expression matrix was normalized using quantile normalization, followed by inverse quantile normalization to map to a standard normal distribution; this step also removed outlier genes. Using the normalized gene expression matrix, clustering was performed by using hierarchical clustering by employing Euclidean metric and Ward linkage. Shown are representative heatmaps of gene expression clustering for different relative abundances of MIGs and non-MIGs. The value below each heatmap represents the fraction of non-MIGs within that heatmap's associated expression matrix.

**Supplementary Fig. S1.3 (related to Fig. 1)**

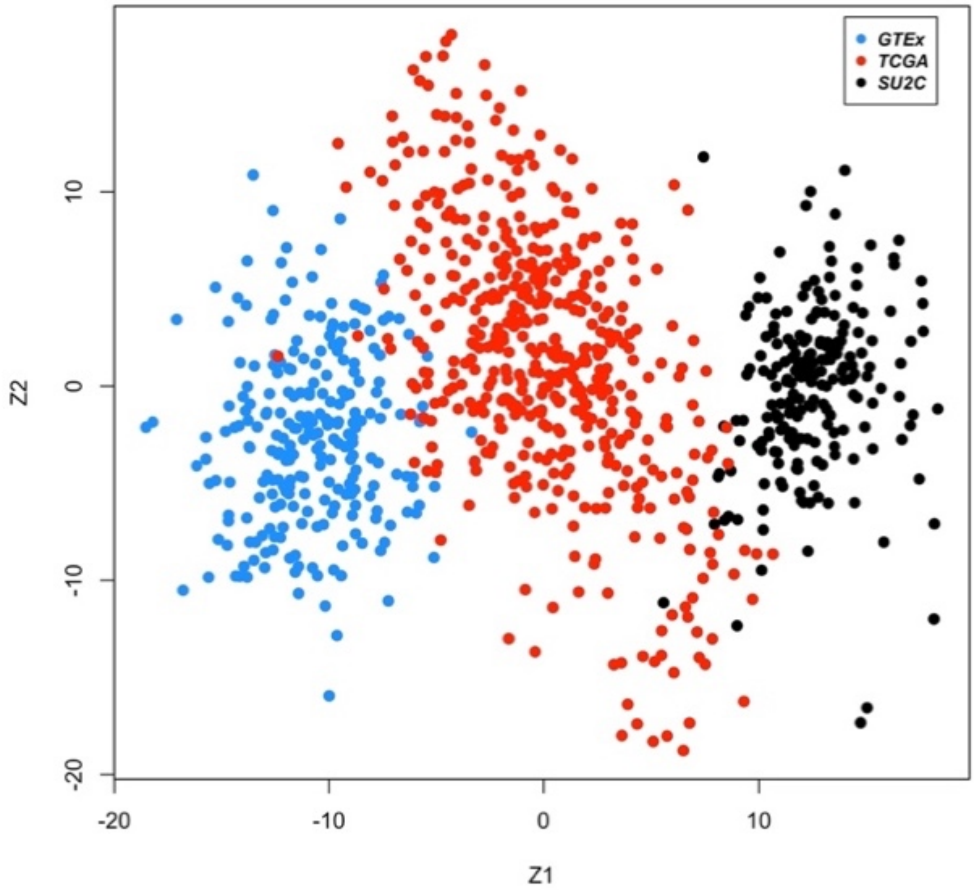


**GTEx, TCGA and SU2C datasets have distinct transcriptomes**: merged MIG expression data from 3 sources: prostate samples from GTEx (healthy prostate tissue), prostate cancer samples from TCGA (primary prostate cancer samples), and prostate cancer samples from SU2C (advanced prostate cancer). PCA analysis was carried out on the normalized gene expression matrix, thereby enabling visualization of the resultant data as the projections onto the space spanned by the first 2 PCs. Notably, this visualization appears to capture the progression from these 3 broad phases of prostate cancer progression, from healthy tissue in GTEx (at lower ends of the first PC) to advanced stages in SU2C (at higher ends of the first PC).

**Supplementary Fig. S1.5 (related to Fig. 1)**

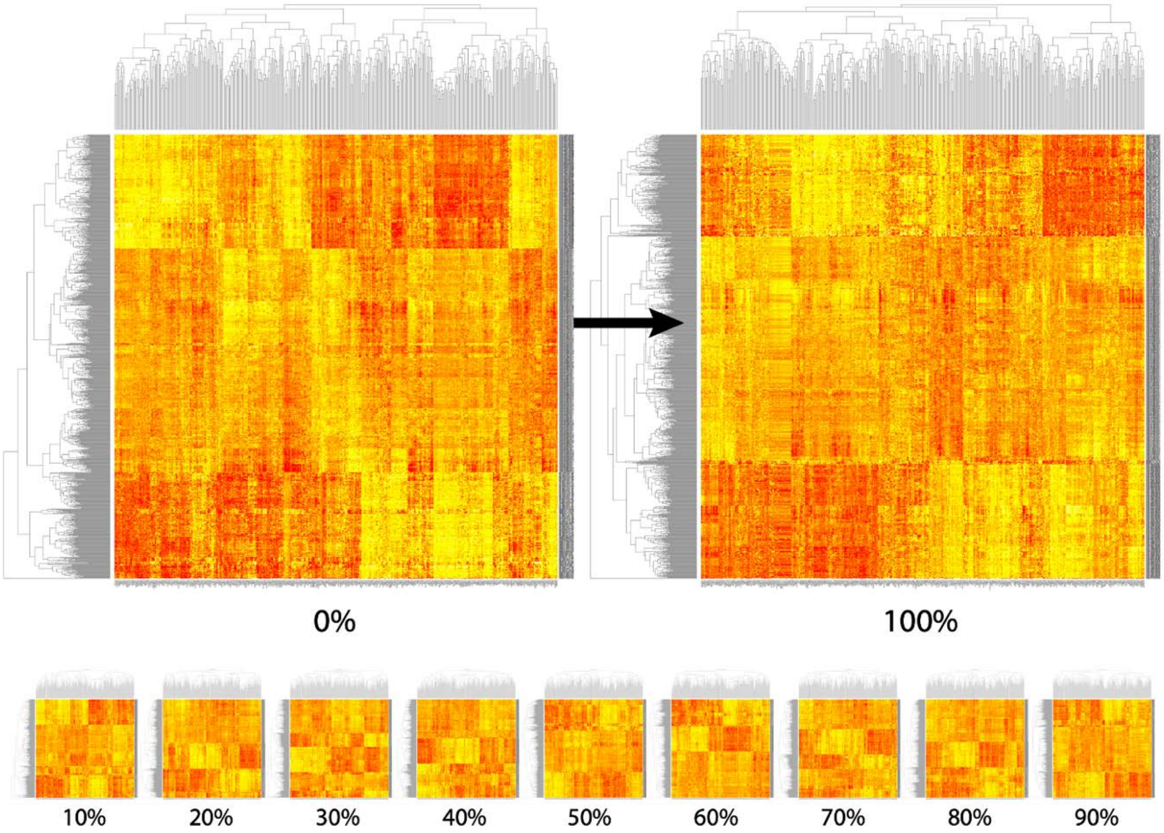


**MIGs segregate advanced PCa from benign and primary cases:** Relative performance of MIGs and non-MIGs with respect to clustering gene expression data from different stages of Prostate cancer progression. Gene expression data were taken from GTEx, TCGA, and SU2C; respectively, these datasets represent increasing stages of tumor progression (from healthy tissue in GTEx to advanced PCa tumors in SU2C). Clustering was performed using an n_cluster value of K=3. These 11 heatmaps correspond to the simulations detailed for Fig. 1F and constitute visual representations of the relative performance of MIGs and non-MIGs with respect to clustering gene expression data from these three datasets. Shown are representative heatmaps of gene expression clustering for different relative abundances of MIGs and non-MIGs. The value below each heatmap represents to the fraction of non-MIGs within that heatmap's associated expression matrix. Quantile normalization, followed by inverse quantile normalization was used to map to a standard normal distribution; this step also removed outlier genes. Using the normalized gene expression matrix, clustering was performed by using hierarchical clustering by employing Euclidean metric and Ward linkage.

**Supplementary Fig. S1.5 (related to Fig. 1)**

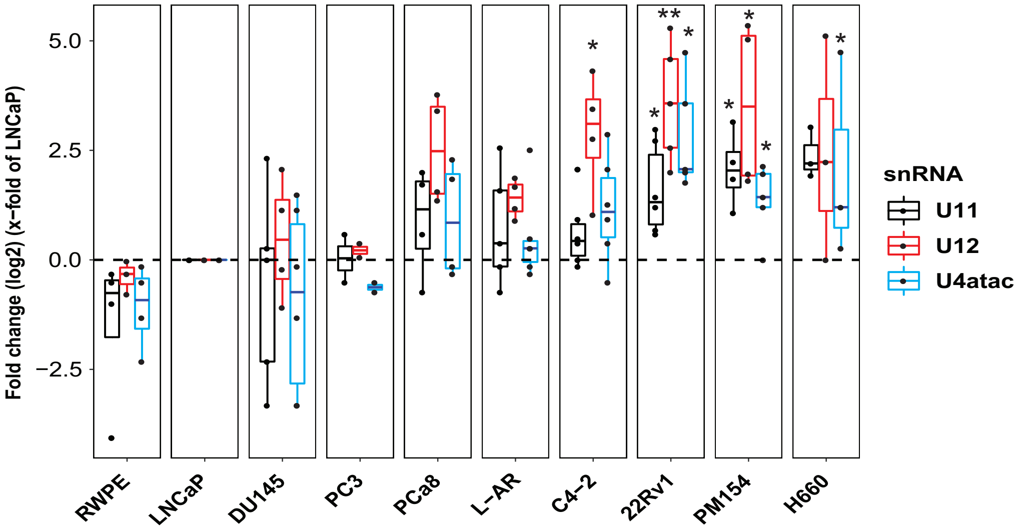


**MiS snRNA expression correlates with PCa progression**. U11, U12, U4atac and U6atac expression as x-fold of the respective LNCaP expression values normalized to the mean of GAPDH and ACTB gene transcription in PCa cell lines of different PCa subtypes (RWPE n=3, LNCaP n=3, DU145 n= 4, PC3 n=3, C4-2 n=4, 22Rv1 n=5, PM154 n=4, PCa8=4 n=7, H660=4). Data are represented using box and whisker plots. Boxplots display values of minimum, first quartile, median, third quartile, and maximum (two-sided unpaired t-test, ns $p > 0.05$, $*p < 0.05$, $**p < 0.01$). Each data point represents a single experiment, experiments were performed in triplicates.

**Supplementary Fig. S1.6 (related to Fig. 1)**

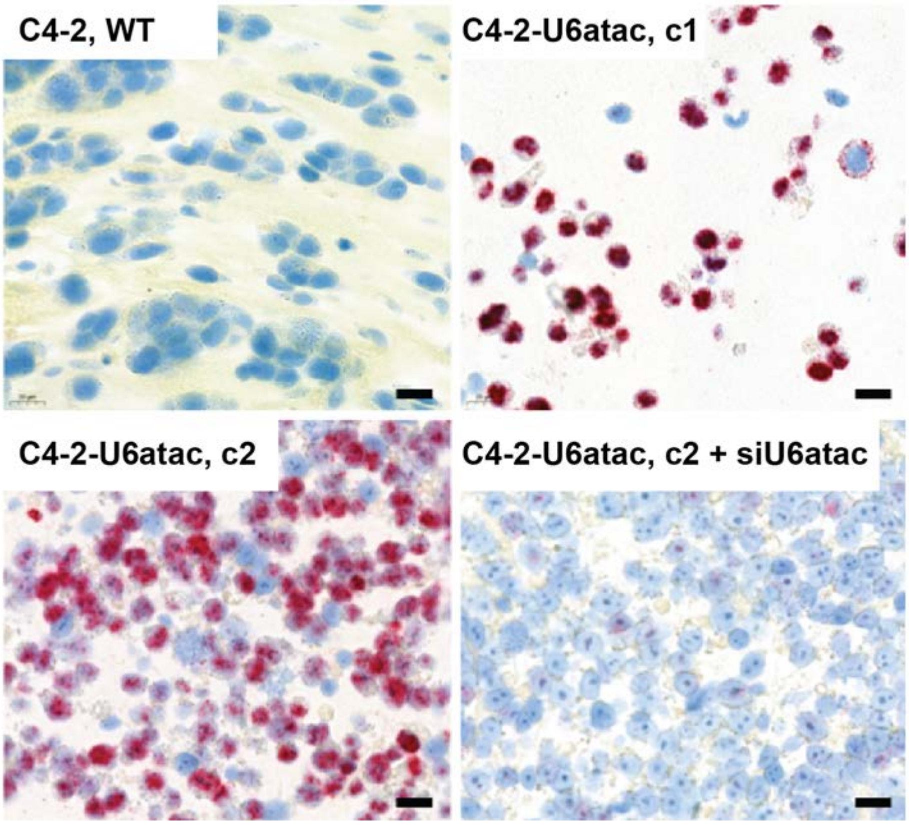


**RNU6atac probe validation**. Representative images of the validation of RNU6atac BaseScope probes in C4-2 wildtype (WT) cells, in C4-2 cells that overexpress U6atac (two different clones (c1 and c2)) and in C4-2 cells overexpressing U6atac (c2) that were treated with siU6atac RNA for 96 hours. Scale bars represent 20µm.

### Supplementary Fig. S1.7 (related to Fig. 1) – not mentioned in mss

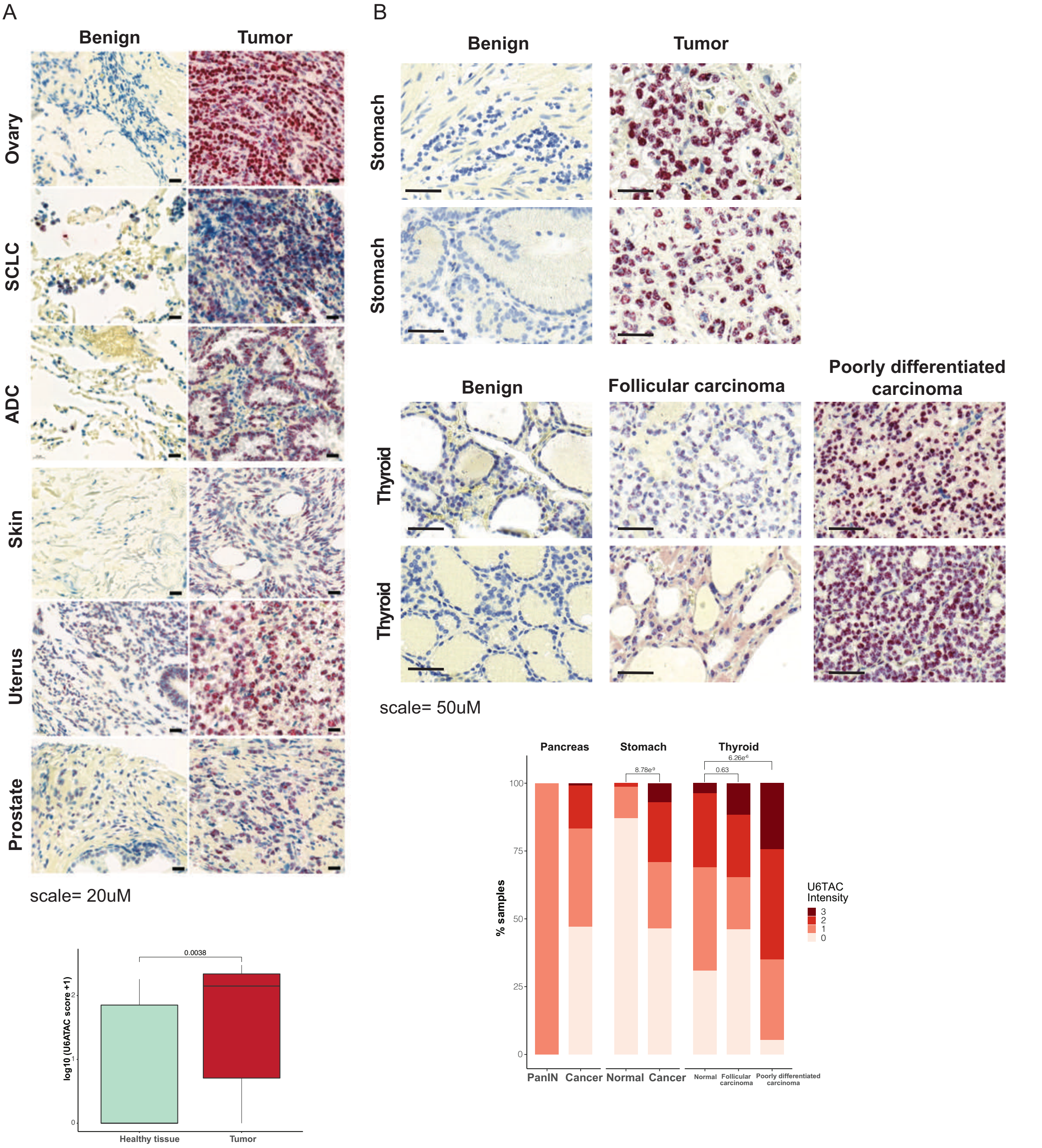


**U6atac expression is increased in cancer. A)** Representative images of RNU6atac BaseScope probe expression in human ovary, small cell lung cancer (SCLC), (ADC), skin, uterus and prostate benign (n=15) and tumor tissue (n=33). Scale bars represent 20µm. Lower panel shows the U6atac score of the TMA analysis (Wilcoxon Test, p=0.0038). **B)** Representative images of RNU6atac BaseScope probe expression in human stomach benign (n=82) and tumor (n=102) tissue and from thyroid benign tissue (n=142), thyroid follicular carcinoma (n=26) and poorly differentiated thyroid carcinoma (n=39). Scale bars represent 50µm. Lower panel quantifies the U6atac intensity in the different samples (Wilcoxon Test, stomach: p=8.78E-9, tyroid follicular carcinoma: p=0.63, thyroid poorly differentiated carcinoma: p=6.26E-6).

### Supplementary Fig. S2.1 (related to Fig. 2)

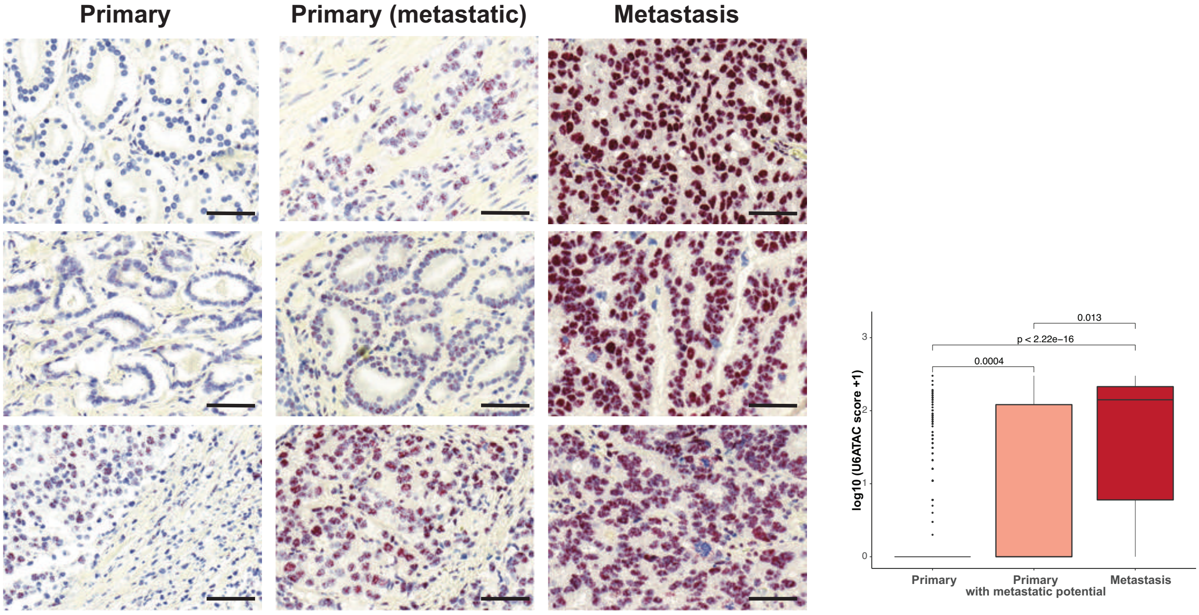


scale= 50uM

**U6atac expression is increased in metastatic PCa.** Representative images of RNU6atac BaseScope probe expression in human primary (n= 325), primary with metastatic potential (n=25) and metastatic PCa (n=32). Scale bars represent 50µm. Right panel quantifies the U6atac score across all samples tested (Wilcoxon Test, primary vs. primary with metastatic potential: p=0.0004, primary vs. metastasis: p< 2.2E-16, primary with metastatic potential vs. metastasis: p=0.013).

### Supplementary Fig. S2.2 (related to Fig. 2)

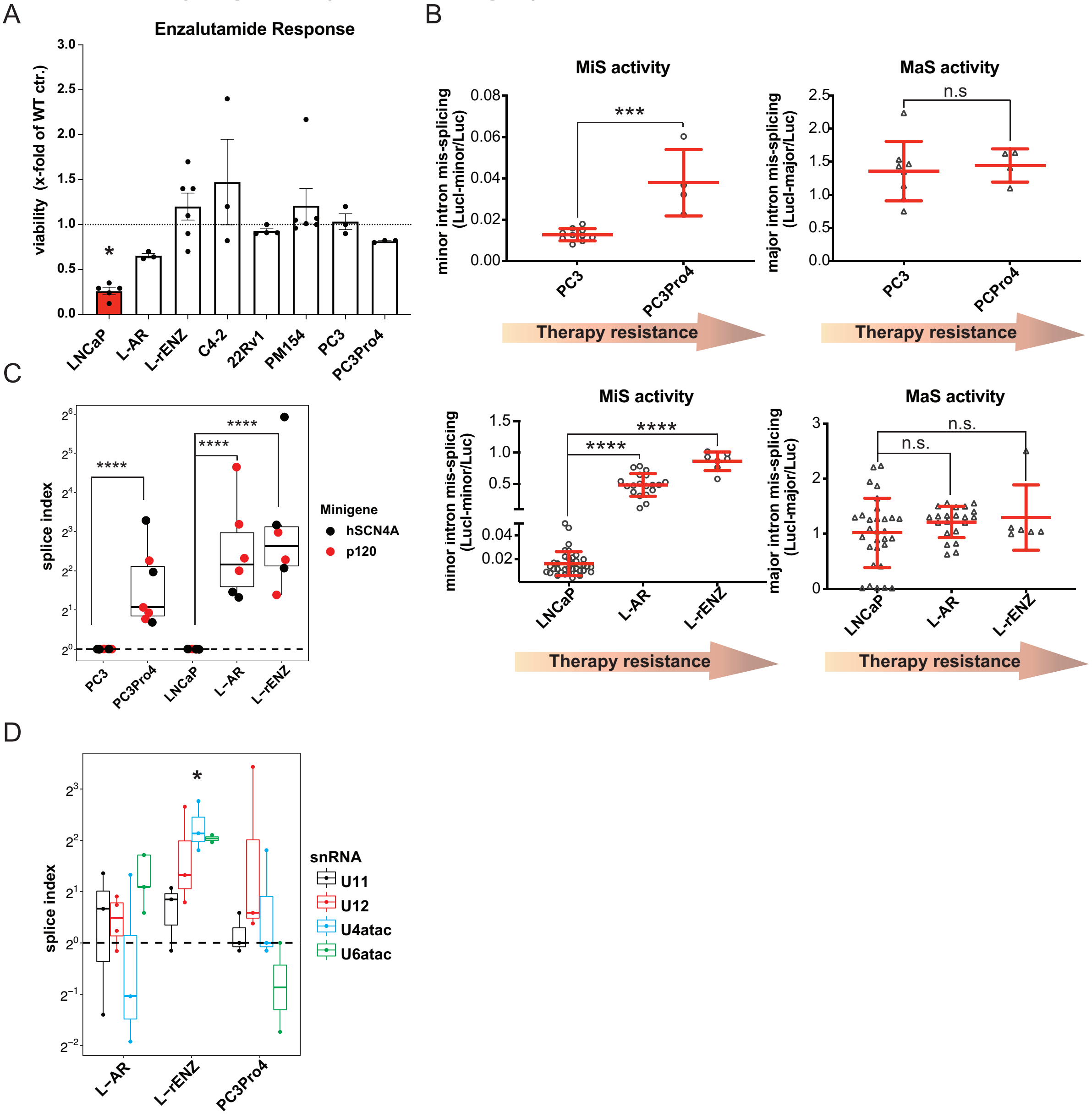


**MiS component expression and MiS activity increases with PCa progression. A)** Cell viability measured using CellTiter-GLO in human PCa cell lines treated with 20uM enzalutamide for 48 hours. Data was normalized to the untreated WT control (ctr.) (LNCaP n=6, L-AR n=3, L-rENZ n=6, C4-2 n=3, 22Rv1 n=4, PM154 n=6, PC3 n=3, PC3Pro4 n=3, mean ± SEM, ordinary one-way Anova; ns $p > 0.05$, $*p < 0.05$). Experiments were performed in triplicates. **B)** Normalized luminescence values of minor/major spliceosome luc-reporter plasmids in PC3 cells (n=10) and its subline PC3Pro4 (n=4) (upper row) and LNCaP cells (n=37), its AR overexpressing subline L-AR (n=19) and enzalutamide resistant subline L-rENZ (n=7, mean ± SD, ordinary one-way Anova; ns $p > 0.05$, $***p$, $****p < 0.0001$). Each data point represents a single experiment, experiments were performed in triplicates. **C)** hSCN4A and p120 reporter plasmid splice index in PC3, PC3Pro4, LNCaP and L-rENZ cells (n=3, two-sided unpaired t-test, $****p < 0.0001$). Splice index was calculated as the ratio of hSCN4A and p120 spliced vs. unspliced expression values of cells transfected with the hSCN4A, and p120 minigene reporter plasmids for 72h. Expression values were normalized to the mean of GAPDH and ACTB gene transcription and calculated as x-fold of siScrambled (siScr.). **D)** U11, U12, U4atac and U6atac expression as x-fold of the respective LNCaP and PC3 expression values normalized to the mean of GAPDH and ACTB gene transcription in L-rENZ (n=3) and

PC3Pro4 cells (n=3). Data are represented using box and whisker plots. Boxplots display values of minimum, first quartile, median, third quartile, and maximum (ordinary one-way Anova, ns $p > 0.05$, $*p < 0.05$). Each data point represents a single experiment, experiments were performed in triplicates.

**Supplementary Fig. S2.3 (related to Fig. 2)**

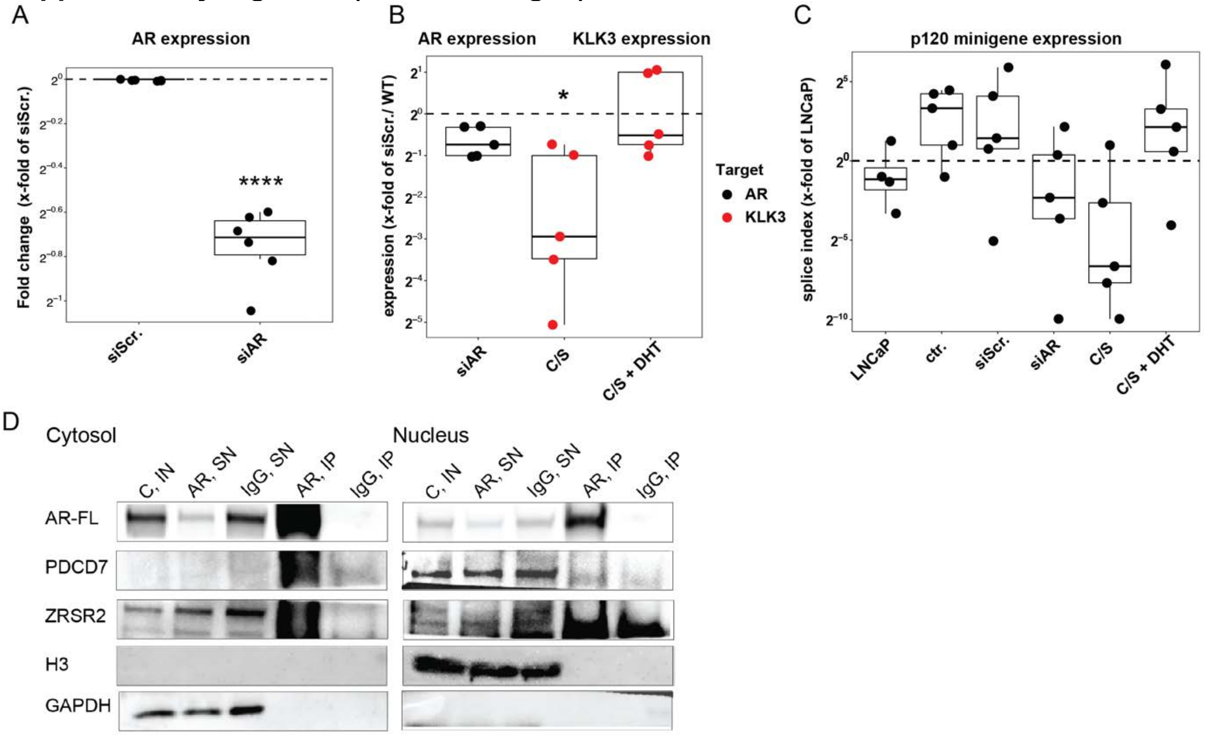


**AR regulates and interacts with the MiS. A)** AR expression as x-fold of siScrambled (siScr.) normalized to the mean of GAPDH and ACTB gene transcription in L-AR cells (n=6) treated for 96h with siRNA against AR (siAR). Data are represented using box and whisker plots. Boxplots display values of minimum, first quartile, median, third quartile, and maximum (two-sided unpaired t-test, $****p < 0.0001$). Each data point represents a single experiment, experiments were performed in triplicates. **B)** AR and KLK3 expression as x-fold of siScrambled (siScr.) normalized to the mean of GAPDH and ACTB gene transcription in L-AR cells (n=5) treated for 96h with siAR, 48h with charcoal stripped media (C/S) or with C7S media (48h) and dihydrotestosterone (DHT, 10nM, 24h). Two-sided unpaired t-test, $*p < 0.05$). Each data point represents a single experiment, experiments were performed in triplicates. **C)** U6atac expression as x-fold of LNCaP WT cells normalized to the mean of GAPDH and ACTB gene transcription in LNCaP, L-rENZ and L-AR cells treated with siAR (72h), C/S media (48h) and C/S media + DHT (24h,10uM). Data are represented using box and whisker plots. Each data point represents a single experiment, experiments were performed in triplicates. **D)** hSCN4A and p120 reporter plasmid splice index in LNCaP (n=5) and L-AR cells (n=4, t two-sided unpaired t-test; ns $p > 0.05$) treated for siAR (96h), C/S media for 48h or with C/S media (48h) and dihydrotestosterone (DHT, 10 nM, 24h). Splice index was calculated as the ratio of hSCN4A and p120 spliced vs. unspliced expression values of cells transfected with the hSCN4A, and p120 minigene reporter plasmids for 72h. Expression values were normalized to the mean of GAPDH and ACTB gene transcription and calculated as ratio of spliced vs. unspliced transcript. **E)** Co-IP of AR-FL followed by immunoblotting for PDCD7, ZRSR2, H3 and GAPDH in cytosolic and nuclear extracts from L-AR cells. Abbreviations: IN= Input, SN= supernatant, IP = Immunoprecipitation. linked PLA secondary probes [mouse, M(+) and rabbit, R(-)]. Experiments were repeated 3 times.

**Supplementary Fig. S2.4 (related to Fig. 2)**

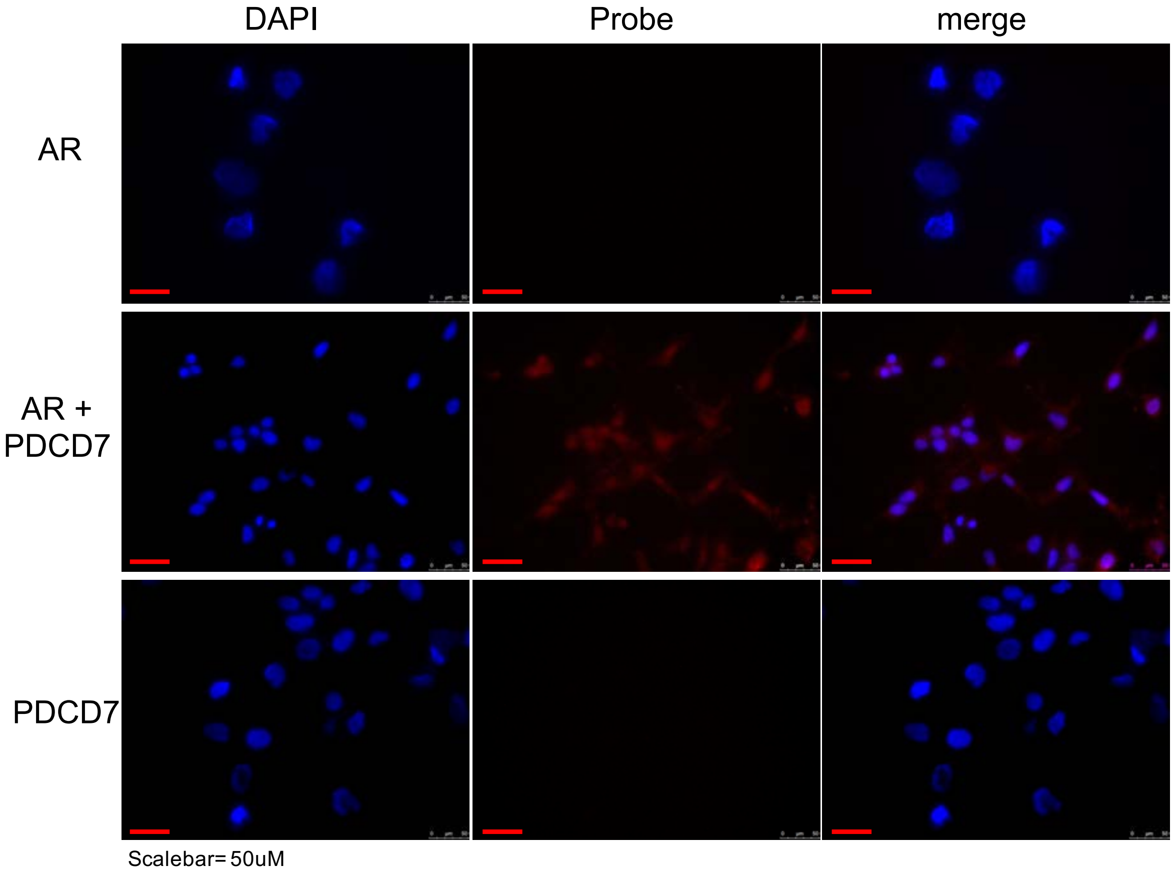


**AR interacts with the MiS protein PDCD7.** Representative 20x confocal photomicrographs of PLA signal from AR and PDCD7 heteromeric formation (red puncta). PLA was performed using anti-PDCD7 antibody (rabbit monoclonal) and Androgene receptor antibody (mouse monoclonal) primary antibody and oligonucleotide-linked PLA secondary probes [mouse, M(+) and rabbit, R(-)]. Scale bars represent 50 μm. Experiments were repeated 3 times.

### Supplementary Fig. S3.1 (related to Fig. 3)

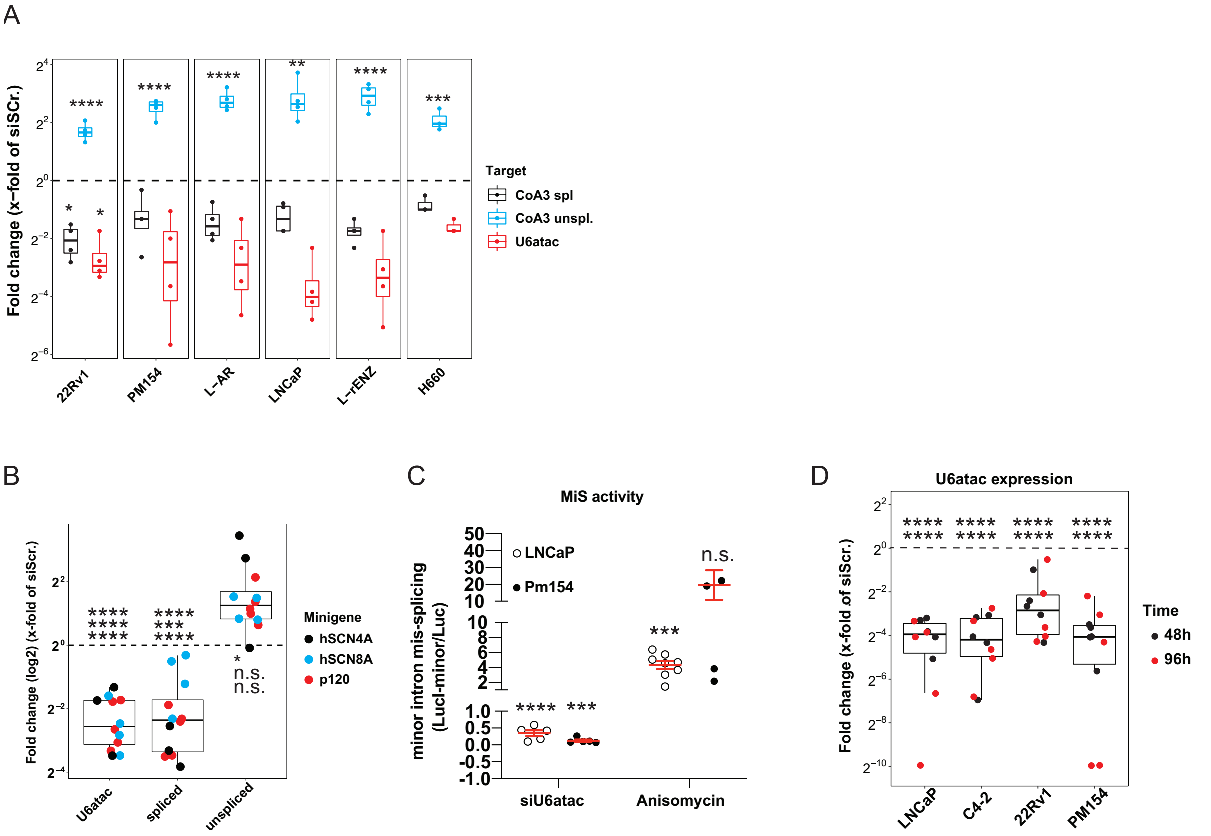


**siU6atac RNA decreases MiS activity. A)** U6atac and p120 spliced and unspliced expression as x-fold of siScrambled (siScr.) normalized to the mean of GAPDH and ACTB gene transcription in 22Rv1, PM154, L-AR, LNCaP, L-rENZ and H660 cells treated with siU6atac RNA for 96 hours. Data are represented using box and whisker plots. Boxplots display values of minimum, first quartile, median, third quartile, and maximum (n=4, ordinary one-way ANOVA, 22Rv1/U6atac: *p=0.0193, 22Rv1/CoA3 spliced: *p=0.0315, LNCaP: **p=0.0017, H660: ***p=0.0003, ****p < 0.0001). Each data point represents a single experiment, experiments were performed in triplicates. **B)** U6atac and hSCN4A, hSCN8A, p120 spliced and unspliced expression as x-fold of siScrambled (siScr.) normalized to the mean of GAPDH and ACTB gene transcription in C4-2 cells transfected with the hSCN4A, hSCN8a and p129 minigene reporter plasmids and treated with siU6atac RNA for 96 hours (hSCN4A n=3, hSCN8A n= 4, p120 n=5). Data are represented using box and whisker plots. Boxplots display values of minimum, first quartile, median, third quartile, and maximum (two-sided unpaired t-test, ns p > 0.05, ***p < 0.001, ****p < 0.0001). Each data point represents a single experiment, experiments were performed in triplicates. **C)** Normalized luminescence values of minor spliceosome luc-reporter plasmids in LNCaP (siU6atac n=5, Anisomycin n=8) and PM154 (siU6atac n=5, Anisomycin n=4) cells treated with siU6atac (96h) or Anisomycin (1ug/ml for 4h) (mean ± SEM, ordinary one-way Anova; ns p > 0.05, ***p < 0.001, ****p< 0.0001). Each data point represents a single experiment, experiments were performed in triplicates. **D)** U6atac snRNA expression as x-fold of siScrambled (siScr.) normalized to the mean of GAPDH and ACTB gene transcription in LNCaP, C4-2, 22Rv1 and PM154 cells treated with siU6atac RNA for 48 and 96 hours (n=5) Data are represented using box and whisker plots. Boxplots display values of minimum, first quartile, median, third quartile, and maximum (one-way Anova; ****p < 0.0001). Experiments were performed in triplicates.

**Supplementary Fig. S3.2 (related to Fig. 3)**

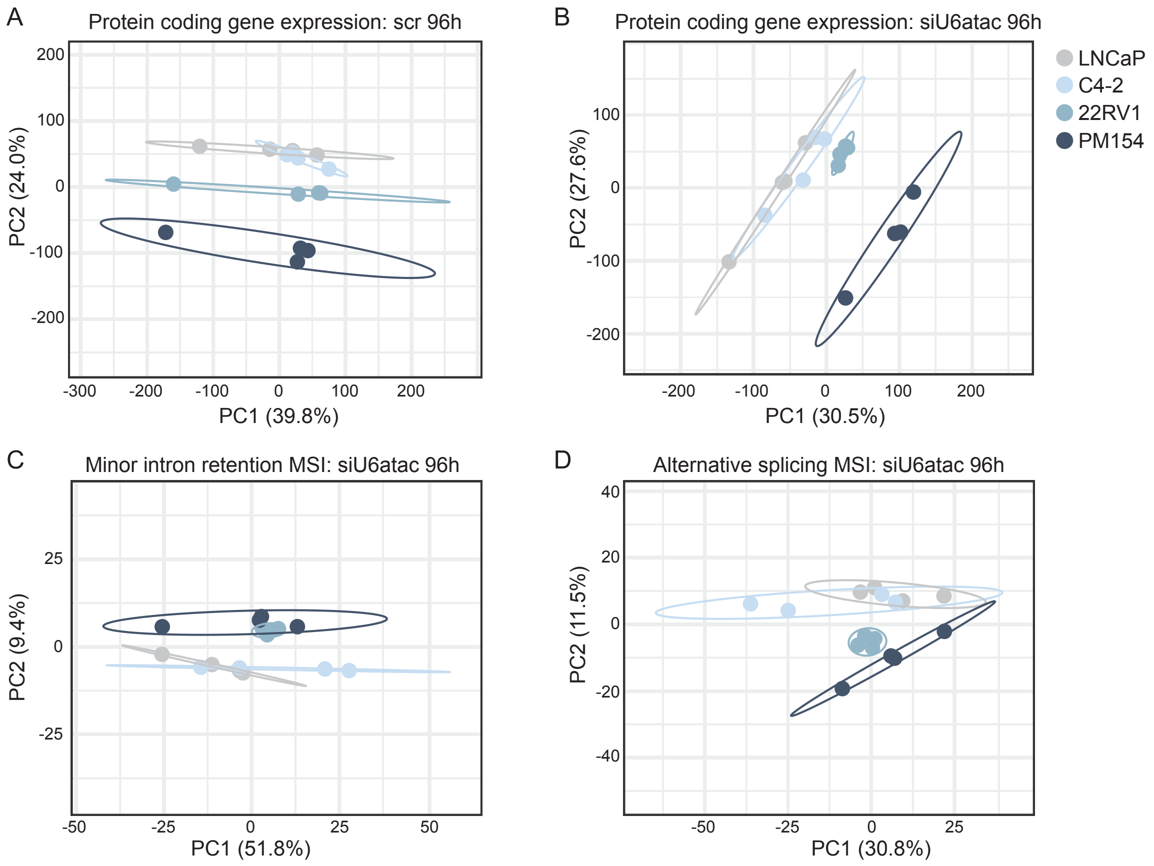


**Response to MiS inhibition in C4-2 is closer to LNCaP, whereas 22Rv1 is closer to PM154 A-B)** Principal component analysis (PCA) of log10 transformed transcript per million (TPM) values for all expressed protein coding genes in 96h **(A)** siScr or **(B)** siU6atac-treated LNCaP, C4-2, 22RV1, and PM154 cells. **(C-D)** PCA for mis-splicing index (MSI) values for all minor introns that pass filtering criteria for **(C)** minor intron retention or **(D)** alternative splicing in 96h siU6atac-treated LNCaP, C4-2, 22RV1, and PM154 cells. Ellipses show 95% confidence intervals.

**Supplementary Fig. S3.3 (related to Fig. 3)**

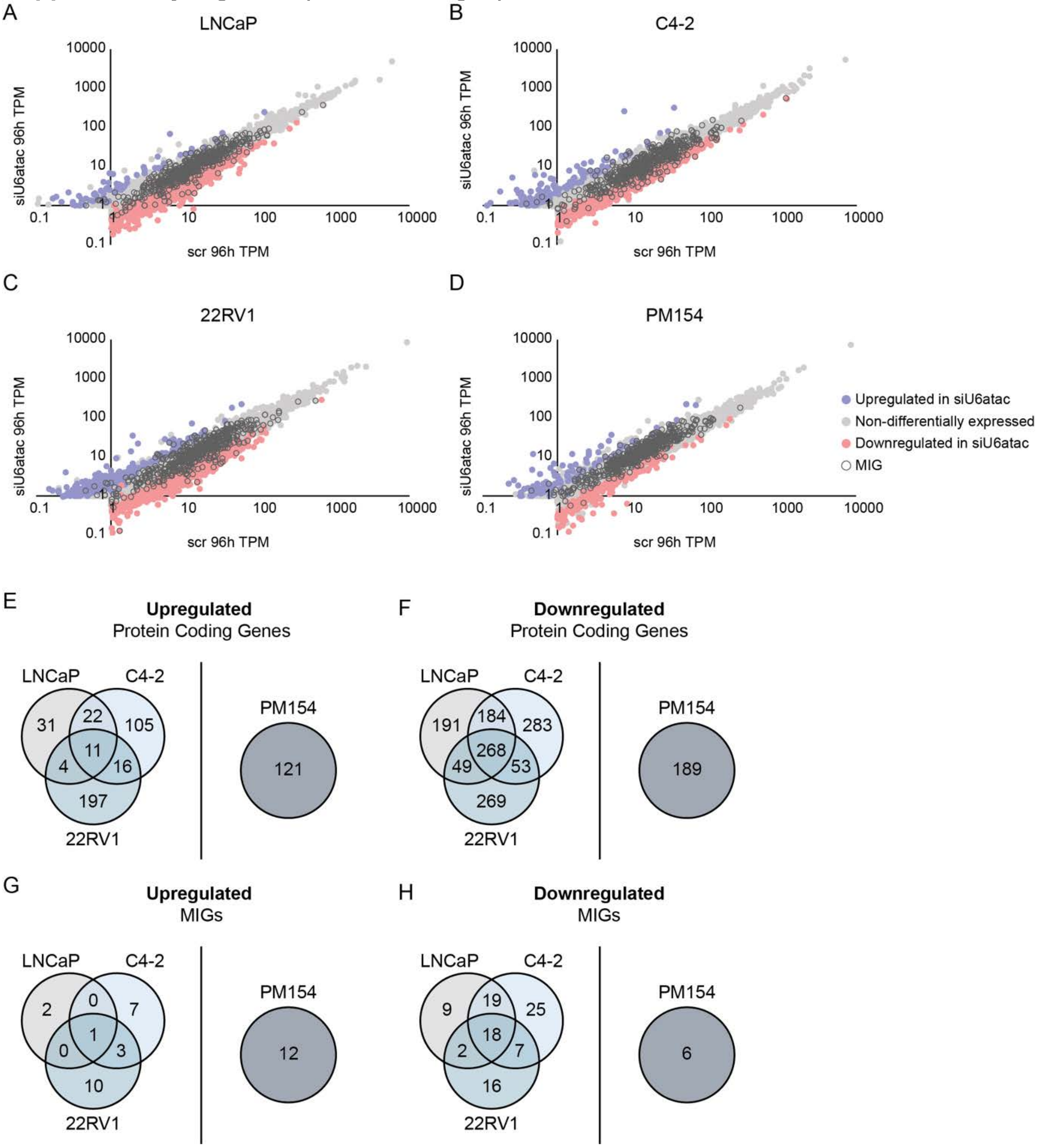


**siU6atac changes global transcriptome in PCa.. A-D)** Scatter plots showing expression of all protein coding genes with transcript per million (TPM) values ≥ 1 in 96h treated siScr (Y-axis) or siU6atac (X-axis) (**A**) LNCaP cells, (**B**) C4-2 cells, (**C**) 22RV1 cells, and (**D**) PM154 cells. Purple dots show genes significantly upregulated (log2 fold-change (FC) ≥ 1, *P* ≤ 0.01) in siU6atac treated cells; red dots show genes significantly downregulated (log2FC ≤ -1, *P* ≤ 0.01) in siU6atac treated cells; and open dark grey circles show minor intron containing genes (MIGs). **E-H**) Overlap (left) and total number (right) of (**E**) upregulated protein coding genes, (**F**) downregulated protein coding genes, (**G**) upregulated MIGs, and (**H**) downregulated MIGs in the LNCaP, C4-2, and 22RV1 cell lines and PM154 organoid, respectively.

**Supplementary Fig. S3.4 (related to Fig. 3)**

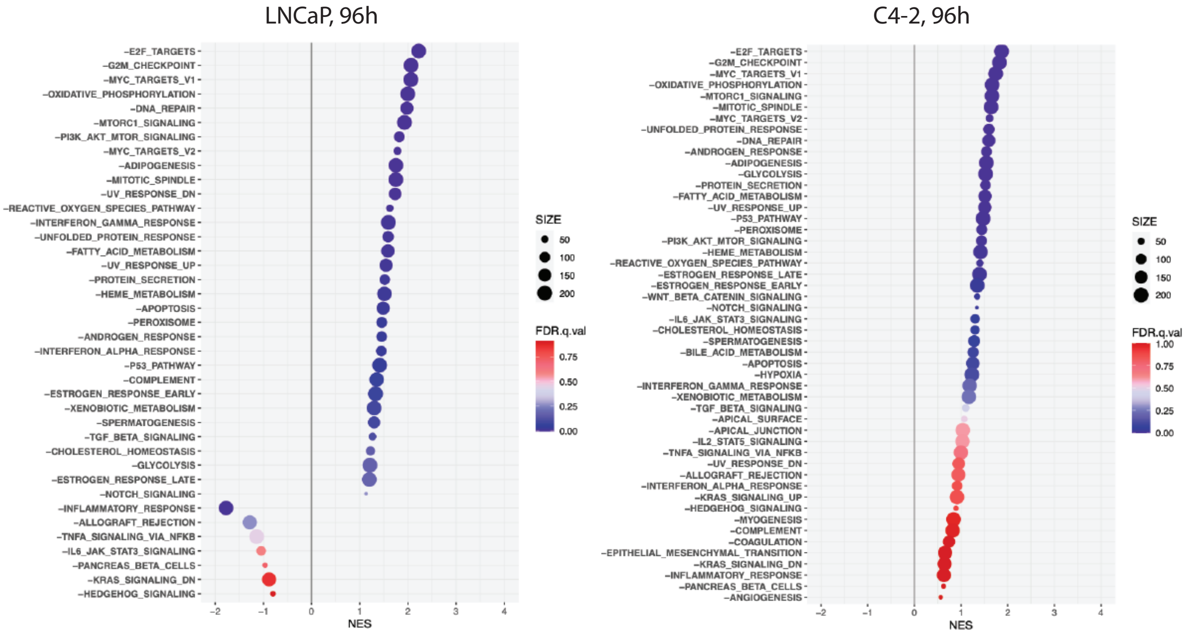


**GSEA analysis of U6atac KD cells.** Dot plot of GSEA (hallmark gene set) of RNAseq data of LNCaP and C4-2, cells treated with siU6atac (96h) (compared to the Scrambled siRNA control). Analysis was performed for all downregulated transcripts.

**Supplementary Fig. S4.1 (related to Fig. 4)**

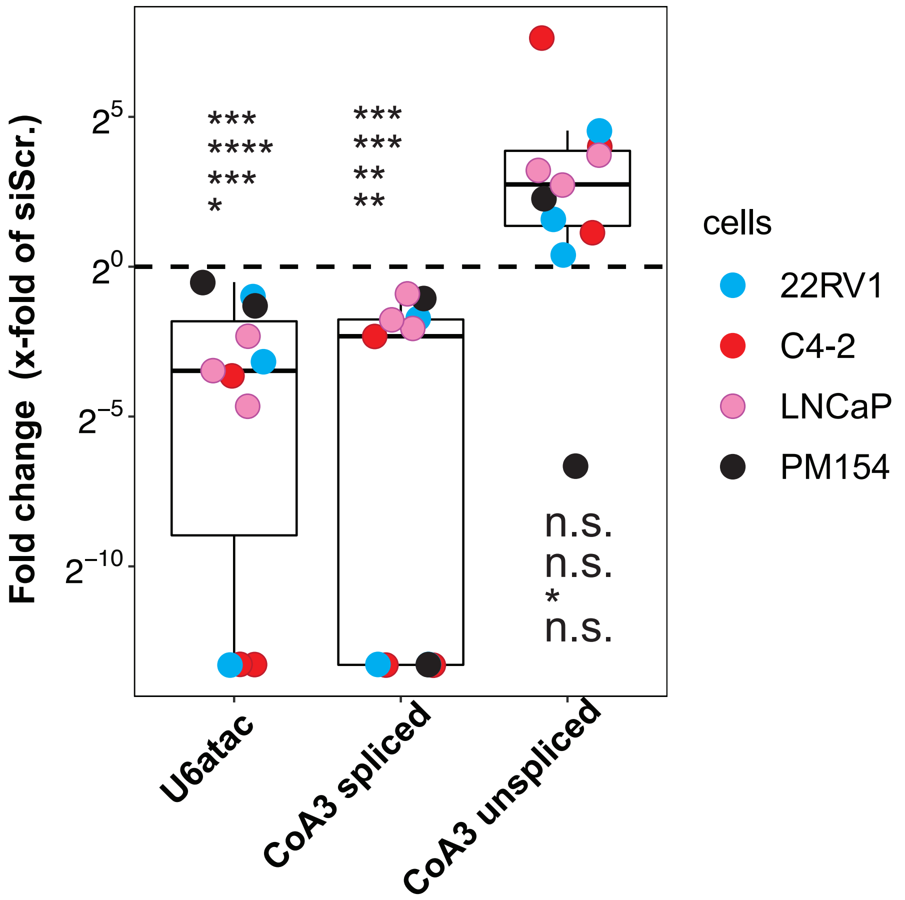


**siU6atac decreases U6atac and spliced CoA3 transcript expression.** U6atac snRNA, CoA3 spliced and unspliced expression as x-fold of siScrambled (siScr.) normalized to the mean of GAPDH and ACTB gene transcription in LNCaP, C4-2, 22Rv1 cells and PM154 organoids treated with siU6atac RNA for 96 hours (n=3) Data are represented using box and whisker plots. Boxplots display values of minimum, first quartile, median, third quartile, and maximum (ordinary one-way Anova; 6Uatac: $*$p =0.0294, $***$p =0.004 (22Rv1), 0.0002 (LNCaP, $****$p < 0.0001; CoA3 spliced: $***$p =0.0025 (PM154), 0.0035 (22Rv1), $***$p =0.0003)

**Supplementary Fig. S4.2 (related to Fig. 4)**

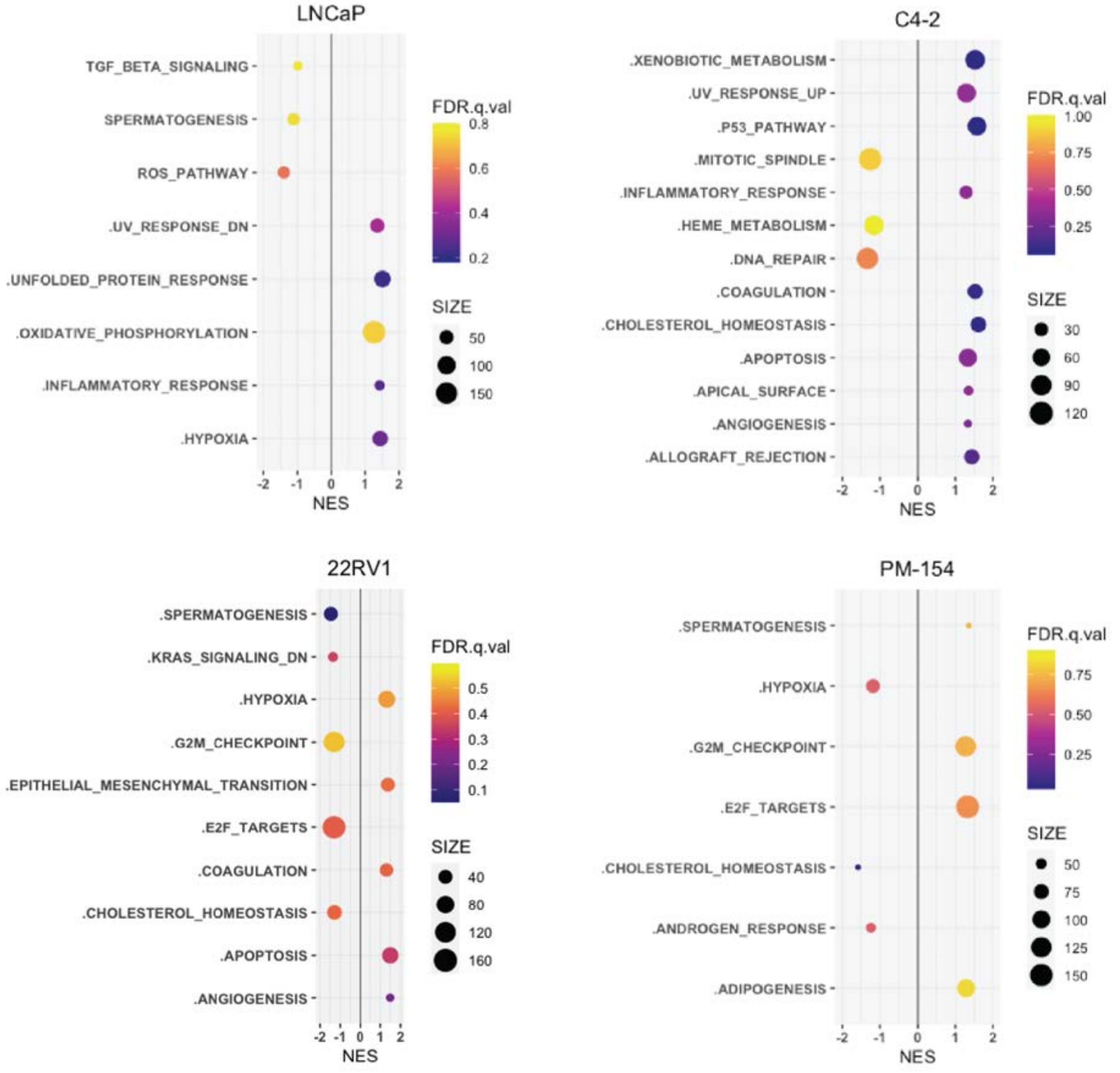


**Molecular Pathways increased/decreased after siU6atac.** Dot plot of GSEA (hallmark gene set) of decreased (left) and increased (right) proteins in LNCaP, C4-2, 22Rv1 and PM154 cells treated with siU6atac (96h) (compared to the Scrambled siRNA control).

**Supplementary Fig. S4.3 (related to Fig. 4)**

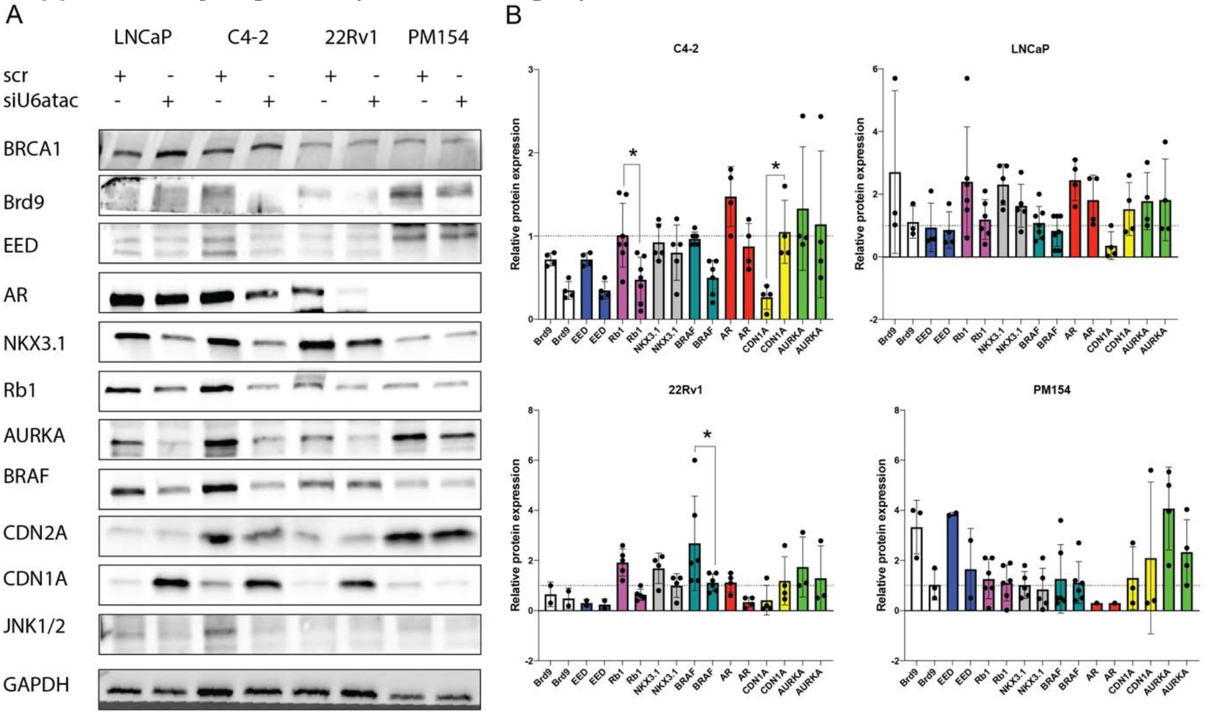


**siU6atac impacts protein expression in PCa. A)** BRCA1, Brd9, EED, AR, NKX3.1, Rb1, AURKA, BRAF, CDN2A, CDN1A and JNK1/2 Immunoblot of LNCaP, C4-2, 22Rv1cells and PM154 organoids treated for 96h with siU6atac or siScrambled (siScr.) **B)** Immunoblot quantifications. Values were normalized to GAPDH *(*ordinary one-way Anova, C4-2*:* ∗p =0.0485 (Rb1), 0.0196 (CDN1A); 22Rv1: ∗p =0.0230).

**Supplementary Fig. S4.4 (related to Fig. 4)**

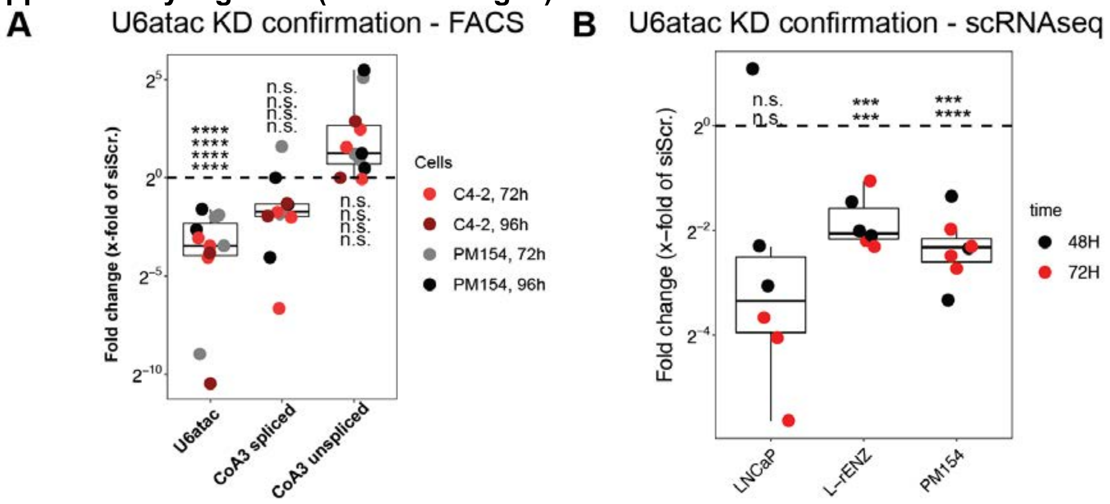


**siU6atac decreases U6atac expression and triggers a G1/G0 arrest. A)** U6atac snRNA, CoA3 spliced and unspliced expression as x-fold of siScrambled (siScr.) normalized to the mean of GAPDH and ACTB gene transcription in C4-2 cells and PM154 organoids treated with siU6atac RNA for 72h and 96 hours (n=3) Data are represented using box and whisker plots. Boxplots display values of minimum, first quartile, median, third quartile, and maximum (ordinary one-way Anova; U6atac: ****$p < 0.0001$, CoA3 spliced and unspliced: ns $p > 0.05$). **B)** U6atac snRNA as x-fold of siScrambled (siScr.) normalized to the mean of GAPDH and ACTB gene transcription in LNCaP, L-rEnz cells and PM154 organoids treated with siU6atac RNA for 48h and 96 hours (n=3) Data are represented using box and whisker plots. Boxplots display values of minimum, first quartile, median, third quartile, and maximum (ordinary one-way Anova; PM154: ****$p = 0.0001$(48h), ****$p < 0.0001$(72h), L-rENZ: ***$p = 0.0002$(48h and 72h); LNCaP: ns $p > 0.05$).

**Supplementary Fig. S5.1 (related to Fig. 5)**

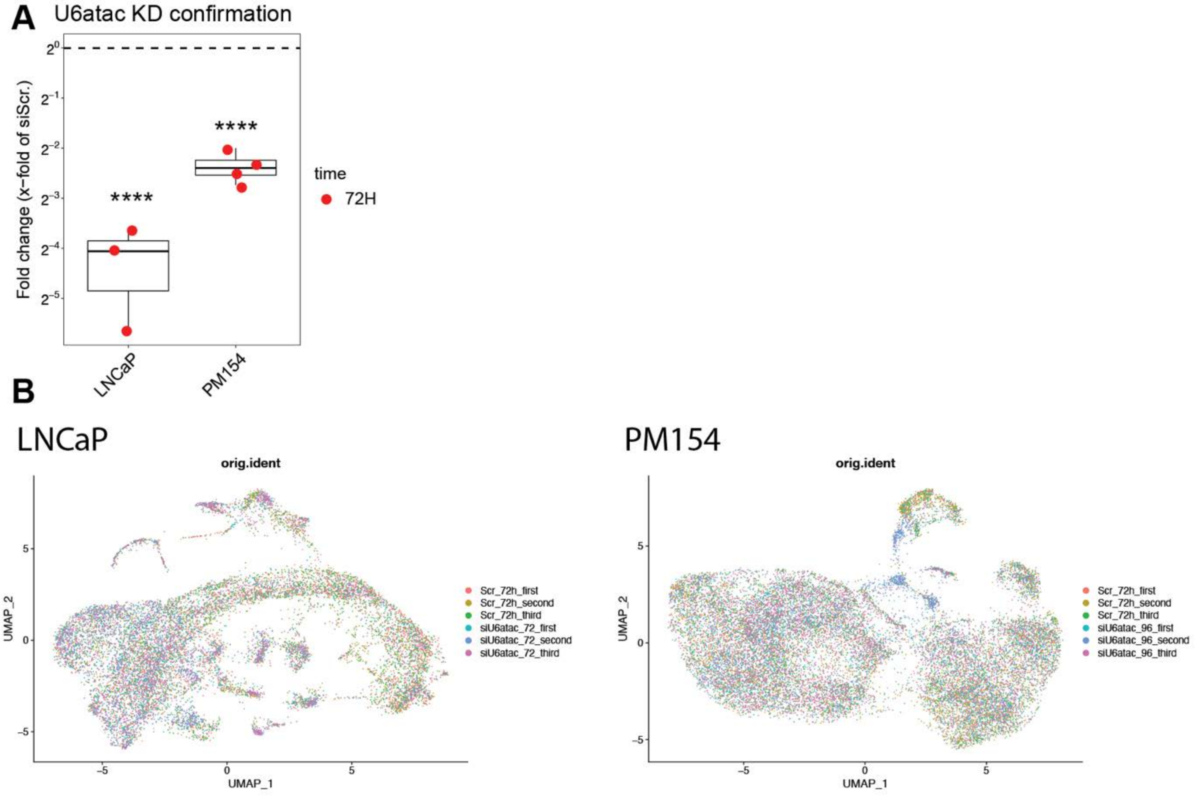


**scRNAseq metrics. A)** U6atac snRNA as x-fold of siScrambled (siScr.) normalized to the mean of GAPDH and ACTB gene transcription in LNCaP cells and PM154 organoids treated with siU6atac RNA for 72 hours (n=3). Data are represented using box and whisker plots. Boxplots display values of minimum, first quartile, median, third quartile, and maximum (ordinary one-way Anova, ∗∗∗∗p < 0.001) **B)** UMAP representation showing the contribution of each sample in each cluster.

**Supplementary Fig. S5.2 (related to Fig. 5)**

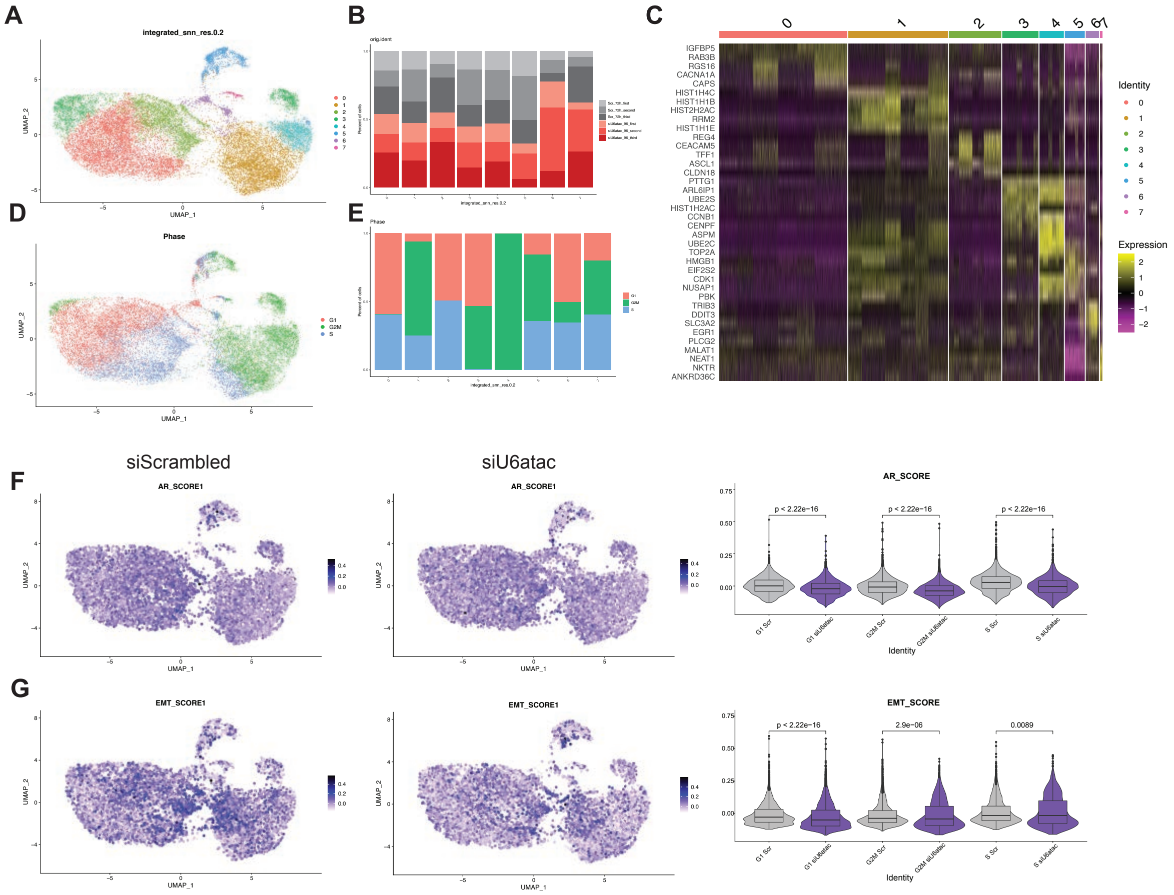


**H**

Bulk RNAseq

| Cell lines | AR score |
|---|---|
| LNCaP_scr_96 | 0.849831928 |
| LNCaP_siU6atac_96 | 0.832368982 |
| PM154_scr_96 | 0.148353093 |
| PM154_siU6atac_96 | 0.110428527 |

**scRNAseq in PM154 organoids A)** UMAP representation of PM154 cell line showing the clusters at the optimal resolution (0.2). **B)** Histogram of PM154 cell line showing the contribution of each sample in each cluster. **C)** Heatmap of LNCaP cell line showing the expression of top 5 marker genes in each cluster. **D)** UMAP representation of LNCaP cell line showing the cell cycle phase for each cell. **E)** Histogram of LNCaP cell line showing the percentage of cell cycle phase in each cluster. **F)** UMAP representation of PM154 cell line showing the AR score calculated on siScrambled and siU6atac samples. Violin representations of PM154 cell line showing the AR score calculated for each cell cycle phase. The pvalue was calculated using a Wilcoxon test. **G)** UMAP representation of PM154 cell line showing the EMT score calculated on siScrambled and siU6atac samples. Violin representations are showing the EMT score calculated for each cell cycle phase. The pvalue was calculated using a Wilcoxon test. **H)** AR score of LNCaP and PM154 cell lines calculated on siScrambled and siU6atac samples using bulk RNAseq.

### Supplementary Fig. S5.3 (related to Fig. 5)

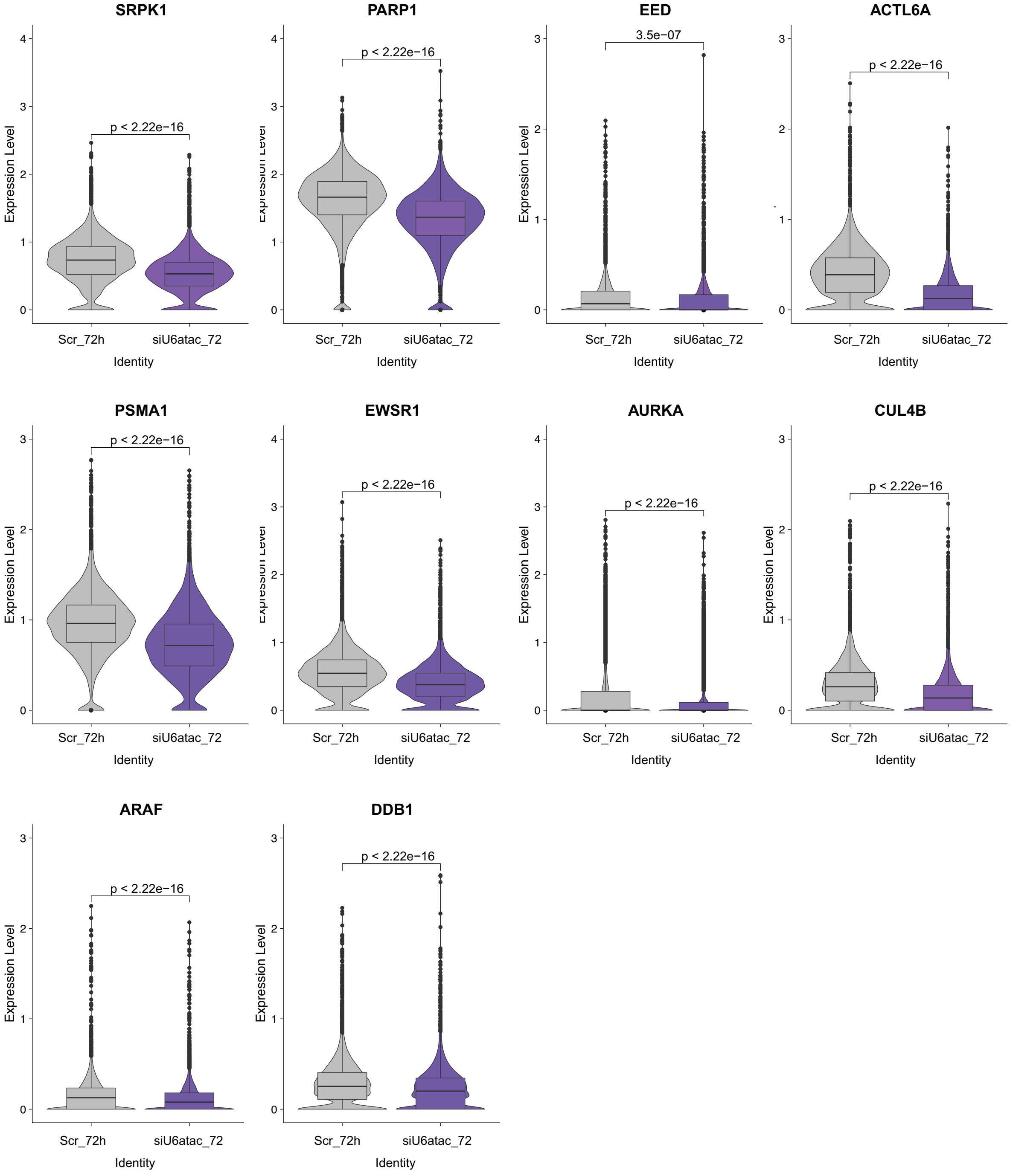


**scRNAseq analysis of MIG expression in LNCaP cells** Violin plots showing the expression levels of selected MIGs, which represent critical PCa nodes (Fig. 1B, C) in LNCaP cells.

### Supplementary Fig. S5.4 (related to Fig. 5)

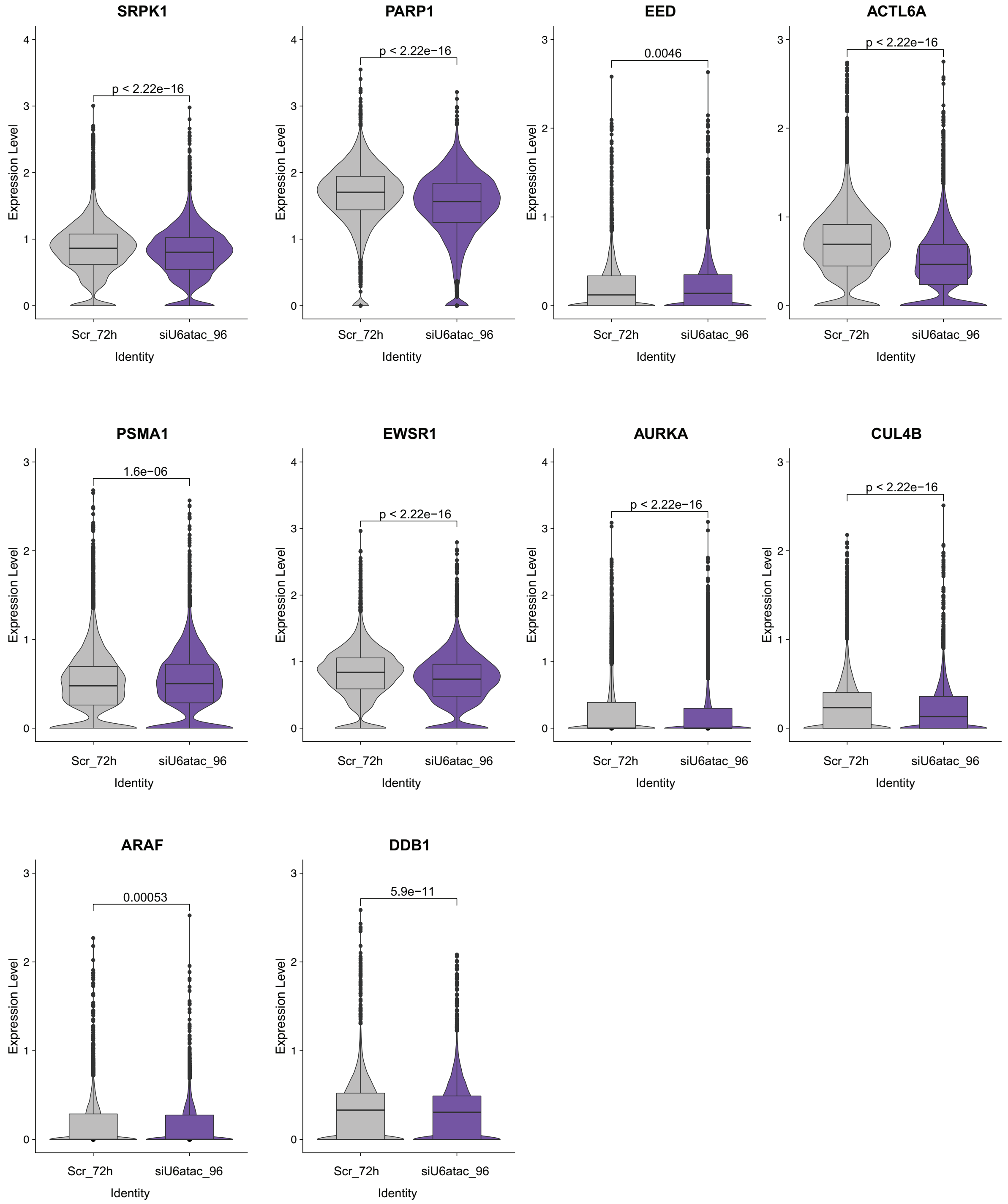


**scRNAseq analysis of MIG expression in PM54 organoids.** Violin plots showing the expression levels, which represent critical PCa nodes (Fig. 1B, C) in PM154 cells.

**Supplementary Fig. S6.1 (related to Fig. 6)**

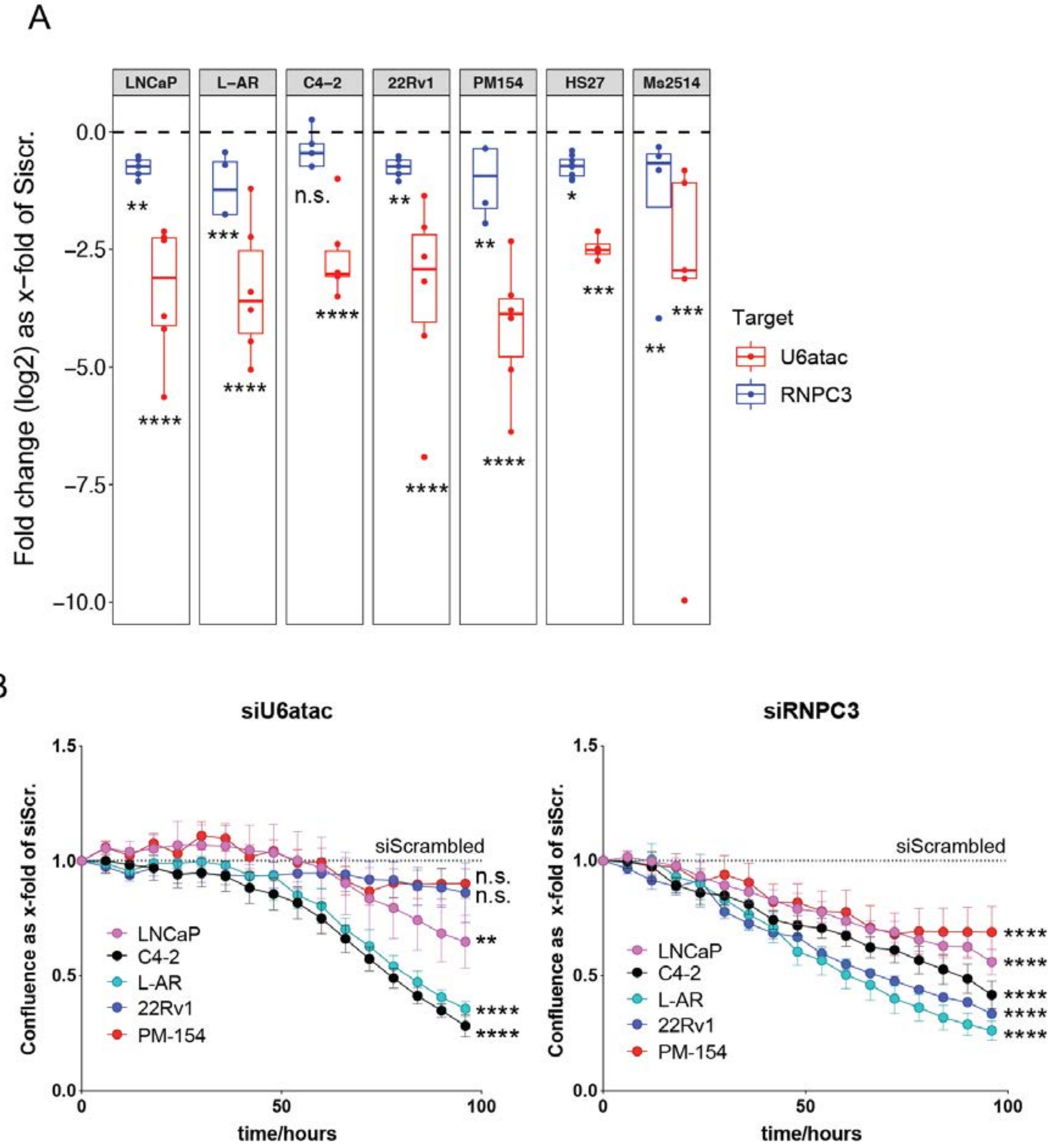


**siU6atac and siRNPC3 impact PCa cell growth. A)** U6atac snRNA and RNPC3 expression as x-fold of siScrambled (siScr.) normalized to the mean of GAPDH and ACTB gene transcription in LNCaP, L-AR, C4-2, 22Rv1, PM154, HS27 and MS2514 cells and organoids treated with siU6atac (n=5) and siRNA against RNPC3 (siRNPC3) (n=3) for 96 hours. Data are represented using box and whisker plots. Boxplots display values of minimum, first quartile, median, third quartile, and maximum (two-way Anova; LNCaP: ∗∗p =0.0076, ∗∗∗∗p < 0.00’1; L-AR: ∗∗∗p =0.0006, ∗∗∗∗p < 0.0001; C4-2: ns p > 0.05, ∗∗∗∗p < 0.0001; 22Rv1: ∗∗p=0.0048, ∗∗∗∗p < 0.0001; PM154: ∗∗p=0.0024, ∗∗∗∗p < 0.0001; HS27: ∗p=0.0190, ∗∗∗p =0.0005; Ms2514: ∗∗p=0.0025, ∗∗∗p =0.0007). **B)** Growth curves of LNCaP, C4-2, L-AR, 22Rv1 cells and PM154 organoids treated for 0, 48, 72 and 96h with siU6atac. Data is normalized to the siScrambled control and represents pooled results from 8 (siU6atac) and 6 (siRNPC3) biologically independent experiments (mean values +/-SEM, ordinary two-way Anova ; ns p > 0.05, ∗∗p < 0.01, ∗∗∗∗p < 0.001).

### Supplementary Fig. S6.2 (related to Fig. 6)

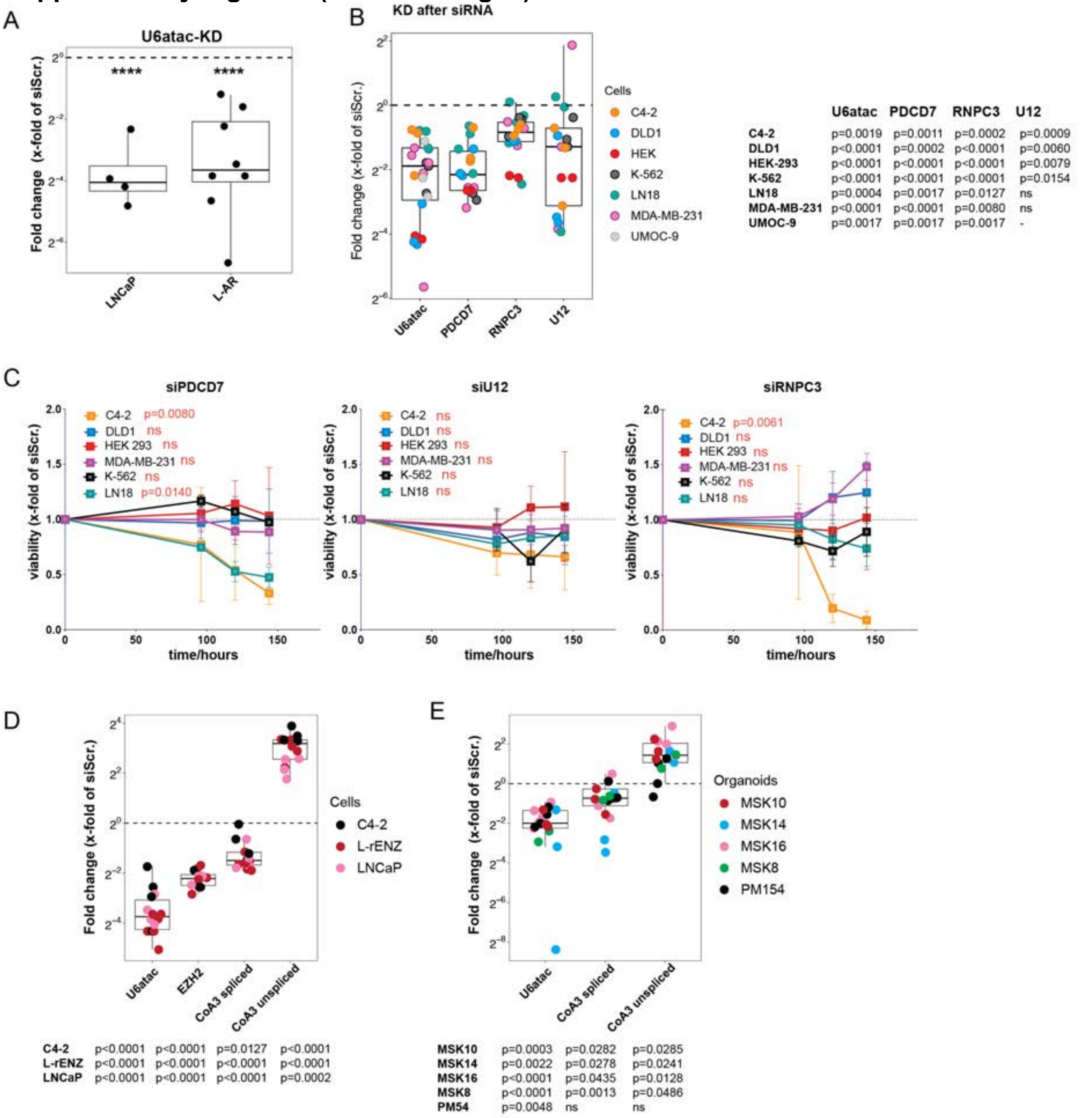


**MiS inhibition blocks viability and growth of cancer cells. A)** U6atac snRNA expression as x-fold of siScrambled (siScr.) normalized to the mean of GAPDH and ACTB gene transcription in LNCaP (n=4) and L-AR (n=8) cells treated with siU6atac for 96 hours. Data are represented using box and whisker plots. Boxplots display values of minimum, first quartile, median, third quartile, and maximum (ordinary one-way Anova, ∗∗∗∗p< 0.0001). **B)** U6atac and U12 snRNA and RNPC3 and PDCD7 expression as x-fold of siScrambled (siScr.) normalized to the mean of GAPDH and ACTB gene transcription in C4-2, DLD1, HEK 293, K-562, LN18, MDA-MB-231 and UMOC-9 cells treated with siU6atac, siPDCD7, siRNPC3 and siU12 for 96 hours (n=3). Data are represented using box and whisker plots. Boxplots display values of minimum, first quartile, median, third quartile, and maximum (unpaired two-tailed t-test; p-values indicated in figure). **C)** Cell viability assay measured using CellTiter-GLO in in C4-2, DLD1, HEK 293, K-562, LN18 and MDA-MB-231 treated cells treated with siPDCD7, siRNPC3 and siU12 (48, 72, 96 and 120h). Data is normalized to the siScrambled (siScr.) control and represents pooled results from 4 biologically independent experiments (mean values +/-SEM, two-way Anova; p-values indicated in figure) **D)** U6atac snRNA, CoA3 spliced and unspliced expression after siU6atac (96h) treatment and EZH2 expression after siEZH2 (96h) treatment as x-fold of siScrambled (siScr.) normalized to the mean of GAPDH and ACTB gene transcription in LNCaP (n=4), C4-2 (n=4) and L-rENZ (n=6) cells. Data are represented using box and whisker plots. Boxplots display values of minimum, first quartile, median, third quartile, and maximum (unpaired two-tailed t-test; p-values indicated in figure). **E)** U6atac snRNA, CoA3 spliced and unspliced expression as x-fold of siScrambled (siScr.) normalized to the mean of GAPDH and ACTB gene transcription in MSK10 (n=3), MSK14 (n=3), MSK16 (n=5), MSK8 (n=3) and PM154 (n=4) patient derived PCa organoids treated with siU6atac for 96 hours. Data are represented using box and whisker plots. Boxplots display values of minimum, first quartile, median, third quartile, and maximum (unpaired two-tailed t-test; p-values indicated in figure).

**Supplementary Fig. S6.3 (related to Fig. 6)**

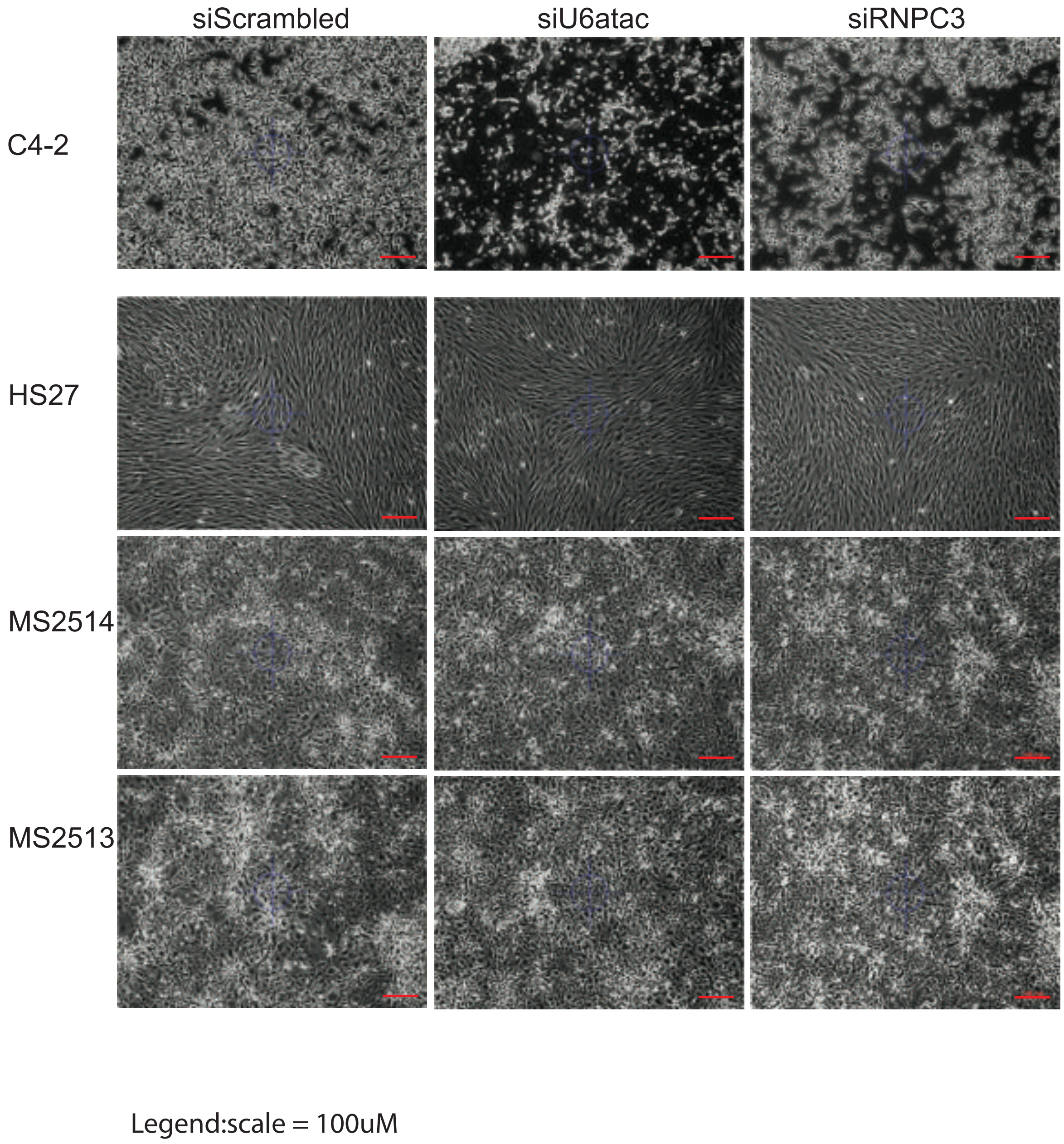


**MiS inhibition has stronger impact on cancer than on normal cells.** Brightfield microscopy of C4-2, HS27, MS2514 and MS2513 cells 96 hours after siU6atac and siRNPC3 treatment. Cells treated with Scrambled siRNA are shown as control. Scale bars (red): 100μm.
